# Protein thiol alterations drive aberrant phase separation in aging

**DOI:** 10.1101/2023.11.07.566021

**Authors:** Thibaut Vignane, Martín Hugo, Christian Hoffmann, Antionia Katsouda, Jovan Petric, Han Wang, Marko Miler, Ferran Comas, Dunja Petrovic, Suyuan Chen, Jan Lj. Miljkovic, Jordan L Morris, Suvagata Roy Chowdhury, Julien Prudent, Natalija Polovic, Michael P. Murphy, Andreas Papapetropoulos, Dragomir Milovanovic, Milos R. Filipovic

## Abstract

Cellular homeostasis relies on precise regulation through chemical processes, such as protein posttranslational modifications (PTM) and physical processes, such as biomolecular condensation. Aging disrupts this balance, increasing susceptibility to diseases and death. However, the mechanisms behind age-related pathogenesis remain elusive. In this study, we dissected various cysteine PTMs and their impact on protein-mediated biomolecular condensation in aging brain. Our findings reveal that aging is associated with significant remodeling of cysteine PTMs, which impacts protein ability to participate in liquid-liquid phase separation (LLPS). Specifically, aging leads to an increase in protein sulfenylation and sulfonylation, which promotes LLPS and through conformational change increases the propensity of proteins to aggregate. Protein persulfidation, a protective thiol modification, prevents this by causing condensate dissolution. We demonstrate that age-induced alterations in cysteine PTMs influence the LLPS properties of synapsin-1 and G3BP2, resulting in disruptions in neurotransmitter release and stress granule formation, respectively. Additionally, our study uncovers that GAPDH is susceptible to LLPS and cysteine sulfonylation exacerbates its transition from condensates to aggregates. Mice deficient in cystathionine gamma-lyase, a pro-longevity gene that regulates intracellular persulfide levels, exhibit a shorter lifespan and spontaneous development of neurofibrillary tangles.

---

Aging could be seen as a progressive decline of fitness caused by the increasing deleteriome, accumulation of “damage” that is both random and deterministic (*1*). Oxidative damage, particularly one caused by reactive oxygen species, is often highlighted as one of the main drivers of aging (*2*). Versatile redox chemistry of the sulfur amino acid cysteine (*3*), makes it one of the main targets for such age-accumulated damage and also an excellent buffer for the first line of defense against it (*4*). However, not all oxidative posttranslational modifications (PTM) of cysteine are considered detrimental. It has been postulated recently that protein persulfidation, an evolutionarily conserved PTM caused by the gasotransmitter hydrogen sulfide (H_2_S), could protect proteins from excessive oxidation and loss of function (fig. S1) (*5*). Intriguingly, cystathionine gamma lyase (CSE, gene name *cth*), a key H_2_S producing enzyme involved in regulation of protein persulfidation (PSSH) levels, has been identified as a pro-longevity gene in multi-tissue RNA-seq analyses across different mammalian species (*6*, *7*). However, due to the often stochastic nature of their formation, cysteine PTMs are often disregarded and not considered as key regulators of cellular function (*8*).

Liquid-liquid phase separation has been shown to be a mechanism through which components of the cytoplasm (proteins and RNAs) can assemble into distinct compartments (condensates) not delimited by a membrane (*9–13*). Condensate assembly is tightly regulated in the intracellular environment, and failure to control condensate properties, formation and dissolution can lead to protein misfolding and aggregation (*14*–*17*). As these are considered to be hallmarks of aging (*18*, *19*), significant efforts have been put into understanding the importance of phase separation in aging (*12*, *20*, *21*) but causal relationships are lacking, largely due to the complexity and the uncertainty of the aging process. One mechanism by which aging could influence phase separation is through chemical modification such as PTMs of proteins.

We hypothesized that age-induced switch from protein persulfidation towards cysteine oxidation to sulfenic (PSOH) and finally sulfonic acid (PSO_3_H) would affect the ability of proteins to phase separate and promote their aggregation, thereby putting more deleterious pressure on the aging cell causing it slip into pathophysiology.

## Aging causes changes in sulfenylome and persulfidome

General approaches to detect thiol oxidation (*22*) fail to distinguish between PSOH and PSSH (*23*), as both of these modifications are oxidative by nature but intrinsically interconnected albeit with distinct biological functions (*3*, *5*, *24*, *25*). To understand if and how aging affects thiol oxidation we applied selective chemoproteomic approaches for global PSOH and PSSH detection (*5*, *26*) (fig. S2) in the frontotemporal region of 10-week, 10-month, and 18-month-old mice. Sulfenylated proteins were found in most of the cellular compartments (Fig. 1A). Given that PSOH is an early step in the thiol oxidation and considered to be a transient modification with multiple fates (fig. S1), it came as a surprise to observe that the abundances of sulfenylated proteins steadily increased throughout the age (Fig. 1B, fig. S3A-D, table S1). To discern age-dependent patterns of change, we grouped proteins into clusters using the fuzzy c-means method. About one third of all endogenously sulfenylated proteins (fig. S3A,B) were found to significantly change their PSOH levels at some point as the mouse aged with about a half of those that significantly changed showing a continuous age-dependent increase (Fig. 1C, fig. S3C), with these proteins involved in critical pathways such as the ones controlling neurogenerative diseases, protein turnover, folding and synaptic transmission being particularly enriched (Fig. 1D, fig. S3E-F). To exclude that this effect originates from the variation in protein expression levels, label free quantitative proteomic analysis of the same samples was performed (fig. S4, table S2). As expected, the aging did affect the brain proteome to a small extent (fig. S4), which agrees with previous reports (*27*), but the differences in protein expression could not be the reason for the significant observed changes in sullfenylation (fig. S3G-I). Instead, age-induced increase of steady-state levels of PSOH (Fig. 1B) indicates that brain cells experience increasing oxidative stress with aging. This is also reflected in the age-induced changes in protein levels of some key antioxidant enzymes (fig. S5).

**Figure 1.**
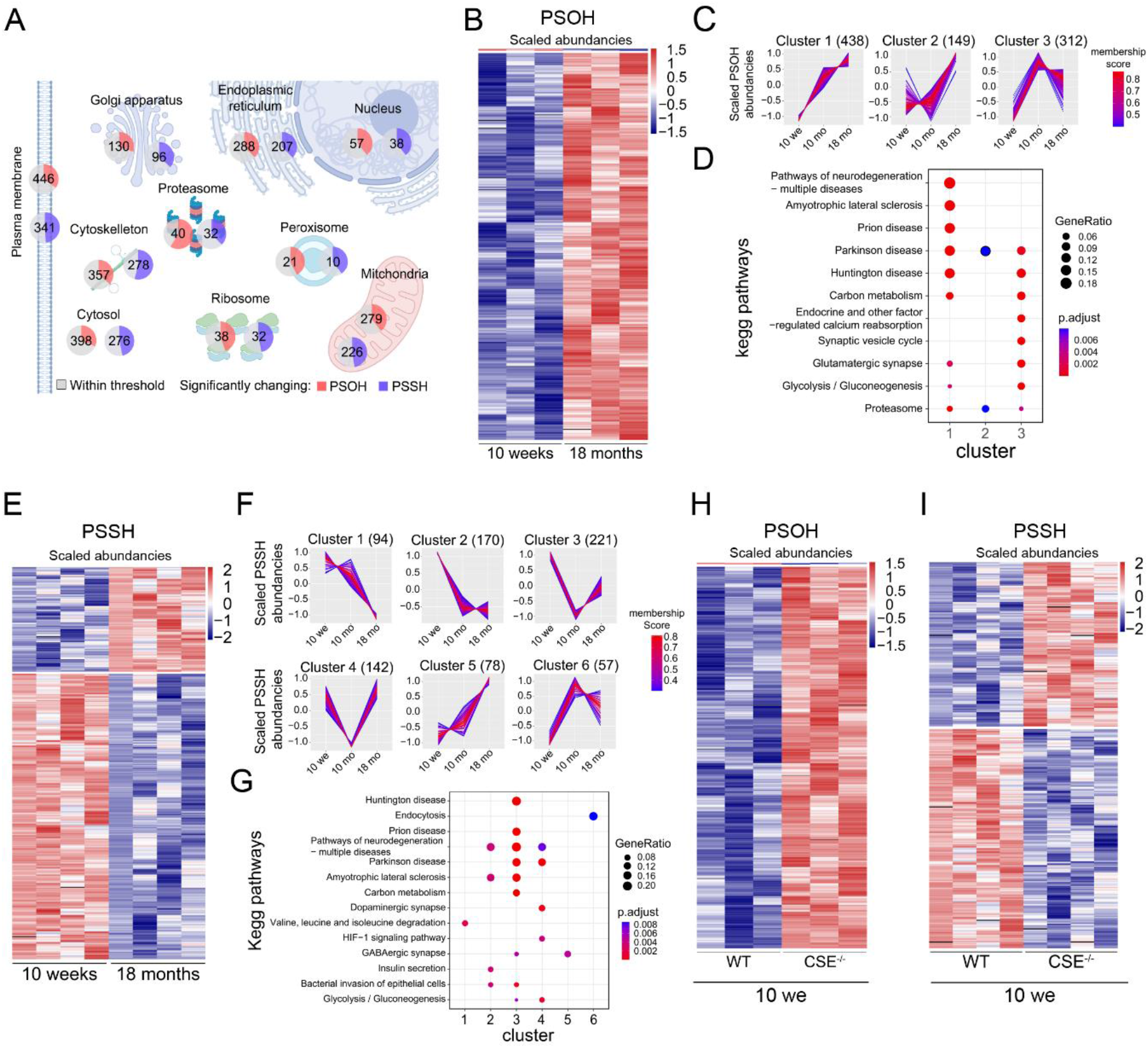
Chemoproteomic analysis of protein sulfenylation and persulfidation reveals substantial alteration of cysteine modifications in aging. (A) PSOH (in red) and PSSH (in blue) changes in subcellular compartments. Colors indicate changes in 18-month-old mice compared to 10-week-old mice; grey represents the proportion of proteins in which the cysteine modification is not changing. The number of proteins for each subcellular compartment is indicated. Protein subcellular localization was determined using DAVID and Gene ontology cellular compartment. (B) Heatmap showing the significant changes of protein sulfenylation in 18-month-old mice compared to 10-week-old mice (Welch’s test, p<0.05). (C) Hierarchical cluster analysis of the 912 proteins with PSOH levels significantly changing at least once in all 3 ages based on trend profile. (D) Kegg pathway enrichment analysis was applied for each cluster shown in (C). Benjamini adjusted p-value was used to filter for significant enrichment. (E) Heatmap showing the significant changes of PSSH in 18-month-old mice compared to 10-week-old mice (Welch’s test, p<0.05). (F) Hierarchical clustering of the 901 proteins for which persulfidation was found to be changing at least once in all 3 ages, into 6 different groups based on the trend profile. (G) Corresponding Kegg pathway analysis for the 6 clusters. Benjamini adjusted p-value was used to filter for significant enrichment. (H-I) Heatmaps of the significant changes (Welch’s test, p-value<0.05) (H) in sulfenylation and (I) persulfidation found in the CSE^-/-^ 10-week-old mice compared to the wild type (WT) 10-week-old mice.

In contrast, protein persulfidation decreased with age (Fig. 1E, fig. S6A-C, table S3). We identified almost 2000 endogenously persulfidated proteins, which is ∼1000 proteins more than were reported to be persulfidated to date using different labeling methods (fig. S6D) (*28–30*). Similar to sulfenylation, persulfidated proteins were found to be present in most of the cellular compartments (Fig. 1A). About a half of them showed significant change at least at one point of age (Fig. 1E,F), with 4 out of 6 clusters exhibiting age-dependent decline (∼80%) and exhibiting enrichment for different metabolic pathways, neurodegeneration, neuronal structure organization and development, etc (Fig. 1G, fig. S6E,F. Total proteome alterations did not significantly affect the observed age-related decline in PSSH (fig. S6G-I). Changes in protein persulfidation correlated well with the decline in protein levels of all three sulfide-producing enzymes, while no change was observed in expression levels of sulfide-oxidizing enzymes (fig. S7).

To test the importance of hydrogen sulfide (H_2_S)-induced protein persulfidation in protection against thiol oxidation, we compared PSOH and PSSH levels in wild type *vs* CSE knockout mice (Fig. 1H-I, fig. S8, table S4). CSE is reported to be a key enzyme involved in regulation of H_2_S levels; its decline is associated with Alzheimer’s disease and spinocerebellar ataxia (*31–33*) and mice lacking it exhibit a Huntington disease-like phenotype (*34*). Indeed, much higher PSOH levels were observed in 10-week-old CSE^-/-^ mice compared to wild type animals of the same age (Fig. 1H, fig. S8A). Moreover, ∼200 proteins showed higher PSOH levels in 10-week-old CSE^-/-^ brains than in 18-month-old wild type brain samples, suggesting that the absence of CSE results in increased steady-state levels of oxidative damage (table S4). Consistent with this protective hypothesis (fig. S1), PSSH levels were reduced in CSE^-/-^ mice (Fig 1I, fig. S8B).

Although there are no established methods to selectively label and enrich protein sulfonylation, we analyzed the proteome data obtained by lysing the tissues with 4-chloro-7-nitrobenzofurazan, which would block all thiols including sulfenic acids (fig. S2) and therefore prevent unwanted cysteine hyperoxidation during sample preparation. Out of 645 detected sulfonylated peptides, 382 were absent in all wild type 10-week-old mice, suggesting that both aging and absence of CSE are characterized with increase of protein sulfonylation (table S5). Molecular dynamics simulation (*35*) of the effect that sulfonylation could have on structures of several targets found to be exclusively present in CSE^-/-^ mice, showed significant conformational changes in the protein structure of affected targets (fig. S9). Finally, immunoblotting for the well-defined sulfenylated proteins, DJ-1 and PRDX, also showed clear differences between aged wild type and CSE^-/-^ mice (fig. S10).

## Cysteine PTMs contribute to the liquid-liquid phase separation of synapsin 1

Having noticed that activity of some of the metabolic enzymes does not seem to be affected by aging, despite the noticeable thiol oxidation change (fig. S11), we next wanted to test the hypothesis that the age-induced thiol oxidation remodeling of proteins could be affecting their mesoscale organization in neurons. Interestingly, we noticed that synapsin 1, the most abundant phosphoprotein in nerve terminals known to phase separate (*36*), showed age-dependent increase in PSOH and decrease in PSSH (Fig. 2A). These changes were already as profound in 10-week-old CSE^-/-^ mice as they were in 18-month-old wild type animals (Fig. 2A). Similar age-induced effects could be seen for other synapsin isoforms (fig. S12A).

**Figure 2.**
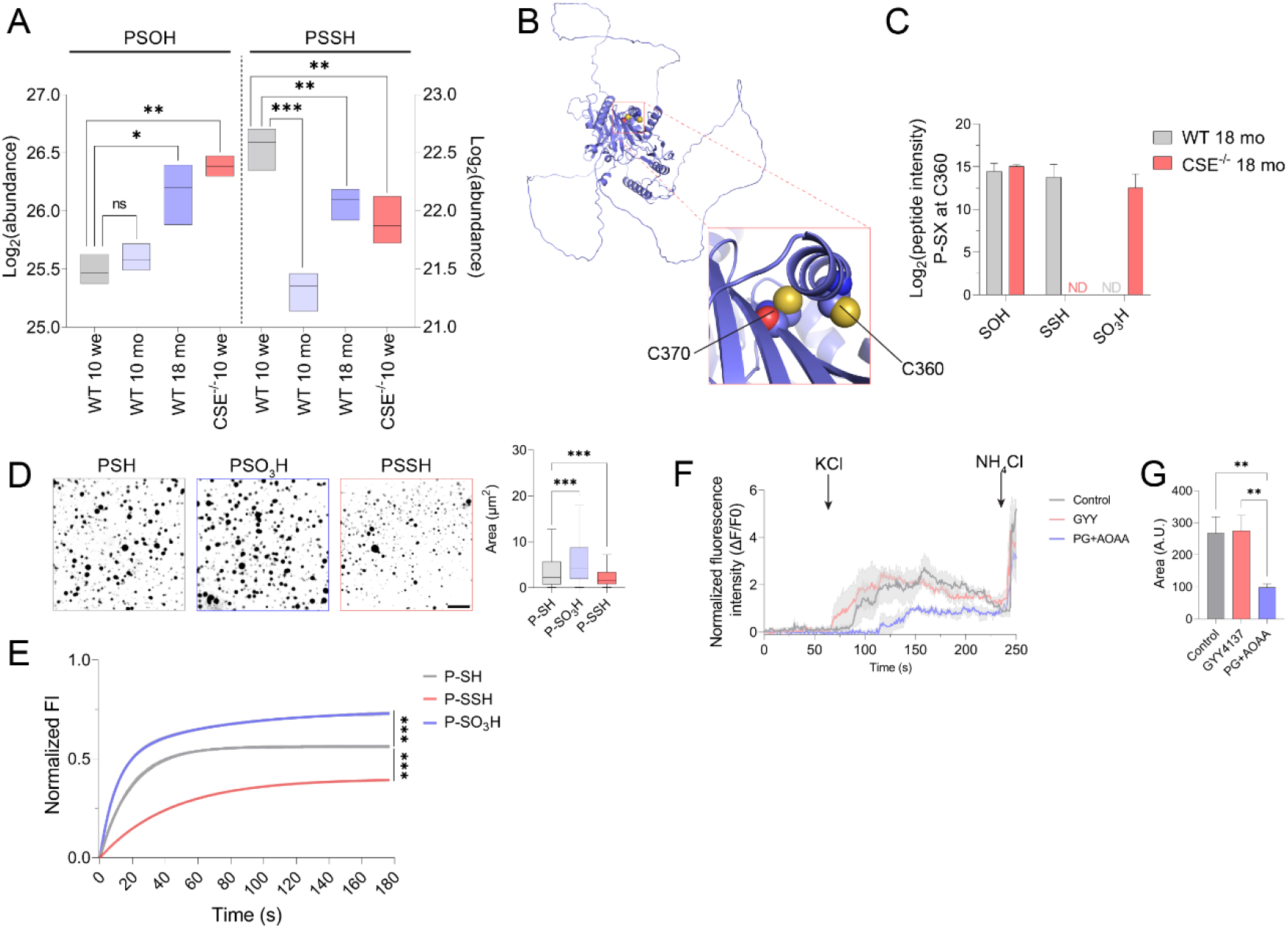
Synapsin 1 liquid-liquid phase separation is controlled by its cysteine’s oxidative status. (**A**) Boxplot of the age-induced sulfenylation (PSOH) and persulfidation (PSSH) of synapsin 1 (n = 4, Welch’s test, p-value: * <0.05, **<0.01, ***<0.001). (**B**) Structure of synapsin 1 highlighting two of its cysteines C360 and C370. (**C**) Changes of sulfenylation (SOH), persulfidation (SSH), and sulfonylation (SO_3_H) of the cysteine 360 but not C370, could be observed in WT and CSE^-/-^ 18 months old mice (n = 3, ND: Non-Determined). (**D**) Representative microscopy images for EGFP-synapsin 1 condensates (10 µM in the presence of 3% PEG3000) formed under different cysteine modifications. The Tukey box and whiskers plots represent the distribution of the droplet area in µm^2^ among the different cysteine modifications (one-way ANOVA, p-value: * <0.05, **<0.01, ***<0.001). n ≥ 4. Scale bar = 20 µm. (**E**) Fluorescence recovery after photobleaching quantification of EGFP-synapsin 1. Means are shown as line and standard error as a shade of grey (n ≥ 3, two-way ANOVA, Sidak posthoc test, p-value: ***<0.001). (**F-G**) The neurotransmitter release from mouse primary neurons without or pre-treated with 100 µM GYY4137 or 1 mM mixture of propargylglycine (PG) and aminooxyacetic acid (AOAA) for 1 h. Synaptic vesicle release was triggered by perfusion of neurons with 92.5 mM KCl; the fluorescence of synaptophysin-pHluorin was used as a readout. Means are shown as lines and standard error of the mean as a shade of grey. The area under the curve was calculated for each measurement (**G**) (n ≥ 3 animals, a few preparations per animal, one-way ANOVA, Tukey’s HSD Test for multiple comparisons p-value: **<0.01).

Synapsin 1 has two cysteines (Fig. 2B) and we observed that C360 could be found in all three forms, PSOH, PSSH and PSO_3_H, the latter being found only in aged CSE^-/-^ mice (Fig. 2C, fig. S12B). C360 is a part of the highly structured, ATP-binding domain of synapsin 1 responsible for its multimerization. Posttranslational modification of a cysteine in that region is not expected to affect phase separation, which has been shown to depend primarily on the intrinsically disordered regions of synapsin 1 (*36*). Despite the fact that PSOH and PSSH are intrinsically unstable (*37*, *38*), we designed the protocol to prepare all these different forms of synapsin 1 (fig. S13A), and compared their ability to phase separate. Indeed, when incubated with crowding agent (3 % PEG 3,000), regardless of cysteine redox form, synapsin 1 readily formed condensates (Fig. 2D, fig. S13B,C). However, synapsin 1 PSO_3_H formed much bigger droplets than fully reduced protein, while droplets formed by synapsin 1 PSSH were significantly smaller (Fig. 2D, fig. S13B). We found that these condensates harbored dynamic behavior and recovered promptly after photobleaching, but they exhibited different kinetics of recovery (Fig. 2E, fig. S13D,E). The percentage and the rate of recovery increased in synapsin 1 PSO_3_H (11.6 s *vs.* 13.0 s; 25 % increase in recovery) while both parameters decreased in synapsin 1 PSSH (29.9 s *vs.* 13.0 s; 30 % decrease in recovery) (Fig. 2E). From the *t*_1/2_, we estimated the apparent diffusion coefficient (*D*app) of the protein molecules to be 8.6, 7.7 and 3.3 x 10^-4^ μm^2^ s^−1^ for the PSH, PSO_3_H and PSSH, respectively. The size and FRAP recovery kinetics of droplets formed from synapsin 1 treated with just H_2_O_2_ (to form PSOH/disulfide mix) or H_2_S (expected not to react at all) was undistinguishable from those formed by fully reduced synapsin 1 (fig. S13C-E). No effect of H_2_O_2_ treatment alone also excludes the possibility that methionine oxidation could be playing a role (*39*). These data clearly indicated that cysteine redox status of synapsin 1 controls its phase separation propensity.

Smaller condensates and slower diffusion suggest that when persulfidated, the ability of synapsin 1 to phase separate is limited. If this were the case, then H_2_S-induced PSSH formation would have profound effects on the dispersion of synaptic vesicles and thus on the neurotransmitter release. Indeed, the exposure of primary neurons to H_2_S resulted in much faster KCl-induced dispersion of synapsin 1 from presynaptic terminal in primary hippocampal neurons (fig. S13F). Consistent with these findings, we also observed that the treatment of primary mouse neurons with the slow releasing H_2_S donor GYY4137 resulted in a faster neurotransmitter release, while the inhibition of CSE not only delayed the release by ∼2-fold but also resulted in overall lower release of synaptic vesicles (Fig. 2F,G, movie S1-3), confirming that the cysteine redox changes affect neurotransmitter release even at the endogenous level. Furthermore, these results provided a mechanistic explanation for the long-suggested role of endogenously produced H_2_S in neurotransmitter release (*40–42*), particularly in the context of those neurodegenerative diseases, which are associated with lower CSE expression levels (*31–34*).

## Cysteine redox status of G3BP2 affects stress granule formation

In addition to molecular crowding-induced phase separation, proteins undergo condensate formation when interacting with RNAs (*43–48*). One such subset of ribonucleoprotein granules is stress granules, initiated by different stressors to cause arrest of translation initiation and protection of RNAs (*17*, *49*). Prompted by the observations that the thiol oxidation status of synapsin 1 affects its ability to phase separate, we compared the datasets obtained for aging PSSH and PSOH with proteomes of stress granules and found that ∼1/3 of all proteins reported to be associated with stress granules have significantly lower PSSH levels and significantly higher PSOH levels (fig. S14A). Among them, we observed that G3BP2 (Ras GTPase-activating protein-binding protein 2), one of the key initiators of stress granule formation (*45*, *46*), has ∼2-fold increase in PSOH and a similar decrease in PSSH levels, caused by aging or absence of CSE (Fig. 3A). G3BP2, the predominant paralogue expressed in brain (*50*), like G3BP1, has one cysteine residue (C73) located within the structured beta-sheet domain (Fig. 3B) that appeared early in evolution (fig. S14B-C).

**Figure 3.**
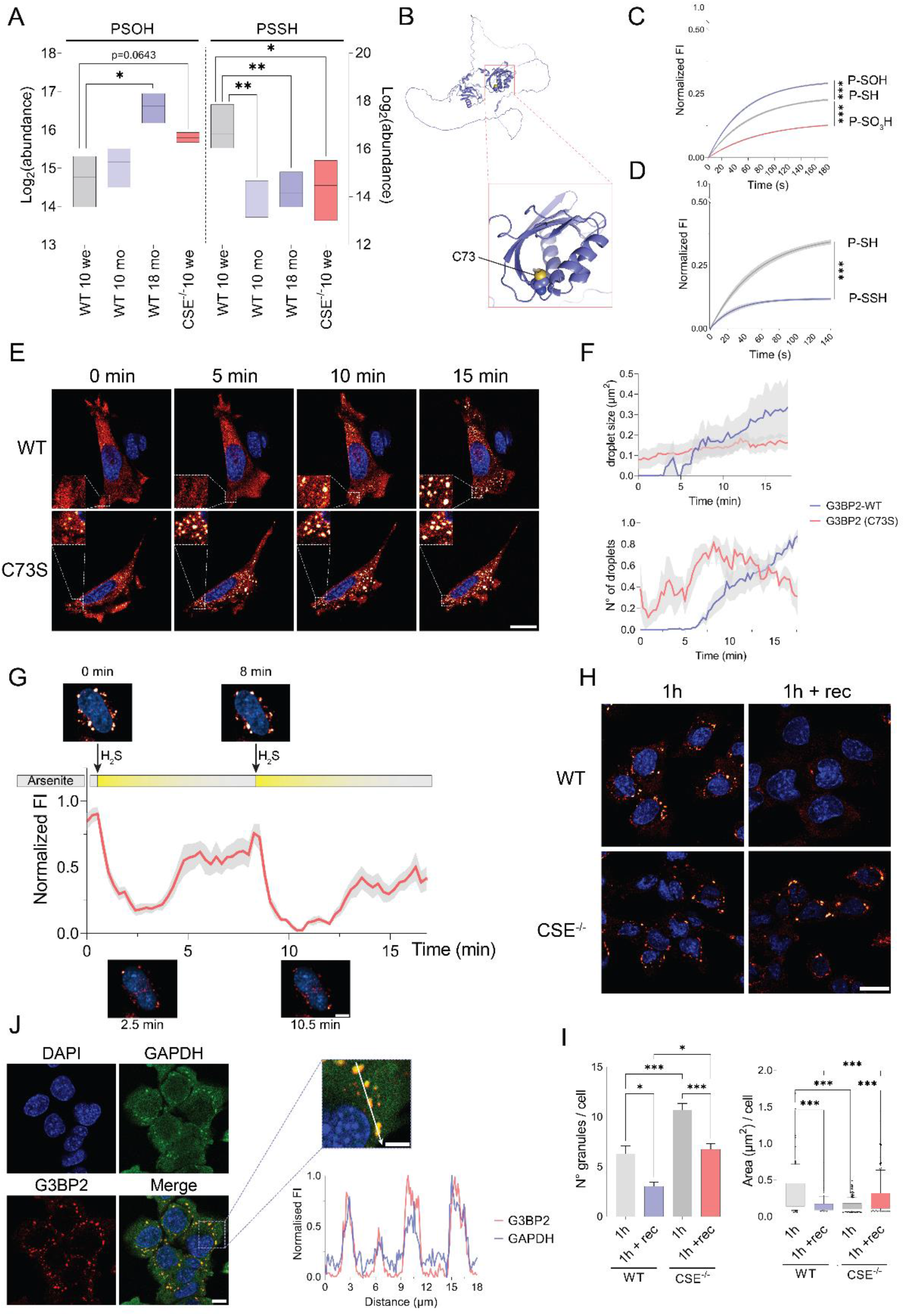
G3BP2 phase separation and associated stress granule formation are controlled by the modifications of its unique cysteine. (**A**) Boxplot of sulfenylation (PSOH) and persulfidation (PSSH). Log2 abundancies of G3BP2 in aging WT mice and CSE^-/-^ mice (n = 4, Welch’s test, p-value: * <0.05, **<0.01). (**B**) G3BP2 alpha fold predicted structure (AF-P97379-F1-model_v4.pdb). Cysteine 73 is unique and present in the NTF2-like domain of the protein. (**C-D**) FRAP quantification of the G3BP2-Cy3 for (**C**) the sulfenylated (PSOH) and sulfonylated (PSO_3_H) G3BP2 and (**D**) persulfidated (PSSH) form of G3BP2. Means are shown as lines and standard error as a shade of grey (n ≥ 3, Two-way ANOVA, Sidak posthoc test, p-value: ***<0.001). (**E**) Arsenite stress (500 µM) induced G3BP2 condensate formation in mouse embryonic fibroblasts (MEF) expressing wild type (WT) or C73S mutant of G3BP2-GFP (representative confocal images of the real-time recording of at least 3 biological replicates). Scale bar = 20 µm. (**F**) Quantification of the size of the WT or C73S G3BP2-GFP condensates (top graph) and the number of condensates over time (bottom graph). Means are shown as lines and standard error as a shade of grey (at least 3 biological replicates). (**G**) Confocal live cell monitoring of G3BP2-GFP condensate dissolution by addition of H_2_S to the mouse embryonic fibroblasts. Na_2_S (100 µM) was injected 15 min after arsenite (500 µM) was added to the cells to initiate stress granule formation. Fluorescence intensity over time was measured for each single condensate (4 biological replicates). Scale bar = 5 µm. (**H**) Representative confocal images of G3BP2-stress granules induced by 1-hour arsenite followed or not by 1h recovery (1h + rec) in WT and CSE^-/-^ MEFs. Images were obtained by immunofluorescence using anti-G3BP2-antibody. n = 3, several images per replicate. Scale bar = 20 µm. (**I**) Quantification of the (**H**) showing the number of stress granules and size of the granules per cell in WT and CSE^-/-^ MEF. Mean and standard error were used for the bar graph (n>100, one-way ANOVA and Tukey’s HSD test for multiple comparisons, p-value: * <0.05, **<0.01, ***<0.001). (**J**) Co-localization of G3BP2 (red) condensates and GAPDH (green) condensates in MEFs after 1h hour arsenite treatment (representative confocal images). The intensity profiles for both GAPDH and G3BP2 were measured along the line represented by the white arrow, using ImageJ software. Scale bar = 10 µm; in the zoomed image = 5 µm.

To address the potential effects of thiol oxidation on G3BP2 phase separation, we first tested PSOH and PSO_3_H forms of the protein. Upon addition of RNA, droplet formation was observed in both cases, with G3BP2 PSO_3_H forming significantly smaller condensates (fig. S15A) that showed slower FRAP recovery and lower % recovery (t_1/2_ = 55.0 s *vs.* 44.5 s, 42 % decrease; Fig. 3C, fig. S15B). Interestingly, granules formed by G3BP2 PSOH showed faster kinetics and % of FRAP recovery (37.6 s *vs*. 44.5 s, 25 % increase; Fig 3C, fig. S15B), suggesting that sulfenylation could increase the diffusion rate of G3BP2. Since G3BP2 contains only one cysteine, we applied a more preparative approach for PSSH generation (fig. S15C). When compared to fully reduced protein, G3BP2 PSSH showed smaller condensates, decrease in % recovery but faster kinetics of FRAP recovery (68 % decrease, 19.6 s) (Fig. 3D, fig. S15D,E). Surprisingly, incubation of G3BP2 with thiol blocking reagent iodoacetamide resulted in significantly bigger granules but almost completely blunted their FRAP recovery (fig. S15D,E).

All these findings suggested that cysteine and its PTMs might play a crucial role in altering stress granule formation *in vivo*. To test that, we expressed GFP tagged wild type and C73S mutant G3BP2 in mouse embryonic fibroblasts and monitored their response to arsenite-induced stress. Cells expressing C73S mutant G3BP2 clearly showed stress granules even in the absence of the stressor (Fig. 3E, movie S4) and their size increased with time only slightly, while their number per cell doubled upon arsenite treatment (Fig. 3F). Cells expressing wild type G3BP2 showed no stress granules prior to arsenite exposure and showed time-dependent increase in granule average number and size once cells were exposed to arsenite (Fig. 3E-F). We conclude that G3BPs cysteine may have evolved to modulate protein’s ability to phase separate upon stress. As arsenite signals *via* ROS and protein sulfenylation (*51*), and considering that G3BP2 PSOH showed better diffusibility and propensity to form droplets (Fig 3C, fig. S15B), we propose that this modification might be a regulator of G3BP2-initiated stress granule formation. Indeed, we could trap G3BP2 PSOH formation upon arsenite treatment and observe time-dependent rise in PSOH, which was more profound in cells lacking CSE (fig. S15F).

Because H_2_S induced dispersion of synaptic vesicles *in cellulo*, we hypothesized that H_2_S-induced persulfidation of G3BP2 might have similar effect on stress granules. Indeed, when H_2_S was added to the cells stressed with arsenite, immediate dissolution of stress granules could be recorded in real-time (Fig. 3G, movie S5). Due to the constant presence of a stressor, fast H_2_S consumption kinetics by the cells (*52*) and its disappearance in a gas phase (*53*), stress granule would reform in time, but full dissolution could be repeated by addition of another bolus of H_2_S (Fig. 3G, movie S5). The reversible nature of this process suggested that endogenously produced H_2_S might play important role in regulating stress granule formation/duration.

As expected, CSE^-/-^ cells treated with arsenite showed a higher number of stress granules per cell albeit their size was significantly smaller (Fig. 3H,I). More interestingly, while wild-type cells almost fully recovered by dissolving their stress granules after this stressor was removed, the size of stress granules in CSE^-/-^ cells continued to increase. Time-resolved stress granule formation also showed that CSE^-/-^ cells form G3BP2 stress granules faster (fig. S16), in agreement with observation that PSOH increases their formation (Fig. 3C). Similar behavior could be observed in striatal neurons from Huntington’s disease (STHdh^Q111/Q111^) mice (Fig. S17), as they are known to have diminished CSE and PSSH levels (*5*). Treatment of cells with slow releasing H_2_S donor GYY4137 reduced the size of stress granules several-fold in both CSE^-/-^ and (STHdh^Q111/Q111^), confirming the effect of H_2_S-induced cysteine persulfidation in melting of stress granules (fig. S17A-C).

## Cysteine sulfonylation but not persulfidation stimulates GAPDH phase separation

Co-localization of G3BP2 condensates with one of the main glycolytic enzymes, glyceraldehyde-3-phosphate dehydrogenase (GAPDH) (Fig. 3J, fig. S16A, S17A,D), came as a surprise, considering that GAPDH is a highly structured protein (fig. S18), although recent studies suggested that some glycolytic enzymes could form glycolytic “G-bodies” under hypoxia (*54*, *55*). Puzzled by this observation we tested if purified human recombinant GAPDH could undergo phase separation. Indeed, increasing concentrations of PEG resulted in condensates that showed liquid-like behavior and were dynamic in FRAP experiments (fig. S19A-C). More importantly, the interaction with RNAs stimulated the condensate formation even more efficiently (Fig. 4A).

**Figure 4.**
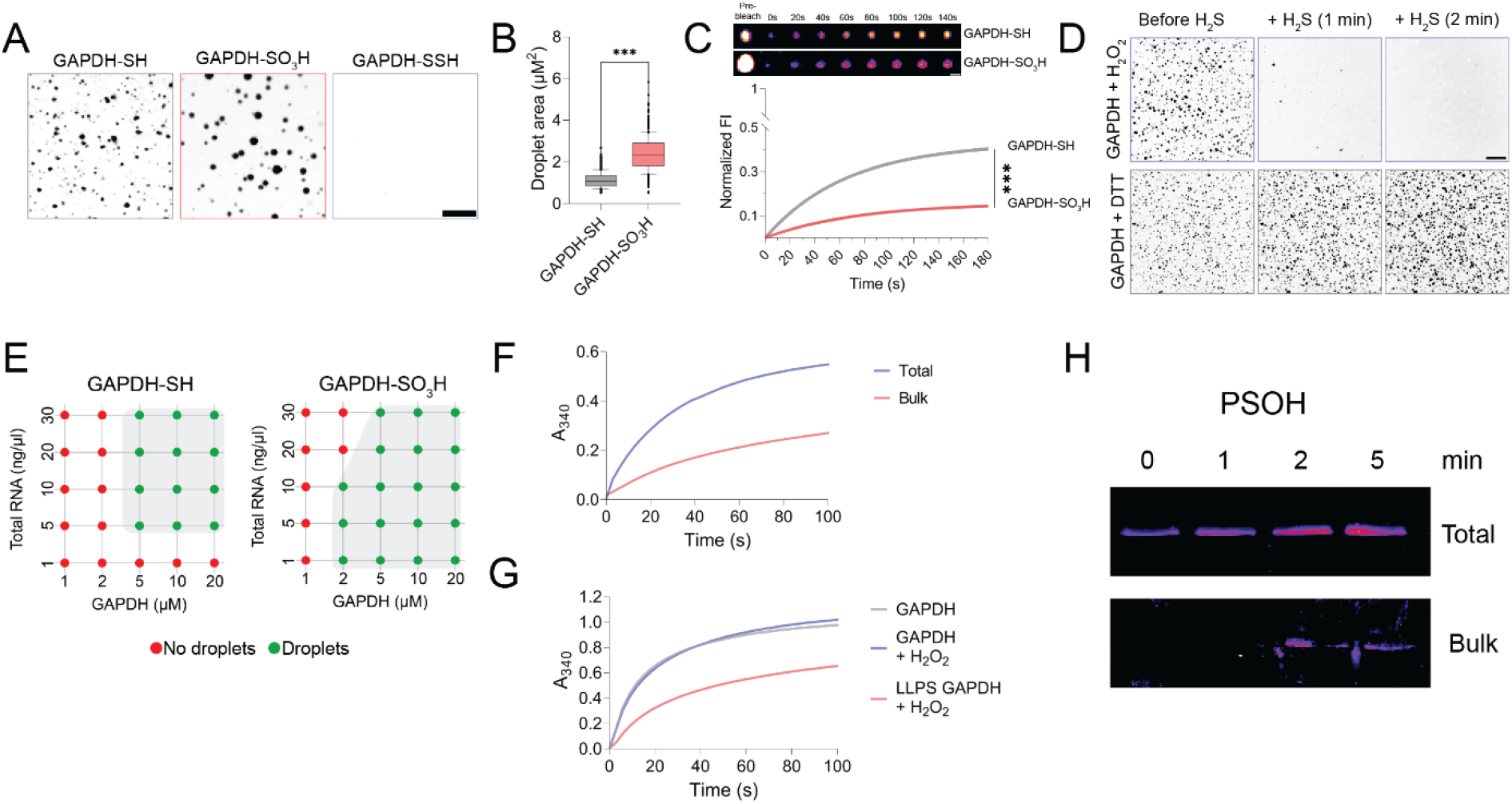
Liquid-liquid phase separation of GAPDH is triggered by RNA and controlled by cysteine modifications. (**A**) Representative images of GAPDH-Cy3 condensate formation from fully reduced (GAPDH-SH), sulfonylated (GAPDH-SO_3_H) and persulfidated (GAPDH-SSH) GAPDH (n = 5). Scale bar = 10 µm. (**B**) Quantification of the GAPDH condensates area (in µm^2^) in the reduced and sulfonylated form (n = 4). (**C**) FRAP quantification of the GAPDH-Cy3 and representative microscopy images. Means are shown as lines and standard deviation as a shade of grey (Two-way ANOVA, Sidak posthoc test, p-value: ***<0.001). Scale bar = 2 µm. (**D**) Representative microscopy images of GAPDH condensate dissolution triggered by persulfidation. 20 µM of GAPDH was incubated for 5 min with RNA in the presence of 100 µM H_2_O_2_ or 100 µM DTT. Droplets were recorded before and 1 and 2 min after the addition of H_2_S (200 µM). Scale bar = 40 µm. (**E**) Summary of the phase separation behavior of the reduced and sulfonylated GAPDH with increased concentration of total RNA. Corresponding images are shown in fig. S21. (**F**) GAPDH activity in total and bulk fractions (fig. S22A) was measured through NADH production at 340 nm. n = 3. (**G**) Kinetics of soluble and phase separated (LLPS) GAPDH (10 µM) inactivation by H_2_O_2_ (100 µM). n = 3. (**H**) BTD trap assay for PSOH formation in the reaction of H_2_O_2_ with GAPDH in total and bulk solution. n = 2.

GAPDH active site cysteine reacts ∼2 orders of magnitude faster with H_2_O_2_ than other protein thiols (*3*), eventually resulting in cysteine sulfonylation and inhibition of enzyme activity. Using the protocol to efficiently generate GAPDH PSO_3_H (fig. S19D,E), we tested how cysteine oxidation of GAPDH affects its phase separation. Our data suggest that oxidation of GAPDH results in much bigger condensate formation when compared to the fully reduced enzyme (Fig. 4A,B, fig. S19F,G), suggesting that not only sulfonylation but also sulfenylation and/or disulfide formation enhances phase separation. While the condensates of fully reduced GAPDH showed FRAP recovery, this recovery was significantly hampered in droplets formed by oxidized GAPDH (Fig. 4C, fig. S20). In contrast to GAPDH PSO_3_H, GAPDH PSSH did not form any condensates in the tested range of protein/RNA concentrations (Fig. 4A). This was confirmed by testing if H_2_S-induced persulfidation could dissolve already formed GAPDH condensates. To test this, we incubated GAPDH with RNA and H_2_O_2_ or DTT for 5 min and recorded condensate formation. Addition of H_2_S completely dissolved condensates within 2 min under the conditions of PSSH formation (H_2_O_2_ + H_2_S), while they continued to grow in GAPDH treated with DTT and H_2_S, as no chemical reaction could occur on cysteine residues (Fig. 4D). Conversely, sulfonylation of GAPDH significantly enforced phase separation, with as low as 16/1 protein/RNA ratio being sufficient to efficiently result in droplet formation (Fig. 4E, fig. S21).

Next, we tested if and how phase separation affects GAPDH enzymatic activity. After allowing 45 min to the mixture of GAPDH and RNA to phase separate, we centrifuged the droplets to generate bulk solution and compared its GAPDH activity to original reaction mixture (total) (fig. S22A). Significant difference in reaction rates, monitored as NADH formation, was observed, with total solution resulting in faster and more efficient activity (Fig. 4F). The difference between total and bulk solution is caused by condensates which constitute as little as 1.5-2% of the reaction volume. The observed activation of GAPDH suggests that, once in the phase, enzyme probably occupies different conformation which makes it more active but could also make it more prone to oxidation.

One of the major questions in redox biology is how is it possible to have so many oxidized proteins when kinetics of the reaction between thiols and H_2_O_2_ is so slow that it cannot outcompete H_2_O_2_ detoxifying enzymes (*56*). To address the possibility that phase separation might be the reason for such efficient thiol oxidation, we investigated the reaction of soluble or phase separated GAPDH with H_2_O_2_, at a concentration that does not inhibit enzyme activity significantly over this time frame. As hypothesized, the activity of GAPDH that was mixed with RNAs so as to cause phase separation was significantly inhibited by H_2_O_2_ (fig. 4G). In the presence of physiological concentration of bicarbonate, which is known to enhance thiol oxidation by H_2_O_2_ by forming HCO_4_^.-^ (*57*), this effect was even more profound (fig. S22B,C). The same inhibition rate in both total and bulk solution suggests that the inhibition was almost completely caused by the oxidation of GAPDH in the phase. Indeed, trapping of PSOH showed that when phase separated, GAPDH becomes more prone to cysteine oxidation even in the absence of H_2_O_2_ (Fig. 4H) confirming that phase separation changes accessibility and reactivity of GAPDH cysteine, making it more prone for oxidation.

## Sulfonylation leads to protein aggregation

Mouse GAPDH contains 3 cysteines (C150, C154 and C245) that are reported to be prone to oxidation, with the oxidation of catalytic C154 being considered as responsible for the loss of enzyme activity (Fig. 5A). Indeed, when analyzing the cysteine oxidation status of mouse brain samples, we observed that all three cysteines are found to be sulfonylated, but it was C245 that showed the strongest age-induced increase which was even more pronounced in CSE^-/-^ mice (Fig 5B, fig. S23). C245 is located in beta-sheet domain. Molecular dynamic modeling of sulfonylated C245 showed profound changes in the GAPDH structure, particularly in beta-sheet domain which seems to be more flattened (Fig. 5C). This observation was further complemented by FT-IR analysis of the structure, where significantly higher percentage of beta-sheets could be detected in GAPDH PSO_3_H, especially in the spectral region (1616–1625 cm^−1^) attributed to aggregation prone inter/intramolecular beta-sheets (Fig. 5D, fig. S24).

**Figure 5.**
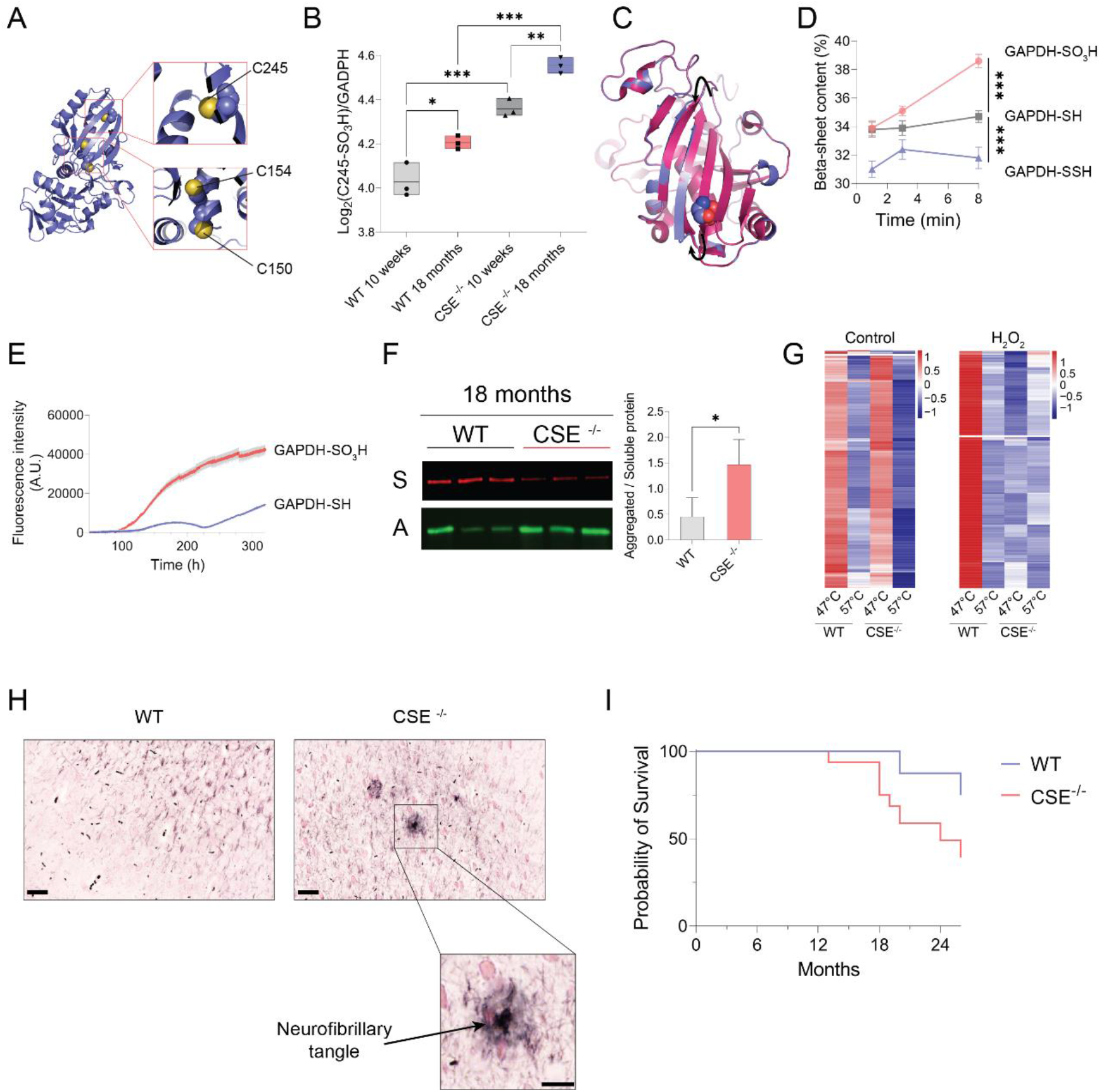
Cysteine sulfonylation drives aggregation through phase transition with LLPS. (**A**) Structure of human GAPDH (PDB:1U8F) highlighting the two cysteines in the active site (C150 and C154) and the cysteine 245 on the surface for the protein. (**B**) Log2 abundancies of the cysteine 245 PSO_3_H peptide normalized to the protein level in aging WT and CSE^-/-^ mouse brain (Two-way ANOVA and Tukey’s HSD test for multiple comparisons, p-value: * <0.05, **<0.01, ***<0.001). (**C**) Molecular dynamics simulation of GADPH structural changes upon C245 sulfonylation. The pink color shows the original crystal structure and the blue color is the predicted model. Black arrows show the flattening of the beta sheets. (**D**) Fourier transform infrared (FT-IR) analysis of the beta-sheet content (%) in the sulfonylated, reduced, and persulfidated GAPDH over time. (**E**) Proteostat aggregation assay on 20 µM reduced and over-oxidized GAPDH in the presence of 30ng/µl total RNA. Using these concentrations GAPDH undergoes phase separation triggering the aggregation of the protein. (**F**) GAPDH aggregation in 18-month-old WT and CSE^-/-^ brain samples. Soluble and aggregated fraction were obtained by ultracentrifugation and GAPDH visualized by western blot (A: aggregated fraction, S: soluble fraction). The ratio of aggregated over soluble fraction was quantified and shown as a bar graph with standard deviation (n = 3 animals, t-test, p-value: *< 0.05). (**G**) Thermal proteome stability assay between WT and CSE^-/-^ MEF cells. (**H**) Gallyas silver staining of WT and CSE^-/-^ 18-month-old mouse brain sections for the neurofibrillary tangles. Global stained sections are shown in fig. S26. n = 4 animals. Scale bar = 40 µm; zoomed image 20 µm. (**I**) Survival rates of CSE^-/-^ vs. WT mice. n = 16 per group.

GAPDH aggregates have been reported in brains from Alzheimer’s disease patients (*58*, *59*) and having beta-sheet domain more exposed could make GAPDH prone to aggregation. We tested if natively folded GAPDH would aggregate but could not observe any aggregation in neither reduced or sulfonylated GAPDH (fig. S25). However, addition of RNA to induce phase separation caused clear protein aggregation of hyperoxidized GAPDH and somewhat slower and less efficient aggregation in reduced GAPDH (Fig. 5E), which could be explained by the observed spontaneous thiol oxidation for the proteins which are in phase (Fig. 4H). Age-induced decline in protein persulfidation and increase of sulfonylation, particularly in CSE^-/-^ mice should result in increased GAPDH aggregation in brain. To test it we separated soluble and aggregated fraction of brain lysates from old WT and CSE^-/-^ mice and observed that indeed there was a ∼3-fold increase in GAPDH in aggregated fraction in mice lacking CSE (Fig. 5F)

As observed in the molecular dynamics modeling for some of the identified sulfonylated targets, cysteine is usually located within or at the beginning/end of the beta-sheet and it causes significant conformational change in the protein (fig. S9). If that were generally true, then proteins with hyperoxidized cysteine should be more prone to aggregation, analogous to GAPDH. To test this hypothesis, we treated mouse embryonic fibroblasts of wild type and CSE^-/-^ cells with H_2_O_2_ and performed modified thermal proteome profiling, focusing only on the beginning and the end of the melting curve (47 °C vs 57 °C). As expected, the proteome of the cells having lower H_2_S production to rescue thiol oxidation from hyperoxidation (fig. S1) was significantly affected by H_2_O_2_, resulting in majority of proteins losing their stability and aggregating from the solution even at 47 °C (Fig. 5G, table S6).

That this is a global phenomenon, present at the organismal level, was assessed by comparing the brain sections of 18-month-old mice from CSE^-/-^ and wild type. As mice do not spontaneously develop neurodegenerative diseases, complete absence of Gallayas silver staining for neurofibrillary tangles was observed in wild type mice, while CSE^-/-^ mice exhibited numerous lesions throughout the whole brain section not only limited to the frontotemporal region, but including cerebellum and hippocampus (Fig. 5H, fig. S26). Finally, having observed significant death rate even before CSE^-/-^ mice reached 18 months, we compared the lifespans of those animals and found that CSE^-/-^ lifespan is shorter than wild type control (Fig. 5I). These observations suggest that aging, coupled with the additional reduction of H_2_S formation, contributes to aberrant phase separation and results in increased protein aggregation and a shorter lifespan.

## Discussion

The emergence of life is inherently intertwined with the presence of hydrogen sulfide (H_2_S), which, alongside cyanide, played a fundamental role in the genesis of biomolecules (*60*). Cysteine was then used to catalyze first peptide synthesis (*61*), and processes like liquid-liquid phase separation and biocondensate formation have indisputably guided evolution towards the life forms we recognize today (*62–64*). These basic chemical and biophysical processes remained imprinted in cellular function, maintaining its homeostasis. This work now identifies the transformative potential of cysteine posttranslational modifications on protein phase separate, highlighting the protective and regulatory impact of protein persulfidation, modification formed by H_2_S, on differently formed biomolecular condensates. Notably, protein persulfidation drives the dispersion of phase-separated condensates. In contrast, cysteine sulfonylation appears to drive proteins towards enhanced phase separation, ultimately resulting in protein aggregation. Moreover, our study underscores how phase separation governs the genesis of thiol modifications by augmenting thiol’s reactivity and predisposing them to oxidation, which might be the key mechanism for signaling effects assigned to ROS (*3*, *56*, *65*). The compartmentalization of ROS action in the cytoplasm within signaling complexes that separate out in solution establishes a new paradigm for ROS signaling. ROS could modify and damage the proteins even if its levels do not go up; they simply have to react with proteins in biomolecular condensates. Oxidized proteins will then have tendency to stay longer within condensate and even aggregate forming a vicious cycle feedback loop. This is further emphasized by recent reports that interface of biomolecular condensates serves as source of ROS (*66*).

The role of ROS in aging has stirred both acclaim and controversy (*1*, *67*, *68*). While pivotal in early developmental stages, where it could predispose the organism for a longer life (*69*, *70*), the growing body of evidence confirms its detrimental roles in aging organism (*71*). Our research demonstrates that steady state levels of protein sulfenylation, the first thiol oxidation step in the reaction with ROS, increases continuously with aging leading eventually to cysteine sulfonylation and that this process is, in large, prevented by production of H_2_S. The demonstrated effects of aging-controlled increase of cysteine sulfonylation and decrease of persulfidation on the phase separation bear profound implications for aging cells and organisms. In the context of the brain, this could manifest as compromised synaptic release due to synapsin 1, increased and unresolved formation of stress granules due to G3BP2 and finally accumulation and aggregation of proteins, such as GAPDH, all of which are observed in different aging-associated neurodegenerative disease. Indeed, CSE is found to be downregulated in several neurodegenerative diseases (*32–34*) and CSE^-/-^ mice are already reported to show some phenotypic traits of Huntington’s disease (*34*). We now report multiple neurofibrillary tangles in the brain of these mice, as well as their shorter life. The emergence of H_2_S-releasing pharmaceuticals offers a potential avenue for ameliorating the age-triggered thiol oxidation-mediated effects on phase separation, holding promise for the treatment of neurodegenerative disorders. This is best exemplified by recent findings showing that CSE is one of the pro-longevity genes commonly found in different lifespan extension interventions (*6*, *7*).

## METHODS

### Mice

Wild-type and CSE KO animals (*72*) were bred/housed in ventilated cages, under specific pathogen-free, temperature controlled (22 °C) and 12 h light/dark cycle conditions in full compliance with the guidelines of the Federation of Laboratory Animal Science Association recommendations in the Laboratory Animal Unit of Biomedical Research Foundation of the Academy of Athens (BRFAA) and allowed unrestricted access to chow diet and water. All studies were performed on male mice at the indicated age.

### Cell culture

CSE^−/−^ mouse embryonic fibroblasts (MEF) are kind gift from Professor Bindu Paul (Johns Hopkins University) and were generated from CSE^+/+^ and CSE^−/−^ mice and immortalized using SV40T antigen (*73*). MEF cells were cultured in DMEM, supplemented with 2 mM L-glutamine, 1 % penicillin-streptomycin, and 10 % fetal bovine serum at 37 °C and 5 % CO2. The striatal progenitor cell line STHdh^Q7/Q7^, expressing wild-type huntingtin, and STHdh^Q111/Q111^, expressing mutant huntingtin harbouring 111 glutamine repeats (referred to as Q7 and Q111 cells, respectively) were maintained in DMEM (low glucose, no pyruvate) supplemented with 2 mM L-glutamine, 1 % penicillin-streptomycin, and 10 % fetal bovine serum at 33 °C and 5 % CO2.

### Persulfidation enrichment for proteomic analysis

The brain frontal cortexes were lysed in cold HEN Buffer (50 mM HEPES, 1 mM EDTA, 0.1 mM neocuproine, 1 % IGEPAL, 2 % SDS, pH 7.4) supplemented with 5 mM 4-Chloro-7-nitrobenzofurazan. Frozen tissue was disrupted in lysis buffer using the TissueLyser II (Qiagen) and 5 mm stainless steel beads. Persulfidation enrichment was performed using previously described method (5). Enriched proteins were eluted by incubating beads with 2.25 M ammonia solution for 16 h and collected the following morning for lyophilization. Lyophilized samples were dissolved in digestion buffer (50 mM ammonium bicarbonate [ABC], 1 mM calcium chloride) and digested overnight using trypsin at trypsin: protein ratio 1:20. The desalting was performed on Supel™-Select HLB SPE Tube. Eluates were evaporated under vacuum until dryness. Peptides were dissolved in 0.1 % TFA, and digestion and desalting quality control was performed using an Ultimate 3000 Nano Ultra High-Pressure Chromatography (UPLC) system with a PepSwift Monolithic® Trap 200 µm * 5 mm (Thermo Fisher Scientific).

### Sulfenylation enrichment for proteomic analysis

Brain frontal cortexes were lysed in cold HEN Buffer supplemented with 0.5 mM Benzo[c][1,2]thiazine-based BTD (BTD) synthesized in house following established protocol.4 Lysates were incubated for 2 h at 37 °C followed by two methanol/chloroform precipitations. The sulfenylation enrichment was carried out using a modified method from Fu et al, 2019 (*26*). Shortly, samples were resuspended in HEPES + 2 % SDS and treated with 10 mM DTT for 1 h at 37 °C, followed by 40 mM Iodoacetamide (IAM) for 30 min at RT. Excess DTT and IAM were removed by methanol chloroform precipitation, and proteins were resuspended in 50 mM HEPES pH 7.0, 2 % SDS. 1.5 mg of proteins were incubated for 2 h at 37 °C with 0.2 mM Az-UV-Cleavable-biotin, 1 mM Ascorbic acid, and 0.1 mM Copper-TBTA to perform the click-chemistry reaction. Precipitated proteins were dissolved in Streptavidin Binding buffer (50 mM NaOAc, pH 4.5) and 0.2 % SDS. Protein concentration was performed, and 0.9 mg of proteins were incubated with Pierce™ High Capacity Streptavidin Agarose beads. After 3 h of mixing at RT, beads were loaded onto columns and washed with Streptavidin washing buffer (50 mM NaOAc and 2 M NaCl, pH 4.5). Sulfenylated proteins were then released by incubating beads for 3 h under UV light (365 nm) in 25 mM ammonium bicarbonate buffer. Supernatants were collected and lyophilized. Samples were then digested and desalted as detailed above. Peptides were dissolved in 0.1 % TFA, and digestion quality control was performed as described.

### LC/MS analysis

Peptides were analyzed by high-resolution LC-MS/MS using an Ultimate 3000 Nano Ultra High-Pressure Chromatography (UPLC) system (Thermo Fisher Scientific) coupled with an Orbitrap Eclipse™ Tribrid™ Mass Spectrometer via an EASY-spray (Thermo Fisher Scientific). Peptide separation was carried out with an Acclaim™ PepMap™ 100 C18 column (Thermo Fisher Scientific) using a 65 min linear gradient from 3 to 35 % of B (84 % Acetonitrile, 0.1 % Formic Acid) at a flow rate of 250 nL/min. The Orbitrap Eclipse™ was operated in a DDA mode, and MS1 survey scans were acquired from m/z 300 to 1,500 at a resolution of 120,000 using the Orbitrap mode. MS2 scans were carried out for 3 seconds with high-energy collision-induced dissociation (HCD) at 32 % using the Normal speed IonTrap mode.

Data were evaluated with PEAKS ONLINE software using 15 ppm for precursor mass tolerance, 0.5 Da for fragment mass tolerance, specific tryptic digest, and a maximum of 3 missed cleavages. For persulfidation detection, NBF (+163.00961594 Da) on C, K, R, DCP (+168.0786442 Da) on C, N-term acetylation (+42.010565 Da), and methionine oxidation (+15.994915 Da) were added as variable modifications. For sulfenylation detection, BTD–UV-cleavable probe (cleaved, +418.1311 Da), carbamidomethylation (+57.021464 Da) on C, N-term acetylation (+42.010565 Da), and methionine oxidation (+15.994915 Da) were added as variable modifications, PSM and proteins were filtered at FDR 1 %. Data were normalized using eigenMS on R studio (*74*).

### Total proteome analysis

Brain frontal cortexes were disrupted in lysis buffer (50 mM TRIS, 150 mM NaCl, 2% SDS, 1% IGEPAL, pH 7.8) using the Tissuelyser II and twenty 1.4 mm ceramic beads (Qiagen, #13113). Lysates were incubated with Benzonase following manufacturer recommendation for 30 min and precipitated. Samples were then resuspended in 50 mM HEPES, 2 % SDS, and incubated with 10 mM DTT at RT followed by incubation with 50 mM IAM for 1h at 37°C. Precipitation was used to stop the reaction and concentrate proteins. For tryptic digestion, samples were resuspended in 6 M guanidine hydrochloride.250 µg of protein were digested overnight at 37 °C using trypsin with a 1:20 trypsin to sample ratio (Promega, #V5117). The next day, peptides were desalted evaporated under vacuum until dryness as described above.

Peptides were dissolved in 0.1 % TFA, and digestion quality control was performed as described above. Peptides were analyzed by high-resolution LC-MS/MS using an Ultimate 3000 Nano Ultra High-Pressure Chromatography (UPLC) system (Thermo Fisher Scientific) coupled to a timsTOF Pro (Bruker) equipped with a CaptiveSpray source. Peptide separation was carried out with an Acclaim™ PepMap™ 100 C18 column (Thermo Fisher Scientific) using a 65 min linear gradient from 3 to 35 % of B (84 % Acetonitrile, 0.1 % Formic Acid) at a flow rate of 250 nL/min. The timsTOF Pro (Bruker) was operated in PASEF mode using Compass Hystar 5.0.36.0. The TIMS setting was as follows: Mass Range 100 to 1700 m/z, 1/K0 Start 1.6 Vs/cm2 End 1.6 Vs/cm2, Ramp time 100 ms, Lock Duty Cycle to 100 %, Capillary Voltage 1500V, Dry Gas 3 l/min, Dry Temp 180°C. PASEF settings: 10 MS/MS scans, charge range 0-5, active exclusion for 0.4 min, Scheduling Target intensity 20000, Intensity threshold 2500, CID collision energy 49 eV. Data were evaluated with PEAKS ONLINE software using 20 ppm for precursor mass tolerance, 0.5 Da for fragment mass tolerance, specific tryptic digest, and a maximum of 3 missed cleavages. Carbamidomethylation (+57.021464 Da) on C, N-term acetylation (+42.010565 Da), and methionine oxidation (+15.994915 Da) were added as variable modifications, PSM and proteins were filtered at FDR 1 %. Data were normalized based on the total ion count (TIC).

### Direct detection of Sufonylated peptides using LC/MS

50 µg of the NBF-labelled samples dissolved in PBS 2 % SDS (see section “PSSH enrichment for proteomic analysis”) were trypsinized overnight as described above. Samples were subjected to desalting and Pierce® Detergent Removal columns (ThermoFisher, # 87777). Peptides evaporated under vacuum until dryness as described above.

Peptides were dissolved in 0.1 % TFA, and digestion quality was assessed as previously described. Peptides were analyzed by high-resolution LC-MS/MS using an Ultimate 3000 Nano Ultra High-Pressure Chromatography (UPLC) system (Thermo Fisher Scientific) coupled with an Orbitrap Eclipse™ Tribrid™ Mass Spectrometer via an EASY-spray (Thermo Fisher Scientific). Peptide separation was carried out with an Acclaim™ PepMap™ 100 C18 column using a 160 min linear gradient from 3 to 35 % of B (84 % Acetonitrile, 0.1 % Formic Acid) at a flow rate of 250 nL/min. Parameters were set as described above (see “PSOH enrichment for proteomic analysis” section).

Data were evaluated with PEAKS ONLINE software using 15 ppm for precursor mass tolerance, 0.5 Da for fragment mass tolerance, specific tryptic digest, and a maximum of 3 missed cleavages. NBF (+163.00961594 Da) on C, K, and R, Trioxidation (+47.984744 DA) on C, N-term acetylation (+42.010565 Da), and methionine oxidation (+15.994915 Da) were added as variable modifications, peptide-spectrum match (PSM) and proteins were filtered at FDR 1 %. Data were normalized based on TIC.

### Immunoblotting

Cells or tissues were lysed in HEN buffer supplemented with 20 mM IAM and incubated at 37°C for 2 h to allow complete cysteine blocking. The excess of blocking-reagent was removed by precipitation, as mentioned above. The samples were then resuspended in HEPES + 2 % SDS, and readjusted using DC™ protein assay. One equivalent of Laemmli (4X) buffer supplemented with 10 % β-mercaptoethanol was added to three sample equivalents for SDS-PAGE and boiled at 95 °C for 5 min protected from light. Equal amounts of proteins were resolved by SDS-page using standard protocol. Samples were then immunoblotted with the indicated antibodies and dilutions; (anti-Prdx-SO3 [Abcam, #ab16830], 1:4000; anti-oxDJ-1 [Sigma-Aldrich, #MABN1773], 1:2000; anti-GAPDH-SO3 [Abfrontier, #LF-PA0006], 1:2000; anti-GAPDH-AF488 [Sigma-Aldrich, #MAB374-AF488],1:20000; anti-β-Tubulin [Sigma-Aldrich, #T0198], 1:5000; anti-Rabbit-AF680 [Abcam, #ab175773], 1:20000; anti-Mouse-AF680 [Abcam, #ab175775], 1:20000).

### LDH and PKM activity assays

For all activity assays, the frozen tissues were disrupted in mild lysis buffer (25 mM HEPES, 250 mM sucrose, 20 mM MgCl2, 1 mM EDTA, pH 7.4) using the TissueLyser II and 5 mm Stainless steel beads. Samples were then centrifuged at 14,000 x g for 15 min at 4°C, and supernatant was collected for further application. Protein concentration was performed, samples were readjusted to equal amounts and split for different activity assays. LDH (Sigma, #MAK066) and PKM (Sigma, #MAK072) activity assays were carried out following the manufacturer’s protocol.

### Direct detection of cysteine PTMs of synapsin 1 and GAPDH by LC-MS/MS

Brain tissues were lysed using the same protocol as for immunoblotting. A modified in-gel digestion protocol from (77) was used to identify GAPDH and synapsin 1 cysteine peptides. An equal amount of protein was loaded and separated by SDS page. Gels were then stained for 2 min with a Coomassie Brilliant Blue solution (1.2 mM Coomassie Brillant Blue, 10 % acetic acid, 40 % methanol in distilled water) and destained overnight in a decoloration solution (25 % Methanol, 8 % acetic acid in distilled water). Once destained, the band corresponding to the size of the protein of interest (-/+ 5kDa) was excised. Coomassie Brilliant Blue was removed by alternating washes with 50 mM ABC and 25 mM ABC, 50 % ACN each for 10 min at 37 °C, 450 rpm. Once free from the stain, gel pieces were dried using a speed vacuum concentrator. Gel pieces were then incubated for 10 min at 4 °C with 50 mM ABC, 1 mM CaCl2, and trypsin at 12.5 ng/µl. The excess solution was removed, and samples were trypsinized for 16 h at 37 °C, 750 rpm. The following day, peptides were extracted by incubating the gel pieces two times in 0.1 % TFA for 15 min at 37°C, 450 rpm. This step was followed by two-times incubation with 0.1 % TFA + 60 % ACN for 15 min at 37°C, 450 rpm. Extracts of different pieces were combined, and peptides were evaporated to dryness.

Peptides were dissolved in 0.1 % TFA, and Digestion was controlled using previously described method. Peptides were analyzed by high-resolution LC-MS/MS using an Ultimate 3000 Nano Ultra High-Pressure Chromatography (UPLC) system (Thermo Fisher Scientific) coupled to a tims_3_ TOF Pro (Bruker) equipped with a CaptiveSpray source. Peptide separation was carried out with an Acclaim™ PepMap™ 100 C18 column (Thermo Fisher Scientific) using a 120 min linear gradient from 3 to 35 % of B (84 % Acetonitrile, 0.1 % Formic Acid) at a flow rate of 250 nL/min. The timsTOF Pro (Bruker) was operated as previously described. Data were evaluated with PEAKS ONLINE software using 20 ppm for precursor mass tolerance, 0.5 Da for fragment mass tolerance, specific tryptic digest, and a maximum of 3 missed cleavages. The entire mouse database was used for the search. SO-IAM (+73.020464 Da), SS-IAM (+89.086464 Da), ’Cysteine oxidation to cysteic acid’ (SO -, +47.984745 Da, Unimod ID: 345), carbamidomethylation (+57.021464 Da) on C, N-term acetylation (+42.010565 Da), and methionine oxidation (+15.994915 Da) were added as variable modifications, PSM and proteins were filtered at FDR 1 %. Data were normalized based on TIC.

### EGFP-synapsin 1 in-vitro phase separation

The recombinant full-length EGFP-Synapsin 1 was expressed in a mammalian expression system and purified as previously described (*75*, *76*). Before the experiment, EGFP-synapsin 1 was cleaned up from TCEP using a dialysis column, and the buffer was replaced with the phase separation buffer. Sulfonylated EGFP-synapsin 1 form was obtained by treating SYN1 with 100 µM DTT and 200 µM H_2_O_2_ at 37 °C for 30 min. Persulfidated EGFP-synapsin 1 was formed by treating the protein with 100 µM H_2_O_2_ and 500 µM H_2_S for 5 min at 37 °C before adding PEG. Respective controls were carried out using the same DTT, H_2_O_2_, and H_2_S concentrations. For the FRAP experiment, 3% w/v polyethylene glycol (PEG 3,000) was used to trigger phase separation of 10 µM EGFP-synapsin1. Photobleaching was performed after 5 min incubation with PEG, and recovery was recorded for 10 min.

### Primary neurons experiments

Primary hippocampal neurons were prepared from newborn pups (p0/p1) and the exocytosis assays were performed as described recently.^6^ Neurons were pre-incubated with GYY4137 (100 µM) or propargylglycine (1 mM) and aminooxiacetic acid (1 mM). During imaging, neurons were first perfused with Tyrode solution supplemented with 2.5 mM KCl (low KCl); subsequently, to stimulate exocytosis, neurons were perfused with Tyrode solution supplemented with 92.5 mM KCl (high KCl) for 5 min; finally, neurons were briefly perfused with Tyrode solution containing 50 mM NH4Cl to alkalize the sample and visualize the total fluorescence from synaptophysin-pHlurin.

### G3BP2 in-vitro phase separation

Total RNA from STHdhQ7/Q7 cells was extracted using the RNeasy Midi Kit and following the manufacturer’s protocol. Total RNA was resuspended in RNase-free water, and concentration was assessed using a Nanodrop.

G3BP2 was purchased from Biomol (#349926). First, isolated G3BP2 was labeled with a Cyanine3 NHS ester probe. The protein buffer was briefly exchanged with the labeling buffer (100 mM Sodium Bicarbonate, 50 mM NaCl, pH 8.0) using the Micro Bio-Spin™ 6 Columns following the manufacturer’s protocol. The protein was then incubated for 1 h at RT with Cyanine 3 NHS ester probe at a ratio of 1:3 (protein: Cyanine3). After incubation, the probe excess was removed using the Micro Bio-Spin™ 6 Columns and the buffer was exchanged with phase separation buffer. To minimize the effect of the Cy3-label on phase separation, the labeled proteins were mixed with non-labeled G3BP2 at a 1:9 ratio (labeled: non-labeled). For all experiments, G3BP2 phase separation was triggered with 50 ng/µl of total RNA extract and incubated 7 min prior to FRAP experiments. Recovery was recorded for 10 min.

### G3BP2 WT and C73S mutant plasmid construction

The pDEST-CMV-N-EGFP-wtG3BP2 plasmid was built using the gateway cloning method.(*77*) The human G3BP2 sequence from the pENTR4_G3BP2 plasmid was transferred into the pDEST-CMV-N-EGFP plasmid using the Gateway™ LR Clonase™ following the manufacturer’s instructions. Both plasmids were obtained through Addgene. pENTR4_G3BP2 was a gift from Thomas Tuschl (Addgene plasmid #127105; http://n2t.net/addgene:127105 ; RRID:Addgene_127105). pDEST-CMV-N-EGFP was a gift from Robin Ketteler (Addgene plasmid # 122842; http://n2t.net/addgene:122842; RRID:Addgene_122842) (*78*).

For the C73S mutation, the point mutation was created following established protocol (*79*). Plasmids were first tested for the mutation using PCR primers allowing the elongation of a smaller fragment when the mutation is present (FW: GACTTCAAGGAGGACGG; RV: GTCTCAGCGACGCTG; RV middle primer: ACATGACGAATTTTAGTATGACT). Positive clones were then sent for sequencing for further validation. Both pDEST-CMV-N-EGFP-wtG3BP2 and pDEST-CMV-N-EGFP-C73SG3BP2 plasmids were amplified and extracted using PureLink™ HiPure Plasmid Maxiprep Kit (Invitrogen™, K210007) before use.

### Plasmid transfection and G3BP2 live cell microscopy

WT MEF cells were transfected with pDEST-CMV-N-EGFP-C73SG3BP2 or pDEST-CMV-N-EGFP-wtG3BP2 using a reverse transfection protocol. Briefly, the transfection solution was prepared by mixing 1 part of plasmid DNA (in µg) and 3 parts of TransIT®-2020 Transfection Reagent (Mirius, #MIR5400) in Opti-MEM™ I Reduced Serum Medium (Gibco™, #31985062) and incubated for 30 min. Meanwhile, cells were trypsinized for 3 min, followed by quenching with DMEM supplemented with 10 % fetal bovine serum. After incubation, the transfection solution was added into a glass bottom µ-slide with 8 wells at 50 ng of DNA per well (IBIDI, #80807). 15,000 cells were seeded in each well on top of the transfection reagent and incubated for 24 h. The day after, the media was changed 2 h before starting the experiment to allow cell recovery from transfection stress.

Live cell microscopy was carried out using a Leica SP8 LIGHTNING Confocal Microscope with a 63x water objective with environmental control (37 °C, 5 % CO2) and adaptive focus control. Before recording, cells were treated with 5 µg/mL Hoechst 33342 for 5 min. Media containing Hoechst 33342 was washed 2 times and replaced with FluoroBrite™ DMEM supplemented with 2 mM L-glutamine, 1 % penicillin-streptomycin and 10 % fetal bovine serum.

### G3BP2 live cell microscopy

For the measurements of stress granule formation, cells transformed with WT G3BP2 or (mutant) C73S-G3BP2 were treated with 500 µM sodium metaarsenite (NaAsO_2_), and one image was taken every 3 sec for 20 min. The number and size of stress granules were measured using Fiji.

For the in-cellulo H_2_S dissolution of G3BP2 condensates, cells expressing the WT G3BP2 plasmid were treated for 15 min with 500 µM NaAsO_2_ to allow the formation of G3BP2 condensates. Cells were then recorded every 16 s, and H_2_S was injected at a 100 µM final concentration. After the condensate reformed, a second injection of H_2_S at the same concentration was performed.

### Sulfenylated G3BP2 immuno-pulldown and detection by LC/MS

90 % confluency WT and CSE-/-MEFs 10 mm dish were treated or not with 500 µM NaAsO_2_ for 15 min and lysed in HEN supplemented with 0.5 mM BTD. After 2 h incubation at 37 °C, proteins were precipitated. Samples were redissolved in PBS 0.1 % SDS. An equal amount of proteins were incubated for 1 h with 1 ug of G3BP2 antibody. The SureBeads™ Magnetic Beads attached to protein A were added to the samples and incubated at 4 °C overnight. Samples were incubated one more hour at RT, and beads were washed 3 times with PBS 0.01 % Tween and 4 times with PBS. The beads containing the G3BP2 pulldown were then resuspended in 50 mM ABC and 0.0125 µg/µl trypsin. Samples were trypsinized for 16 h at 37 °C. Supernatants were then collected, and peptides were desalted and dried as previously described. Peptides were dissolved in 0.1 % TFA, and digestion quality control was performed as previously described. Peptides were analyzed by high-resolution LC-MS/MS using an Ultimate 3000 Nano Ultra High-Pressure Chromatography (UPLC) system (Thermo Fisher Scientific) coupled with an Orbitrap Eclipse™ Tribrid™ Mass Spectrometer via an EASY-spray (Thermo Fisher Scientific). Peptide separation was carried out with an Acclaim™ PepMap™ 100 C18 column (Thermo Fisher Scientific) using a 65 min linear gradient from 3 to 35 % of B (84 % Acetonitrile, 0.1 % Formic Acid) at a flow rate of 250 nL/min. The Orbitrap Eclipse™ was operated in a DDA mode, and MS1 survey scans were acquired from m/z 300 to 1,500 at a resolution of 120,000 using the Orbitrap mode. MS2 scans were carried out for 3 seconds with high-energy collision-induced dissociation (HCD) at 32 % using the Normal speed IonTrap mode.

Data were evaluated with PEAKS ONLINE software using 20 ppm for precursor mass tolerance, 0.5 Da for fragment mass tolerance, specific tryptic digest, and a maximum of 3 missed cleavages. The entire mouse database was used for the search. Pure BTD (+245.045 Da) N-term acetylation (+42.010565 Da), and methionine oxidation (+15.994915 Da) were added as variable modifications, PSM, and proteins were filtered at FDR 1 %. Data were normalized based on TIC.

### Immunofluorescence of G3BP2 and GAPDH under NaAsO_2_ treatment

MEF WT and CSE-/- or STHdh cells were split in a µ-Dish 35 mm, high Glass Bottom (IBIDI, #81158). Before fixation, cells were washed two times with warm PBS and fixed with 2 % PFA for 10 min, followed by 10 min with 4 % PFA.

For immunofluorescence, the permeabilization step was performed in PBS 0.1 % TritonX100 for 20 min. Cells were then incubated for 1 h in blocking buffer (1 % BSA, 22.52 mg/ml glycine, 0.1 % Tween in PBS) followed by a 2 h incubation at 37 °C with the respective primary antibody (anti-G3BP2, 1:600 [Proteintech®, 16276-1-AP]; anti-GAPDH-AF488 [Sigma-Aldrich, MAB374-AF488], 1:500). Cells were washed and incubated with secondary antibody (mouse anti-rabbit IgG-CFL 647, [Santa Cruz Biotechnology, INC.; sc-516251]) for 2 h at 37 °C. DAPI (4’,6-Diamidin-2-phenylindol, Dihydrochloride; Thermo Fisher Scientific) staining was performed prior to recording, following the manufacturer protocol. Imaging was performed using Leica SP8 LIGHTNING Confocal Microscope with a 63x oil objective and a resolution of 2048 x 2048. Image analysis was performed using Fiji.

### GAPDH PSH/PSO3H preparation

All GAPDH phase separation experiments were performed using phase separation buffer previously described and microscopy settings as detailed above unless mentioned otherwise. 50 µM of GAPDH was incubated with 500 µM DTT to form GAPDH-SH or 500 µM DTT and 1 mM H_2_O_2_ for increased time to form GAPDH-SO3H. Protein was then cleaned 2-times on Micro Bio-Spin™ P-6 gel Columns (BioRad, #732-6221) where buffer was exchanged with phase separation buffer. To assess sulfonylated protein yield, the protein was run on SDS-PAGE, and immunoblotting was performed using anti-GAPDH-SO3 and GAPDH-AF488. After 30 min treatment, the maximum yield was reached. For further experiments, 30 min incubation was used to form sulfonylated GAPDH.

### GAPDH in-vitro phase separation with PEG

GADPH was labeled with Cyanine 3 NHS ester as previously described. GAPDH phase separation was first tested by incubating 10 µM protein with an increased concentration of PEG 3,000 (1-10 %) for 5 min. For FRAP experiments, 5 % of PEG3000 and 10 µM of GAPDH were used. Photobleaching was performed after 5 min incubation with PEG3000, and recovery was recorded every second for 10 min.

### GAPDH In-vitro phase separation using total RNA extract

Total RNA was extracted as previously described. To minimize the effect of the Cy3 label on phase separation, the labeled protein was mixed with a non-labeled GAPDH at a 1:9 ratio (labeled: non-labeled). Sulfonylated GAPDH was obtained by incubating 10 µM protein with 100 µM DTT and 200 µM H_2_O_2_ at 37 °C for 30 min. Persulfidated GAPDH was obtained by treating the protein with 100 µM H_2_O_2_ and 500 µM H_2_S for 5 min at 37°C before adding the total RNA extract. Respective controls were carried out using the same DTT (reduced GAPDH) or H_2_O_2_ (oxidized GAPDH) concentrations. For the FRAP experiment, 30 ng/µl of total RNA extract was used to trigger phase separation of 10 µM GAPDH. Photobleaching was performed after 5 min of incubation with RNA, and recovery was recorded every second for 10 min.

To assess the dissolution of GAPDH droplets by H_2_S, 20 µM GAPDH was incubated for 5 min with RNA in the presence of 100 µM H_2_O_2_. Before recording, 100 µM DTT was added or not. Droplets were recorded at baseline, 1 min, and 2 min after adding H_2_S.

To build the phase diagram of reduced and sulfonylated GAPDH, both GAPDH forms were freshly prepared as previously described. GAPDH LLPS was then triggered by adding the indicated total RNA concentration. The solution was then transferred into 15 µ-slide Angiogenesis. All pictures were taken after 5 min incubation using a Leica SP8 LIGHTNING Confocal Microscope with a 63x oil objective in bright field mode.

### GAPDH activity in LLPS

For the entire activity assay, total and bulk solutions were prepared in the following way: 10 µM GAPDH was incubated with 30 ng/µl of total RNA in the phase separation buffer. For the total fraction, incubation was performed at RT for 45 min. To prepare the bulk fraction, the protein was incubated for 30 min with RNA and centrifuged for 15 min at 15,000 x g. After centrifugation, equal volumes of total and bulk RNA were taken and incubated with 1 mM NAD+ for 15 min at RT. 15 mM NaAsO_2_ and 250 µM glyceraldehyde-3-phosphate were added to start the reaction. NADH formation was recorded using a spectrophotometer at 340 nm. For GAPDH in-solution activity, the experiment was performed by omitting RNA into the mix. The inhibition of GAPDH activity by H_2_O_2_ was performed by co-incubating total GAPDH LLPS, 100 µM H_2_O_2_, and 1 mM NAD+ for 15 min before recording. The same experiments were carried out in the presence of 2 Mm sodium bicarbonate (NaHCO3).

### BTD-trap assay

Total and bulk fractions of GAPDH were prepared as previously described. Parts of the fractions were then taken and incubated with 1 mM BTD for 2 min. Before adding H_2_O_2_, part of the samples was then withdrawn for the control condition (0 min) and incubated with IAM (20 mM) and SDS (2 %), preventing BTD from reacting further. After that, H_2_O_2_ was added at a final concentration of 10 µM. Part of the sample was taken at 1, 2, and 5 min and directly incubated with catalase, IAM (20 mM), and SDS (2 %) to stop the reaction. Samples were cleaned using Micro Bio-Spin™ 6 Columns before the labeling step. BTD was then labeled with Cy5 using standard click chemistry reaction protocols. Shortly, samples were incubated for 1.5 h at RT in the dark with 150 µM Cyanine5-Azide (Lumiprobe, #D3030), 300 µM Copper (II)-TBTA complex (Lumiprobe, 21050), and 1 mM L-ascorbic acid (Sigma-Aldrich, #A92902). 3 sample parts were mixed with one part of Laemmli (4X) supplemented with 10 % β-Mercaptoethanol and incubated at 95°C for 5 min. An equal amount of samples were then separated by SDS-PAGE. After the run, the gel was incubated in fixation solution (25 % Methanol, 8 % acetic acid in distilled water), and fluorescence was recorded using Amersham Typhoon.

### Aggregation assay in LLPS

20 µM of reduced or sulfonylated GAPDH was incubated with or without RNA (30 ng/µl) for 45 min at RT. The proteins were then mixed with 2 µl of PROTEOSTAT® (Enzo, #ENZ-51023) and 1 x Assay buffer as recommended by the manufacturer’s instructions. Samples were placed in a black 96-well plate with clear bottom (Invitrogen™, #M33089), and the plate was sealed. The proteostat fluorescence was measured every 3 min (with 1 min shaking at 300 rpm before measurement) using a CLARIOstar Plus plate reader (BMG LABTECH). The bottom optic setting was used to excite at 550-/+15 nm and read Proteostat emission at 600 nm.

### Fourier Transform Infrared Spectroscopy (FTIR) and secondary structures calculation

The Nicolet 6700 FTIR spectrometer (Thermo Scientific, Carlsbad, CA, USA) was used to record Infrared (IR) spectra of GAPDH samples in attenuated total reflectance (ATR) mode at 1 cm−1 resolution (*80*). Spectra of samples containing either reduced, sulfonylated or persulfidated GAPDH (30 µM) were analyzed in OMNIC software. Amide I region arises mainly from C = O stretching vibrations thus is very sensitive to changes in secondary structures. Between 1600–1700 cm−1 each secondary structure contributes to the absorption in a certain wavenumber range. The most prominent change detected in oxidized GAPDH after 8 h was signal increase in the regions attributed to beta-sheets in general, namely 1616–1625 cm−1 assigned to aggregation-prone inter/intramolecular beta-sheet, 1626–1640 cm−1 assigned to intramolecular beta-sheet.

### GAPDH aggregation in aging mice brain

Brain tissues were lysed in aggregation lysis buffer (50 mM Tris/HCl pH 8.0, 0.5 M NaCl, 4 mM EDTA, 1 % NP40) supplemented with 1 % Proteinase inhibitors. Frozen tissue was disrupted in lysis buffer using the TissueLyser II and stainless-steel beads. Samples were centrifuged 2-times at 2500 x g for 10 min at 4 °C to remove cell debris. An Equal amount of samples were subjected to ultracentrifugation at 500,000 x g for 15 min at 4°C. Supernatants were kept aside, presenting the soluble fraction. Pellets were washed with 0.5 % SDS in lysis buffer and again ultracentrifuged using the same setting. Pellets were washed again with 1 % SDS in lysis buffer and ultracentrifuged one last time. Pellets were resuspended in 50 mM HEPES pH 7.4, 2 % SDS forming the aggregate fraction. Both supernatant and aggregate fractions were mixed with 1 part of Laemmli (4X) supplemented with 10 % β-Mercaptoethanol and boiled at 95 °C for 15 min. An equal volume of samples was separated on SDS-PAGE and analyzed by immunoblotting using GAPDH-AF488 antibody.

### Thermal proteome profiling MEF WT and CSE-/- under H_2_O_2_ treatment

WT and CSE-/- MEFs were trypsinized and washed 3 times in cold serum-free DMEM. After the last centrifugation, cells were resuspended in 1.2 ml of DMEM supplemented with 10 % FCS and treated for 60 min with 500 µM H_2_O_2_ on ice. After 60 min, an equal number of cells were aliquoted into different tubes and gently centrifuged. Cell pellets were resuspended in 20 µl serum-free media and heated at 25, 47, or 57 °C for 3 min. Cells were brought back to room temperature for 3 min and lysed with freeze-thaw cycles. Extracted proteins were subjected to ultracentrifugation at 100,000 x g for 20 min at 4 °C. The volume corresponding to 50 µg in the 25 °C-exposed cells was used for all samples. Proteins were subsequently denatured using 3M Gu-HCl for reduction and alkylation steps with 500 µM DTT and 5 mM IAM for 30 min at 37 °C, respectively and then subjected to tryptic-digestion. Peptides were desalted as previously described. Eluates were evaporated to dry and resuspended in 100 mM triethylammonium bicarbonate for TMT labeling. Samples were labeled using the TMTpro™ 16plex Label Reagent Set. Samples were combined, desalted, and dried as previously described and subjected to LC/MS analysis by Orbitrap Eclipse™ Tribrid™ Mass Spectrometer via an EASY-spray (Thermo Fisher Scientific). Data were evaluated using FragPipe software (*81*) using 10 ppm for precursor mass tolerance, 0.02 Da for fragment mass tolerance, specific tryptic digest, and a maximum of 3 missed cleavages. Carbamidomethylation (+57.021464 Da), N-term acetylation (+42.010565 Da), and methionine oxidation (+15.994915 Da) were added as variable modifications. TMTpro (+304.207146 Da) was added as fixed modification on K and peptide N-term, peptide-spectrum match (PSM), and proteins were filtered at FDR 1 %. TMT quantification was performed at the MS2 level.

### Brain Section and Gallya’s staining

Brains were extracted and fixed in 4 % paraformaldehyde solution for 24 h at 4 °C, dehydrated in ethanol (30–100 %), and enlightened in xylene. After embedding in Histowax® (Histolab Product AB, Göteborg, Sweden), tissue blocs were sectioned to 5 μm thickness on a rotary microtome (RM 2125RT Leica Microsystems, Wetzlar, Germany). Gallya’s silver staining was performed following the manufacturer’s protocol (Morphisto, Offenbach am Main, Germany).

## Acknowledgments

We thank Ingo Feldman and Susanne Krois (ISAS e.V.) for their technical assistance with MS and the Advanced Medical Bioimaging Core Facility at Charité for the support.

## Author contributions

MRF conceived and supervised the study, designed experiments and analyzed and interpreted data. TV performed majority of the experiments, analyzed the data, prepared the figures. MRF and TV wrote the paper. DM provided LLPS expertise, designed synapsin experiments, supervised CH and HW and analyzed and interpreted data. MH, CH, AK, HW, JP, FC, MM, DP, JLJM also performed experiments and analyzed the data. AK, AP provided aging WT and CSE^-/-^ brain samples, performed lifespan experiments, interpreted and discussed the data. NP performed FT-IR measurements and analysis. JLJM, JLM, SRC, JP and MPM helped with G3BP2 real-time microscopy measurements and data interpretation. SC synthesized BTD. All authors edited the paper with input from the other authors.

## References and Notes

1. V. N. Gladyshev, Aging: progressive decline in fitness due to the rising deleteriome adjusted by genetic, environmental, and stochastic processes. Aging Cell. 15, 594–602 (2016).

2. D. HARMAN, Aging: a theory based on free radical and radiation chemistry. J. Gerontol. 11, 298–300 (1956).

3. C. E. Paulsen, K. S. Carroll, Cysteine-mediated redox signaling: Chemistry, biology, and tools for discovery. Chem. Rev. 113, 4633–4679 (2013).

4. D. Petrovic, E. Kouroussis, T. Vignane, M. R. Filipovic, The Role of Protein Persulfidation in Brain Aging and Neurodegeneration. Front. Aging Neurosci. 13, 1– 12 (2021).

5. J. Zivanovic, E. Kouroussis, J. B. Kohl, B. Adhikari, B. Bursac, S. Schott-Roux, D. Petrovic, J. L. Miljkovic, D. Thomas-Lopez, Y. Jung, M. Miler, S. Mitchell, V. Milosevic, J. E. Gomes, M. Benhar, B. Gonzales-Zorn, I. Ivanovic-Burmazovic, R. Torregrossa, J. R. Mitchell, M. Whiteman, G. Schwarz, S. H. Snyder, B. D. Paul, K. S. Carroll, M. R. Filipovic, Selective Persulfide Detection Reveals Evolutionarily Conserved Antiaging Effects of S-Sulfhydration. Cell Metab. 30, 1152–1170.e13 (2019).

6. A. Tyshkovskiy, P. Bozaykut, A. A. Borodinova, M. V. Gerashchenko, G. P. Ables, M. Garratt, P. Khaitovich, C. B. Clish, R. A. Miller, V. N. Gladyshev, Identification and Application of Gene Expression Signatures Associated with Lifespan Extension. Cell Metab. 30, 573–593.e8 (2019).

7. A. Tyshkovskiy, S. Ma, A. V. Shindyapina, S. Tikhonov, S. G. Lee, P. Bozaykut, J. P. Castro, A. Seluanov, N. J. Schork, V. Gorbunova, S. E. Dmitriev, R. A. Miller, V. N. Gladyshev, Distinct longevity mechanisms across and within species and their association with aging. Cell (2023), doi:10.1016/j.cell.2023.05.002.

8. D. Hanna, R. Kumar, R. Banerjee, A Metabolic Paradigm for Hydrogen Sulfide Signaling via Electron Transport Chain Plasticity. Antioxid. Redox Signal. 38 (2023), doi:10.1089/ARS.2022.0067.

9. C. P. Brangwynne, C. R. Eckmann, D. S. Courson, A. Rybarska, C. Hoege, J. Gharakhani, F. Jülicher, A. A. Hyman, Germline P granules are liquid droplets that localize by controlled dissolution/condensation. Science (80-.). 324, 1729–1732 (2009).

10. S. F. Banani, H. O. Lee, A. A. Hyman, M. K. Rosen, Biomolecular condensates: Organizers of cellular biochemistry. Nat. Rev. Mol. Cell Biol. 18, 285–298 (2017).

11. A. A. Hyman, C. A. Weber, F. Jülicher, Liquid-liquid phase separation in biology. Annu. Rev. Cell Dev. Biol. 30, 39–58 (2014).

12. S. Alberti, A. A. Hyman, Biomolecular condensates at the nexus of cellular stress, protein aggregation disease and ageing. Nat. Rev. Mol. Cell Biol. 22, 196–213 (2021).

13. Y. Shin, C. P. Brangwynne, Liquid phase condensation in cell physiology and disease. Science (80-.). 357 (2017), doi:10.1126/science.aaf4382.

14. T. M. Franzmann, M. Jahnel, A. Pozniakovsky, J. Mahamid, A. S. Holehouse, E. Nüske, D. Richter, W. Baumeister, S. W. Grill, R. V. Pappu, A. A. Hyman, S. Alberti, Phase separation of a yeast prion protein promotes cellular fitness. Science (80-.). 359 (2018), doi:10.1126/science.aao5654.

15. S. Wegmann, B. Eftekharzadeh, K. Tepper, K. M. Zoltowska, R. E. Bennett, S. Dujardin, P. R. Laskowski, D. MacKenzie, T. Kamath, C. Commins, C. Vanderburg, A. D. Roe, Z. Fan, A. M. Molliex, A. Hernandez-Vega, D. Muller, A. A. Hyman, E. Mandelkow, J. P. Taylor, B. T. Hyman, Tau protein liquid–liquid phase separation can initiate tau aggregation. EMBO J. 37, 1–21 (2018).

16. S. Ray, N. Singh, R. Kumar, K. Patel, S. Pandey, D. Datta, J. Mahato, R. Panigrahi, A. Navalkar, S. Mehra, L. Gadhe, D. Chatterjee, A. S. Sawner, S. Maiti, S. Bhatia, J. A. Gerez, A. Chowdhury, A. Kumar, R. Padinhateeri, R. Riek, G. Krishnamoorthy, S. K. Maji, α-Synuclein aggregation nucleates through liquid–liquid phase separation. Nat. Chem. 12, 705–716 (2020).

17. B. Wolozin, P. Ivanov, Stress granules and neurodegeneration. Nat. Rev. Neurosci. 20, 649–666 (2019).

18. C. López-Otín, M. A. Blasco, L. Partridge, M. Serrano, G. Kroemer, The hallmarks of aging. Cell. 153, 1194 (2013).

19. C. López-Otín, M. A. Blasco, L. Partridge, M. Serrano, G. Kroemer, Hallmarks of aging: An expanding universe. Cell (2023), doi:10.1016/j.cell.2022.11.001.

20. I. Harel, Y. R. Chen, I. Ziv, P. P. Singh, P. N. Negredo, U. Goshtchevsky, W. Wang, G. Astre, E. Moses, A. McKay, B. E. Machado, K. Hebestreit, S. Yin, A. S. Alvarado, D. F. Jarosz, A. Brunet, bioRxiv, in press (available at https://www.biorxiv.org/content/10.1101/2022.02.26.482115v1%0Ahttps://www.biorxiv.org/content/10.1101/2022.02.26.482115v1.abstract).

21. S. N. Mouton, D. J. Thaller, M. M. Crane, I. L. Rempel, O. Terpstra, A. Steen, M. Kaeberlein, C. P. Lusk, A. J. Boersma, L. M. Veenhoff, A physicochemical perspective of aging from single-cell analysis of ph, macromolecular and organellar crowding in yeast. Elife (2020), doi:10.7554/ELIFE.54707.

22. H. Xiao, M. P. Jedrychowski, D. K. Schweppe, E. L. Huttlin, Q. Yu, D. E. Heppner, J. Li, J. Long, E. L. Mills, J. Szpyt, Z. He, G. Du, R. Garrity, A. Reddy, L. P. Vaites, J. A. Paulo, T. Zhang, N. S. Gray, S. P. Gygi, E. T. Chouchani, A Quantitative Tissue-Specific Landscape of Protein Redox Regulation during Aging. Cell. 180, 968–983.e24 (2020).

23. J. A. Reisz, E. Bechtold, S. B. King, L. B. Poole, C. M. Furdui, Thiol-blocking electrophiles interfere with labeling and detection of protein sulfenic acids. FEBS J. 280, 6150–6161 (2013).

24. Dóka, T. Ida, M. Dagnell, Y. Abiko, N. C. Luong, N. Balog, T. Takata, B. Espinosa, A. Nishimura, Q. Cheng, Y. Funato, H. Miki, J. M. Fukuto, J. R. Prigge, E. E. Schmidt, E. S. J. Arnér, Y. Kumagai, T. Akaike, P. Nagy, Control of protein function through oxidation and reduction of persulfidated states. Sci. Adv. 6 (2020), doi:10.1126/sciadv.aax8358.

25. J. Yang, V. Gupta, K. S. Carroll, D. C. Liebler, Site-specific mapping and quantification of protein S-sulphenylation in cells. Nat. Commun. 5, 4776 (2014).

26. L. Fu, K. Liu, R. B. Ferreira, K. S. Carroll, J. Yang, Proteome-Wide Analysis of Cysteine S-Sulfenylation Using a Benzothiazine-Based Probe. Curr. Protoc. Protein Sci. 95, e76 (2019).

27. E. Kelmer Sacramento, J. M. Kirkpatrick, M. Mazzetto, M. Baumgart, A. Bartolome, S. Di Sanzo, C. Caterino, M. Sanguanini, N. Papaevgeniou, M. Lefaki, D. Childs, S. Bagnoli, E. Terzibasi Tozzini, D. Di Fraia, N. Romanov, P. H. Sudmant, W. Huber, N. Chondrogianni, M. Vendruscolo, A. Cellerino, A. Ori, Reduced proteasome activity in the aging brain results in ribosome stoichiometry loss and aggregation. Mol. Syst. Biol. 16, 1–22 (2020).

28. L. Fu, K. Liu, J. He, C. Tian, X. Yu, J. Yang, Direct Proteomic Mapping of Cysteine Persulfidation. Antioxidants Redox Signal. 33, 1061–1076 (2020).

29. S. I. Bibli, J. Hu, M. Looso, A. Weigert, C. Ratiu, J. Wittig, M. K. Drekolia, L. Tombor, V. Randriamboavonjy, M. S. Leisegang, P. Goymann, F. Delgado Lagos, B. Fisslthaler, S. Zukunft, A. Kyselova, A. F. O. Justo, J. Heidler, D. Tsilimigras, R. P. Brandes, S. Dimmeler, A. Papapetropoulos, S. Knapp, S. Offermanns, I. Wittig, S. L. Nishimura, F. Sigala, I. Fleming, Mapping the Endothelial Cell S -Sulfhydrome Highlights the Crucial Role of Integrin Sulfhydration in Vascular Function. Circulation. 143, 935–948 (2021).

30. N. Bithi, C. Link, Y. O. Henderson, S. Kim, J. Yang, L. Li, R. Wang, B. Willard, C. Hine, Dietary restriction transforms the mammalian protein persulfidome in a tissue-specific and cystathionine γ-lyase-dependent manner. Nat. Commun. 12 (2021), doi:10.1038/s41467-021-22001-w.

31. D. Petrovic, E. Kouroussis, T. Vignane, M. R. Filipovic, The Role of Protein Persulfidation in Brain Aging and Neurodegeneration. Front. Aging Neurosci. (2021), doi:10.3389/fnagi.2021.674135.

32. D. Giovinazzo, B. Bursac, J. I. Sbodio, S. Nalluru, T. Vignane, A. M. Snowman, L. M. Albacarys, T. W. Sedlak, R. Torregrossa, M. Whiteman, M. R. Filipovic, S. H. Snyder, B. D. Paul, Hydrogen sulfide is neuroprotective in Alzheimer’s disease by sulfhydrating GSK3β and inhibiting Tau hyperphosphorylation. Proc. Natl. Acad. Sci. U. S. A. 118, e2017225118 (2021).

33. P. M. Snijder, M. Baratashvili, N. A. Grzeschik, H. G. D. Leuvenink, L. Kuijpers, S. Huitema, O. Schaap, B. N. G. Giepmans, J. Kuipers, J. L. Miljkovic, A. Mitrovic, E. M. Bos, C. Szabó, H. H. Kampinga, P. F. Dijkers, W. F. A. den Dunnen, M. R. Filipovic, H. van Goor, O. C. M. Sibon, Overexpression of Cystathionine γ-Lyase Suppresses Detrimental Effects of Spinocerebellar Ataxia Type 3. Mol. Med. 21, 758–768 (2015).

34. B. D. Paul, J. I. Sbodio, R. Xu, M. S. Vandiver, J. Y. Cha, A. M. Snowman, S. H. Snyder, Cystathionine γ-lyase deficiency mediates neurodegeneration in Huntington’s disease. Nature. 508, 96–100 (2014).

35. C. Margreitter, D. Petrov, B. Zagrovic, Vienna-PTM web server: a toolkit for MD simulations of protein post-translational modifications. Nucleic Acids Res. 41, 422– 426 (2013).

36. D. Milovanovic, Y. Wu, X. Bian, P. De Camilli, A liquid phase of synapsin and lipid vesicles. Science (80-.). 361, 604–607 (2018).

37. V. Gupta, H. Paritala, K. S. Carroll, Reactivity, Selectivity, and Stability in Sulfenic Acid Detection: A Comparative Study of Nucleophilic and Electrophilic Probes. Bioconjug. Chem. 27, 1411–1418 (2016).

38. M. R. Filipovic, J. Zivanovic, B. Alvarez, R. Banerjee, Chemical Biology of H2S Signaling through Persulfidation. Chem. Rev. 118 (2018), pp. 1253–1337.

39. M. Kato, Y. S. Yang, B. M. Sutter, Y. Wang, S. L. McKnight, B. P. Tu, Redox State Controls Phase Separation of the Yeast Ataxin-2 Protein via Reversible Oxidation of Its Methionine-Rich Low-Complexity Domain. Cell. 177, 711–721.e8 (2019).

40. K. Abe, H. Kimura, The possible role of hydrogen sulfide as an endogenous neuromodulator. J. Neurosci. 16, 1066–1071 (1996).

41. P. K. Kamat, A. Kalani, N. Tyagi, Role of hydrogen sulfide in brain synaptic remodeling. Methods Enzymol. 555, 207–229 (2015).

42. R. Ali, H. A. Pal, R. Hameed, A. Nazir, S. Verma, Controlled release of hydrogen sulfide significantly reduces ROS stress and increases dopamine levels in transgenic C. elegans. Chem. Commun. 55, 10142–10145 (2019).

43. A. Molliex, J. Temirov, J. Lee, M. Coughlin, A. P. Kanagaraj, H. J. Kim, T. Mittag, J. P. Taylor, Phase Separation by Low Complexity Domains Promotes Stress Granule Assembly and Drives Pathological Fibrillization. Cell. 163, 123–133 (2015).

44. D. W. Sanders, N. Kedersha, D. S. W. Lee, A. R. Strom, V. Drake, J. A. Riback, D. Bracha, J. M. Eeftens, A. Iwanicki, A. Wang, M. T. Wei, G. Whitney, S. M. Lyons, P. Anderson, W. M. Jacobs, P. Ivanov, C. P. Brangwynne, Competing Protein-RNA Interaction Networks Control Multiphase Intracellular Organization. Cell. 181, 306–324.e28 (2020).

45. P. Yang, C. Mathieu, R. M. Kolaitis, P. Zhang, J. Messing, U. Yurtsever, Z. Yang, J. Wu, Y. Li, Q. Pan, J. Yu, E. W. Martin, T. Mittag, H. J. Kim, J. P. Taylor, G3BP1 Is a Tunable Switch that Triggers Phase Separation to Assemble Stress Granules. Cell. 181, 325–345.e28 (2020).

46. J. Guillén-Boixet, A. Kopach, A. S. Holehouse, S. Wittmann, M. Jahnel, R. Schlüßler, K. Kim, I. R. E. A. Trussina, J. Wang, D. Mateju, I. Poser, S. Maharana, M. Ruer-Gruß, D. Richter, X. Zhang, Y. T. Chang, J. Guck, A. Honigmann, J. Mahamid, A. A. Hyman, R. V. Pappu, S. Alberti, T. M. Franzmann, RNA-Induced Conformational Switching and Clustering of G3BP Drive Stress Granule Assembly by Condensation. Cell. 181, 346–361.e17 (2020).

47. M. Kato, T. W. Han, S. Xie, K. Shi, X. Du, L. C. Wu, H. Mirzaei, E. J. Goldsmith, J. Longgood, J. Pei, N. V. Grishin, D. E. Frantz, J. W. Schneider, S. Chen, L. Li, M. R. Sawaya, D. Eisenberg, R. Tycko, S. L. McKnight, Cell-free formation of RNA granules: Low complexity sequence domains form dynamic fibers within hydrogels. Cell. 149, 753–767 (2012).

48. S. F. Banani, A. M. Rice, W. B. Peeples, Y. Lin, S. Jain, R. Parker, M. K. Rosen, Compositional Control of Phase-Separated Cellular Bodies. Cell. 166, 651–663 (2016).

49. A. Khong, T. Matheny, S. Jain, S. F. Mitchell, J. R. Wheeler, R. Parker, The Stress Granule Transcriptome Reveals Principles of mRNA Accumulation in Stress Granules. Mol. Cell. 68, 808–820.e5 (2017).

50. D. Kennedy, J. French, E. Guitard, K. Ru, B. Tocque, J. Mattick, Characterization of G3BPs: Tissue specific expression, chromosomal localisation and rasGAP120 binding studies. J. Cell. Biochem. 84, 173–187 (2002).

51. J. M. Hourihan, L. E. Moronetti Mazzeo, L. P. Fernández-Cárdenas, T. K. Blackwell, Cysteine Sulfenylation Directs IRE-1 to Activate the SKN-1/Nrf2 Antioxidant Response. Mol. Cell. 63, 553–566 (2016).

52. V. Vitvitsky, O. Kabil, R. Banerjee, High Turnover Rates for Hydrogen Sulfide Allow for Rapid Regulation of Its Tissue Concentrations. Antioxid. Redox Signal. 17, 22– 31 (2012).

53. R. Wedmann, S. Bertlein, I. Macinkovic, S. Böltz, J. L. Miljkovic, L. E. Muñoz, M. Herrmann, M. R. Filipovic, Working with “H2S”: Facts and apparent artifacts. Nitric Oxide - Biol. Chem. 41, 85–96 (2014).

54. G. G. Fuller, T. Han, M. A. Freeberg, J. J. Moresco, A. G. Niaki, N. P. Roach, J. R. Yates, S. Myong, J. K. Kim, Rna promotes phase separation of glycolysis enzymes into yeast g bodies in hypoxia. Elife. 9, 1–30 (2020).

55. M. Jin, T. Han, Y. Yao, A. F. Alessi, M. A. Freeberg, K. Inoki, D. J. Klionsky, J. K. Kim, A. Karnovsky, J. J. Moresco, J. R. Yates, M. Baba, A. D. Gitler, G. G. Fuller, A. F. Alessi, N. P. Roach, Glycolytic Enzymes Coalesce in G Bodies under Hypoxic Stress. Cell Rep. 20, 895–908 (2017).

56. A. Zeida, M. Trujillo, G. Ferrer-Sueta, A. Denicola, D. A. Estrin, R. Radi, Catalysis of Peroxide Reduction by Fast Reacting Protein Thiols. Chem. Rev. 119, 10829– 10855 (2019).

57. M. Dagnell, Q. Cheng, S. H. M. Rizvi, P. E. Pace, B. Boivin, C. C. Winterbourn, E. S. J. Arnér, Bicarbonate is essential for protein-tyrosine phosphatase 1B (PTP1B) oxidation and cellular signaling through EGF-triggered phosphorylation cascades. J. Biol. Chem. 294, 12330–12338 (2019).

58. Q. Wang, R. L. Woltjer, P. J. Cimino, C. Pan, K. S. Montine, J. Zhang, T. J. Montine, Proteomic analysis of neurofibrillary tangles in Alzheimer disease identifies GAPDH as a detergent-insoluble paired helical filament tau binding protein. FASEB J. 19, 1–12 (2005).

59. V. F. Lazarev, M. Tsolaki, E. R. Mikhaylova, K. A. Benken, M. A. Shevtsov, A. D. Nikotina, M. Lechpammer, V. A. Mitkevich, A. A. Makarov, A. A. Moskalev, S. A. Kozin, B. A. Margulis, I. V. Guzhova, E. Nudler, Extracellular gapdh promotes alzheimer disease progression by enhancing amyloid-β aggregation and cytotoxicity. Aging Dis. 12, 1223–1237 (2021).

60. B. H. Patel, C. Percivalle, D. J. Ritson, C. D. Duffy, J. D. Sutherland, Common origins of RNA, protein and lipid precursors in a cyanosulfidic protometabolism. Nat. Chem. 7, 301–307 (2015).

61. C. S. Foden, S. Islam, C. Fernández-García, L. Maugeri, T. D. Sheppard, M. W. Powner, Prebiotic synthesis of cysteine peptides that catalyze peptide ligation in neutral water. Science (80-.). 370, 865–869 (2020).

62. S. W. Fox, The evolutionary significance of phase-separated microsystems. Orig. Life. 7, 49–68 (1976).

63. E. Spruijt, Open questions on liquid–liquid phase separation. Commun. Chem. 6, 1– 5 (2023).

64. P. Principles, NASA Public Access. 57, 2509–2519 (2020).

65. B. D’Autréaux, M. B. Toledano, ROS as signalling molecules: Mechanisms that generate specificity in ROS homeostasis. Nat. Rev. Mol. Cell Biol. 8, 813–824 (2007).

66. Y. Dai, C. F. Chamberlayne, M. S. Messina, C. J. Chang, R. N. Zare, L. You, A. Chilkoti, Interface of biomolecular condensates modulates redox reactions. Chem. 9, 1594–1609 (2023).

67. R. S. Balaban, S. Nemoto, T. Finkel, Mitochondria, oxidants, and aging. Cell. 120, 483–495 (2005).

68. J. M. Van Raamsdonk, S. Hekimi, Reactive oxygen species and aging in caenorhabditis elegans: Causal or casual relationship? Antioxidants Redox Signal. 13, 1911–1953 (2010).

69. D. Knoefler, M. Thamsen, M. Koniczek, N. J. Niemuth, A. K. Diederich, U. Jakob, Quantitative In Vivo Redox Sensors Uncover Oxidative Stress as an Early Event in Life. Mol. Cell. 47, 767–776 (2012).

70. D. Bazopoulou, D. Knoefler, Y. Zheng, K. Ulrich, B. J. Oleson, L. Xie, M. Kim, A. Kaufmann, Y. T. Lee, Y. Dou, Y. Chen, S. Quan, U. Jakob, Developmental ROS individualizes organismal stress resistance and lifespan. Nature. 576, 301–305 (2019).

71. L. M. Redman, S. R. Smith, J. H. Burton, C. K. Martin, D. Il’yasova, E. Ravussin, Metabolic Slowing and Reduced Oxidative Damage with Sustained Caloric Restriction Support the Rate of Living and Oxidative Damage Theories of Aging. Cell Metab. 27, 805–815.e4 (2018).

72. G. Yang, L. Wu, B. Jiang, W. Yang, J. Qi, K. Cao, Q. Meng, A. K. Mustafa, W. Mu, S. Zhang, S. H. Snyder, R. Wang, H2S as a physiologic vasorelaxant: Hypertension in mice with deletion of cystathionine γ-lyase. Science (80-.). 322, 587–590 (2008).

73. J. I. Sbodio, S. H. Snyder, B. D. Paul, Transcriptional control of amino acid homeostasis is disrupted in Huntington’s disease. Proc. Natl. Acad. Sci. 113, 8843– 8848 (2016).

74. Y. V. Karpievitch, T. Taverner, J. N. Adkins, S. J. Callister, G. A. Anderson, R. D. Smith, A. R. Dabney, Normalization of peak intensities in bottom-up MS-based proteomics using singular value decomposition. Bioinformatics. 25, 2573–2580 (2009).

75. C. Hoffmann, R. Sansevrino, G. Morabito, C. Logan, R. M. Vabulas, A. Ulusoy, M. Ganzella, D. Milovanovic, Synapsin Condensates Recruit alpha-Synuclein. J. Mol. Biol. 433, 166961 (2021).

76. C. Hoffmann, J. Rentsch, T. A. Tsunoyama, A. Chhabra, G. Aguilar Perez, R. Chowdhury, F. Trnka, A. A. Korobeinikov, A. H. Shaib, M. Ganzella, G. Giannone, S. O. Rizzoli, A. Kusumi, H. Ewers, D. Milovanovic, Synapsin condensation controls synaptic vesicle sequestering and dynamics. Nat. Commun. 14, 6730 (2023).

77. F. Katzen, Gateway® recombinational cloning: A biological operating system. Expert Opin. Drug Discov. 2 (2007), doi:10.1517/17460441.2.4.571.

78. A. Agrotis, N. Pengo, J. J. Burden, R. Ketteler, Redundancy of human ATG4 protease isoforms in autophagy and LC3/GABARAP processing revealed in cells. Autophagy. 15, 976–997 (2019).

79. H. Liu, J. H. Naismith, An efficient one-step site-directed deletion, insertion, single and multiple-site plasmid mutagenesis protocol. BMC Biotechnol. 8 (2008), doi:10.1186/1472-6750-8-91.

80. S. Marković, N. S. Andrejević, J. Milošević, N. n. Polović, Structural Transitions of Papain-like Cysteine Proteases: Implications for Sensor Development. Biomimetics. 8 (2023), doi:10.3390/biomimetics8030281.

81. A. T. Kong, F. V. Leprevost, D. M. Avtonomov, D. Mellacheruvu, A. I. Nesvizhskii, MSFragger: Ultrafast and comprehensive peptide identification in mass spectrometry-based proteomics. Nat. Methods. 14, 513–520 (2017).

