## Supplementary Figures for "Protein thiol alterations drive aberrant phase separation in aging"

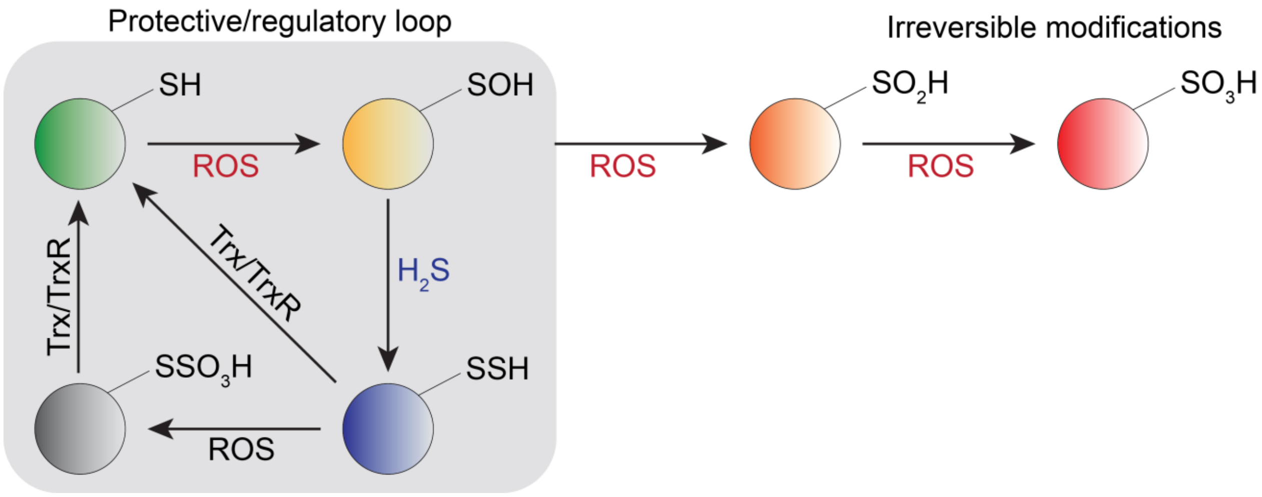


**Supporting Figure S1. Proposed protective effects of protein persulfidation.** During the oxidative stress cysteine residues get oxidized to sulfenic acids. This represents an important signaling event for the cell to either start proliferating or to die (depending on the amount of $H_2O_2$). However, left unreacted or exposed to further ROS, PSOH become oxidized to sulfinic ($PSO_2H$) and sulfonic acids ($PSO_3H$); these modifications are generally irreversible. Furthermore, if buried deep into protein pockets, PSOH could become stabilized and not easily reachable for the reduction. $H_2S$ is small and can reach deep into protein structure. Once formed, PSSH can be reduced back to thiols by the thioredoxin (Trx) system. When the oxidative stress persists (like in aging and many ROS-related diseases), PSSH will act as better scavengers of ROS than PSH resulting in the formation of $PSSO_3H$. The existence of S-S bond makes this species a target for Trx and the restoration of native thiolate. That way the overall structure, function and half-life of thiol-containing protein gets preserved. We propose that this rescue loop exemplifies a remnant of the times when life emerged in an $H_2S$-rich environment and that it represents the simplest way to resolve cysteine oxidation and protect proteins from oxidative damage.

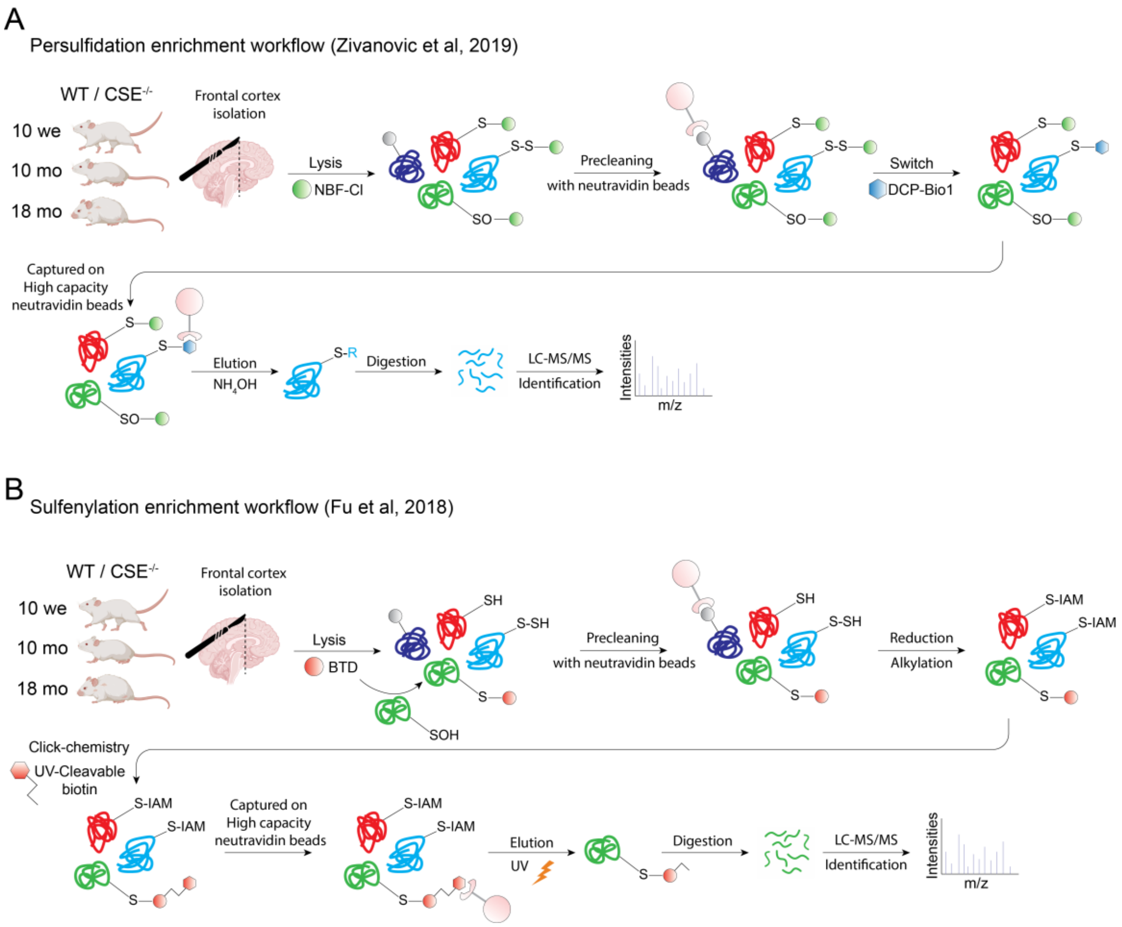


**Supporting Figure S2. Chemoproteomic workflows of sulfenylation and persulfidation enrichment. (A)** The persulfidation enrichment workflow is based on Zivanovic et al. 2019, protocol (*5*). The frontotemporal part of the brain is extracted and lysed in the presence of 5 mM 4-Chloro-7-nitrobenzofurazan (NBF-Cl), blocking all free cysteine and stabilizing persulfides. Endogenously biotinylated proteins (grey) are removed with a pre-cleaning step using neutravidin beads. Persulfides are then selectively labeled using DCP-Bio1 dimedone-based probe. Although it has been shown that dimedone-based probes could also have minor side reaction with very reactive amino groups (*72*), in this method they are already blocked with NBF-Cl. Persulfidated proteins are then pulled down with high-capacity neutravidin beads. To selectively elute the proteins, ammonia ($NH_4OH$) is used, cleaving the probe and releasing the protein from the beads. Once eluted, proteins are digested using trypsin, and peptides are identified by mass spectrometry. (**B**) The sulfenylation enrichment workflow is modified from Fu et al, 2018 (*26*). The frontotemporal part of the brain is extracted and lysed in the presence of 500 µM benzothiazine-based chemoselective probe (BTD). BTD is one of the most selective probes for sulfenylation and has been shown to have no side reactivity with other amino acids (*72*). To avoid unspecific enrichment, the samples are cleaned from endogenous biotinylated protein using neutravidin beads. UV-cleavable biotin probe is then attached to the BTD probe by using click-chemistry and sulfenylated proteins captured on high-capacity neutravidin beads. These proteins can be selectively eluted using UV light that cleaves the probe and releases the protein from the beads. Once eluted, proteins are digested and peptides identified using mass spectrometry.

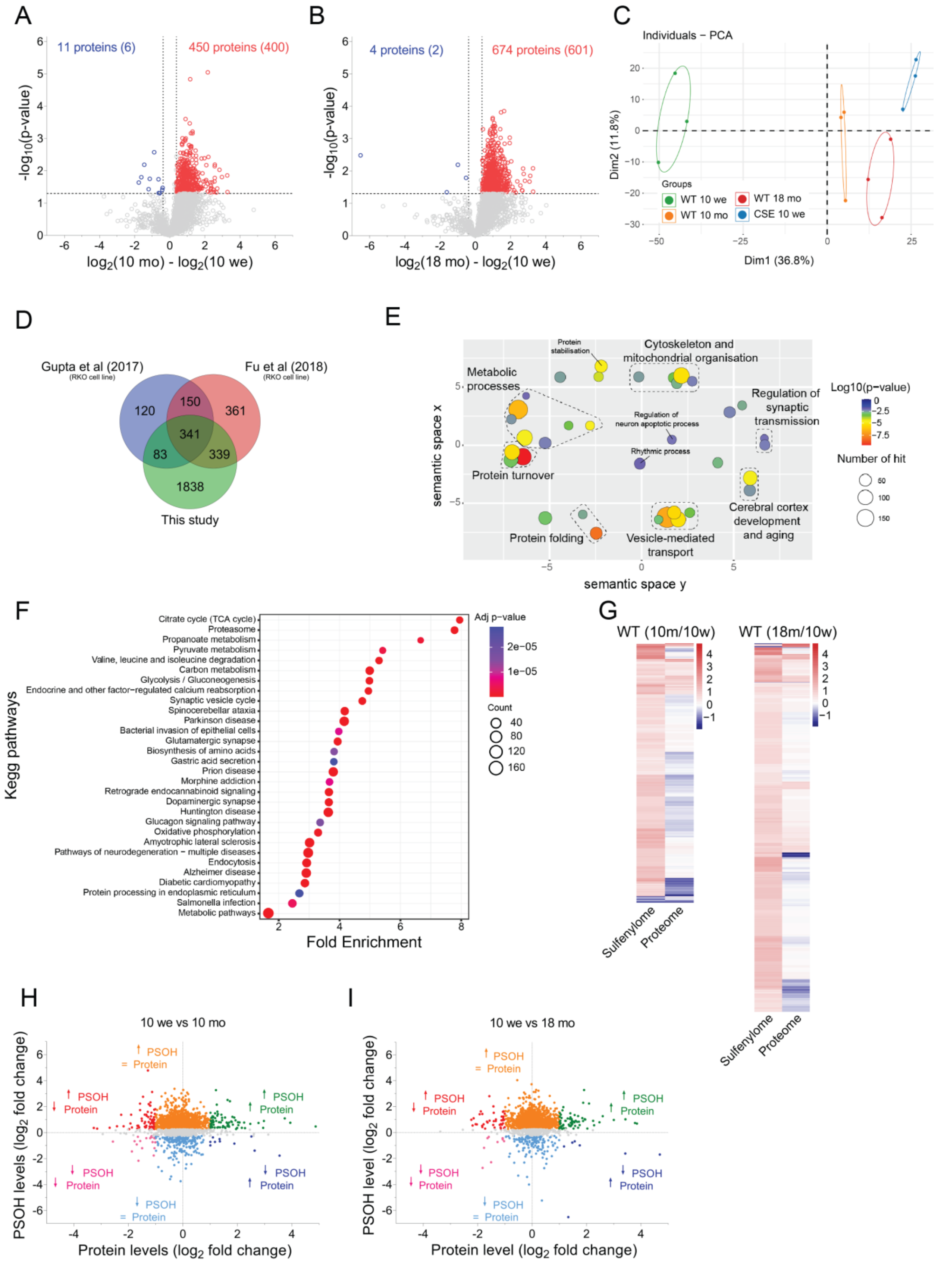
A
11 proteins (6)
450 proteins (400)
-log10(p-value)
log2(10 mo) - log2(10 we)
B
4 proteins (2)
674 proteins (601)
-log10(p-value)
log2(18 mo) - log2(10 we)
C
Individuals − PCA
Groups
WT 10 we
WT 10 mo
WT 18 mo
CSE 10 we
Dim1 (36.8%)
Dim2 (11.8%)
D
Gupta et al (2017)
(RKO cell line)
Fu et al (2018)
(RKO cell line)
120
150
361
341
83
339
1838
This study
E
Protein stabilisation
Cytoskeleton and mitochondrial organisation
Metabolic processes
Regulation of synaptic transmission
Regulation of neuron apoptotic process
Rhythmic process
Protein turnover
Cerebral cortex development and aging
Protein folding
Vesicle-mediated transport
semantic space x
semantic space y
Log10(p−value)
Number of hit
F
Kegg pathways
Citrate cycle (TCA cycle)
Proteasome
Propanoate metabolism
Pyruvate metabolism
Valine, leucine and isoleucine degradation
Carbon metabolism
Glycolysis / Gluconeogenesis
Endocrine and other factor−regulated calcium reabsorption
Synaptic vesicle cycle
Spinocerebellar ataxia
Parkinson disease
Bacterial invasion of epithelial cells
Glutamatergic synapse
Biosynthesis of amino acids
Gastric acid secretion
Prion disease
Morphine addiction
Retrograde endocannabinoid signaling
Dopaminergic synapse
Huntington disease
Glucagon signaling pathway
Oxidative phosphorylation
Amyotrophic lateral sclerosis
Pathways of neurodegeneration − multiple diseases
Endocytosis
Alzheimer disease
Diabetic cardiomyopathy
Protein processing in endoplasmic reticulum
Salmonella infection
Metabolic pathways
Fold Enrichment
Adj p-value
Count
G
WT (10m/10w)
WT (18m/10w)
Sulfenylome
Proteome
H
10 we vs 10 mo
PSOH
Protein
PSOH levels (log2 fold change)
Protein levels (log2 fold change)
I
10 we vs 18 mo
PSOH level (log2 fold change)
Protein level (log2 fold change)

**Supporting Figure SS3. Chemoproteomic quantitative analysis of sulfenylome during aging.** (**A-B**) **Volcano plot analysis of sulfenylome changes.** Volcano plot illustrating the statistical significance plotted against the $\log_2$-fold change of sulfenylated proteins in 10-month-old mice relative to 10-week-old mice (**A**) and 18-month-old mice relative to 10-week-old mice (**B**) (n = 3). Significance was controlled by Welch's t-test (two-sided) with a p-value threshold of <0.05, while a fold change cut-off of 30 % was applied. Sulfenylated proteins showing a significant increase in 10-month-old mice are highlighted in vivid blue, whereas proteins displaying significant decreases are demarcated in striking red. The number of proteins with significantly altered sulfenylation states (either increase or decrease) is indicated, with the number in parentheses reflecting proteins identified through a minimum of 2 unique peptides. (**C**) Principal-component analysis (PCA) plot elucidating the reproducibility across all three replicates. Notably, 10-month and 18-month-old mice distinctly separate from their 10-week-old counterparts. Furthermore, a discernible pattern emerges between 10-week-old mice lacking CSE ($CSE^{-/-}$) in contrast to wild-type (WT) 10-week-old mice. (**D**) Venn diagram comparing the proteins identified within our study against two other data sets (*26*, *73*). Both studies used the same method, but the enrichment was done at the peptide level. (**E**) GO term enrichment (biological process) of the 912 proteins significantly increased at least once across the age. Employing DAVID for enrichment, the outcomes were visualized through REVIGO. Significant GO terms passed the Benjamini adjusted p-value threshold of 0.01. Circle dimensions denote the protein count within specific GO terms, while color gradients communicate the degree of significance. (**F**) Kegg pathway enrichment analysis was performed using DAVID. The graph shows the top 30 significant (Benjamini adjusted p-value < 0.01) and most enriched pathways, with the color gradient indicating adjusted p-values and circle size the number of proteins. (**G**) Heatmaps representing the $\log_2$-fold change of the significantly changing sulfenylated proteins (Welch's test p-value < 0.05) in 10-month or 18-month-old mice compared to 10-week-old animals and their corresponding $\log_2$-fold change of the protein expression level. A notable observation is the limited correlation between the modification and the expression of the proteins, validating the sulfenylome changes. This delineation implies that substantial sulfenylome shifts are authentic and not contingent upon protein expression alterations. (**H-I**) Scatter plot analysis unveiling sulfenylome and protein expression relationships. (**H**) Scatter plot characterizing log2 fold changes within the sulfenylome (y-axis) and protein expression (x-axis) for comparison between 10-month-old and 10-week-old mice. (**I**) Parallel scatter plot analysis for the comparison between 18-month-old and 10-week-old mice. Most of the proteins undergoing sulfenylome modifications do not surpass the significance threshold in protein expression. This observation underscores that the sulfenylome changes detected remain largely uncoupled from concurrent change in protein expression.

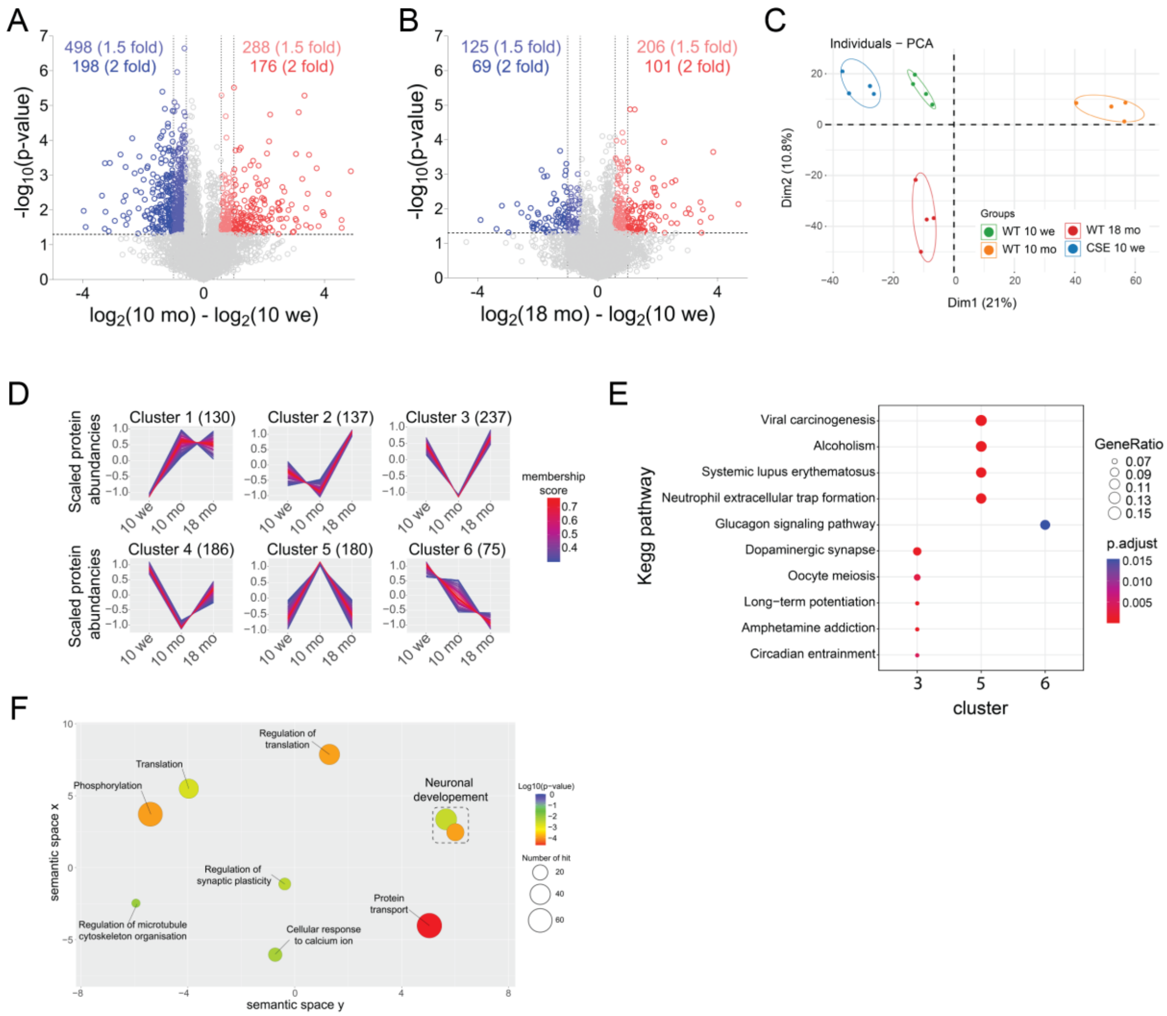


**Supporting Figure S4. Quantitative profiling unveils proteome dynamics during aging.** (**A-B**) Volcano plot analysis of proteome changes. Volcano plot illustrating statistical significance plotted against the $\log_2$-fold change of protein levels in 10-month-old mice relative to 10-week-old mice (**A**) and 18-month-old mice relative to 10-week-old mice (**B**) (n = 4). Significance was controlled by Welch's t-test (two-sided), with a p-value threshold of < 0.05. Fold change cut-offs were set at both 1.5-fold and 2-fold. Notably, proteins exhibiting significant increases are visually represented in light red (fold change > 1.5) or red (fold change > 2), while significantly decreased proteins appear in light blue (fold change > 1.5) or blue (fold change > 2). (**C**) Principal component analysis underscoring the reproducibility across all replicates. The analysis discerns the patterns characterizing aging and highlights the distinct protein expression profile of $CSE^{-/-}$ mice in the aging context. (**D**) Hierarchical cluster analysis of proteins displaying significant changes at least at one time point during aging. (**E**) Kegg pathway enrichment analysis of protein clusters. Only clusters 3, 5, and 6 display significant enrichment in Kegg pathway terms (Benjamini corrected p-value < 0.05). (**F**) Gene Ontology (GO) term enrichment analysis targeting biological processes encompassing the 1035 proteins displaying age-related changes. REVIGO was used to plot the enrichment analysis performed in DAVID. Significant GO terms passed the Benjamini adjusted p-value threshold of 0.01. Circle dimensions denote the protein count within specific GO terms, while color gradients represent the degree of significance.

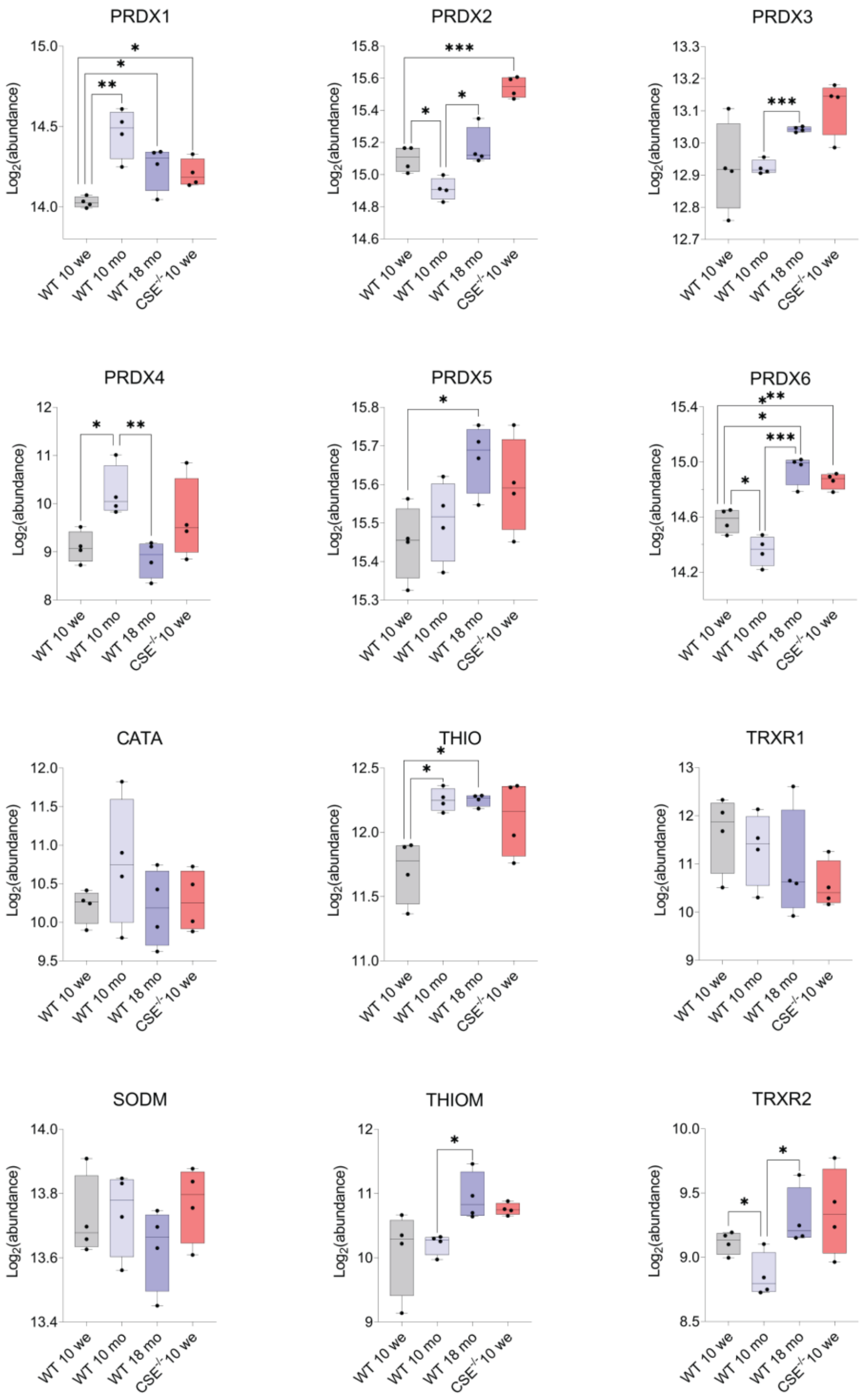


**Supporting Figure S5. Expression profiles of anti-oxidant proteins across aging and in $CSE^{-/-}$ mice.** Whiskers and box plots of the Log2 transformed abundancies of the anti-oxidant proteins identified in the proteome analysis. Reported significance levels accompanying the plots are directly correlated with the p-values calculated through the proteome analysis (Welch's t-test, * < 0.05, ** < 0.01, *** < 0.001). PRDX (1-6): peroxiredoxin (1-6); CATA : Catalase; THIO: thioredoxin, THIOM: mitochondrial thioredoxin; TRXR (1-2): thioredoxin reductase (1-2); SODM: mitochondrial superoxide dismutase.

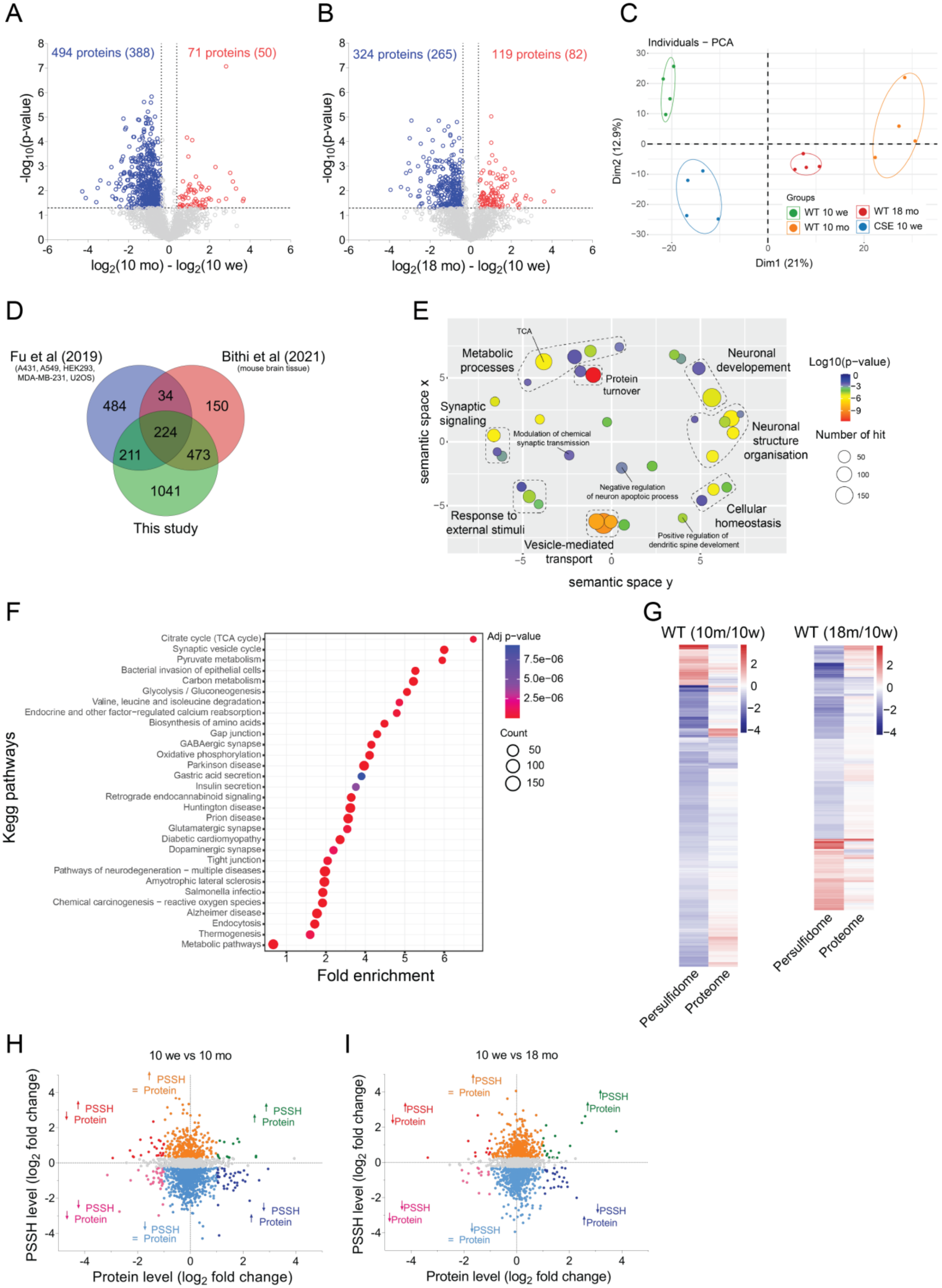
A
494 proteins (388)
71 proteins (50)
-log10(p-value)
log2(10 mo) - log2(10 we)
B
324 proteins (265)
119 proteins (82)
-log10(p-value)
log2(18 mo) - log2(10 we)
C
Individuals − PCA
Dim2 (12.9%)
Dim1 (21%)
Groups
WT 10 we
WT 18 mo
WT 10 mo
CSE 10 we
D
Fu et al (2019)
(A431, A549, HEK293, MDA-MB-231, U2OS)
Bithi et al (2021)
(mouse brain tissue)
484
34
150
224
211
473
1041
This study
E
TCA
Metabolic processes
Protein turnover
Neuronal developement
Synaptic signaling
Modulation of chemical synaptic transmission
Neuronal structure organisation
Negative regulation of neuron apoptotic process
Response to external stimuli
Cellular homeostasis
Positive regulation of dendritic spine develoment
Vesicle-mediated transport
semantic space x
semantic space y
Log10(p-value)
Number of hit
F
Citrate cycle (TCA cycle)
Synaptic vesicle cycle
Pyruvate metabolism
Bacterial invasion of epithelial cells
Carbon metabolism
Glycolysis / Gluconeogenesis
Valine, leucine and isoleucine degradation
Endocrine and other factor−regulated calcium reabsorption
Biosynthesis of amino acids
Gap junction
GABAergic synapse
Oxidative phosphorylation
Parkinson disease
Gastric acid secretion
Insulin secretion
Retrograde endocannabinoid signaling
Huntington disease
Prion disease
Glutamatergic synapse
Diabetic cardiomyopathy
Dopaminergic synapse
Tight junction
Pathways of neurodegeneration − multiple diseases
Amyotrophic lateral sclerosis
Salmonella infectio
Chemical carcinogenesis − reactive oxygen species
Alzheimer disease
Endocytosis
Thermogenesis
Metabolic pathways
Kegg pathways
Fold enrichment
Adj p-value
7.5e−06
5.0e−06
2.5e−06
Count
50
100
150
G
WT (10m/10w)
WT (18m/10w)
Persulfidome
Proteome
H
10 we vs 10 mo
↑ PSSH = Protein
↑ PSSH ↓ Protein
↑ PSSH ↑ Protein
↓ PSSH ↓ Protein
↓ PSSH ↑ Protein
↓ PSSH = Protein
PSSH level (log2 fold change)
Protein level (log2 fold change)
I
10 we vs 18 mo
↑PSSH = Protein
↑PSSH ↓Protein
↑PSSH ↑Protein
↓PSSH ↓Protein
↓PSSH ↑Protein
↓PSSH = Protein
PSSH level (log2 fold change)
Protein level (log2 fold change)

**Supporting Figure S6. Quantiative chemoproteomic analysis of the persulfidome in aging.** (**A-B**) **Volcano plot analysis of persulfidome changes.** Volcano plot depicting statistical significance plotted against the $log_2$-fold change of persulfidated proteins in 10-month-old mice relative to 10-week-old mice (**A**) and 18-month-old mice relative to 10-week-old mice (**B**) (n = 4). Significance was established using Welch's t-test (two-sided), with a p-value threshold of < 0.05. Fold change cut-offs were established at 30%. The numbers of proteins with significantly altered persulfidation states are indicated, with the number in parentheses reflecting proteins identified through a minimum of 2 unique peptides. (**C**) Principal Component Analysis showing a notable differentiation in the persulfidome in aging as well as in the $CSE^{-/-}$ and indicating high reproducibility between replicates. (**D**) Venn diagram unraveling commonly identified proteins between our study and two other studies (*28*, *30*) using different methodologies. (**E**) GO (Biological process) term enrichment analysis of the 714 proteins gathered in clusters 1-4 (**Fig. 1F**), all showing a persulfidation decline in aging. REVIGO was used to plot the enrichment analysis performed in DAVID. Significant GO terms passed the Benjamini adjusted p-value threshold of 0.01. Circle dimensions denote the protein count within specific GO terms, while color gradients depict the degree of significance. (**F**) Kegg pathway enrichment analysis using DAVID. The graph shows the top 30 significant (Benjamini adjusted p-value < 0.01) and most enriched terms, with color gradient signifying the adjusted p-value and circle size the number of proteins. (**G**) Heatmaps representing the $log_2$-fold change of the significantly changing persulfidated proteins (Welch's test p-value < 0.05) in 10-month or 18-month-old mice compared to 10-week-old animals and their corresponding $log_2$-fold change of the protein expression level. (**H-I**) Scatter plot analysis of persulfidome and protein expression changes correlation. (**H**) Scatter plot of the log2 folds changes within the persulfidome (y-axis) and the protein expression (x-axis) for the comparison between 10-month-old and 10-week-old mice. (**I**) Parallel scatter plot analysis for the comparison between 18-month-old and 10-week-old mice. Most of the proteins undergoing persulfidation change do not surpass the significance threshold in protein expression. This observation underscores that the persulfidation changes remain largely uncoupled from concurrent changes in protein expression.

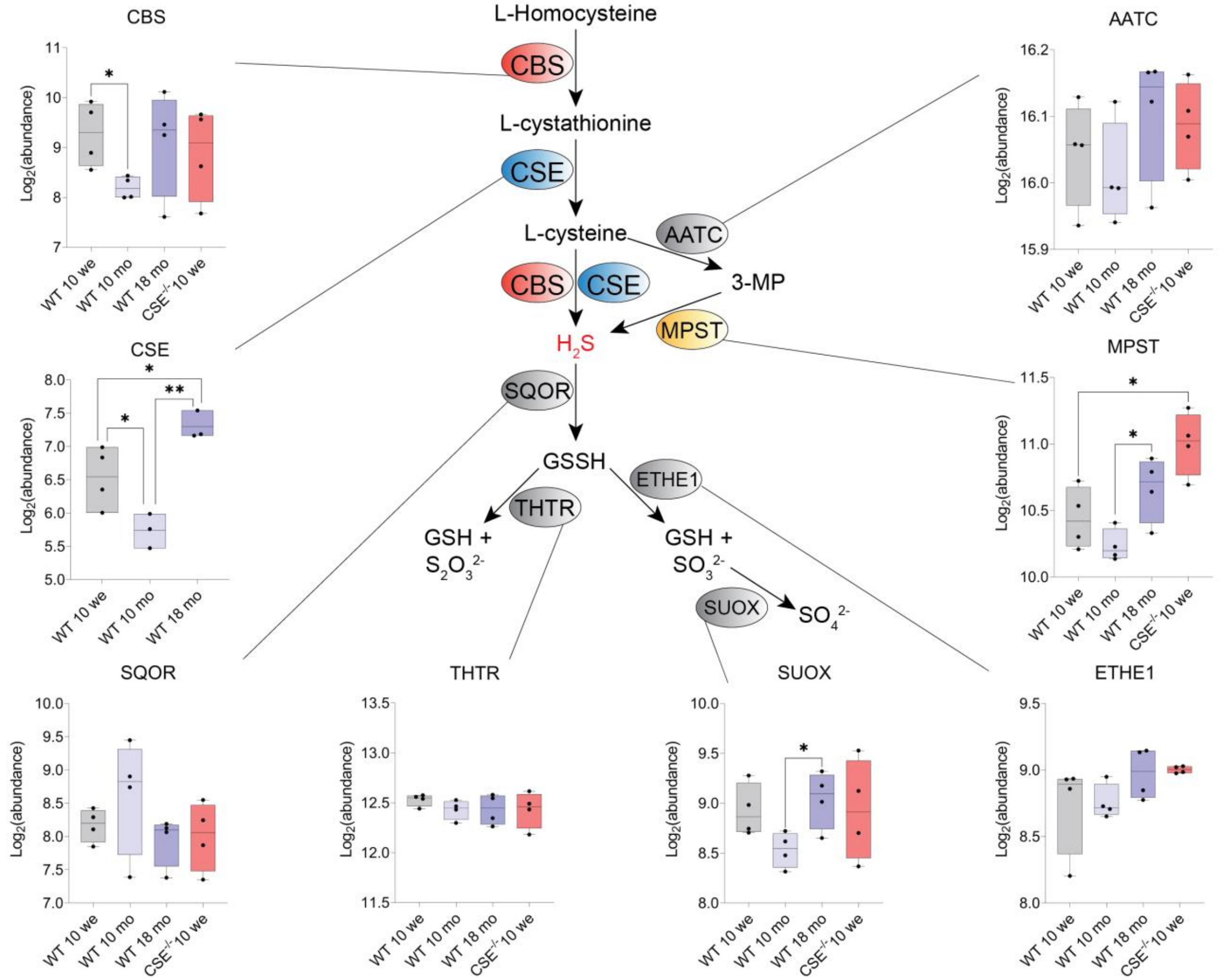


**Supporting Figure S7. Expression levels of transsulfuration pathway enzymes in aging and in CSE$^{-/-}$ mice.** Whiskers and box plots of the log2 transformed abundancies of transsulfuration pathway enzymes identified in the proteome analysis. The reported significance levels displayed with the plots directly correspond to the p-values calculated through the proteome analysis. (Welch's t-test, * < 0.05, ** < 0.01, *** < 0.001). CBS: cystathionine beta-synthase; CSE: cystathionine gamma-lyase; SQOR: sulfide quinone oxidoreductase; THTR: thiosulfate sulfurtransferase; SUOX: sulfite oxidase; ETHE1: persulfide dioxygenase ETHE1; MPST: 3-mercaptopyruvate sulfurtransferase; AATC: aspartate aminotransferase.

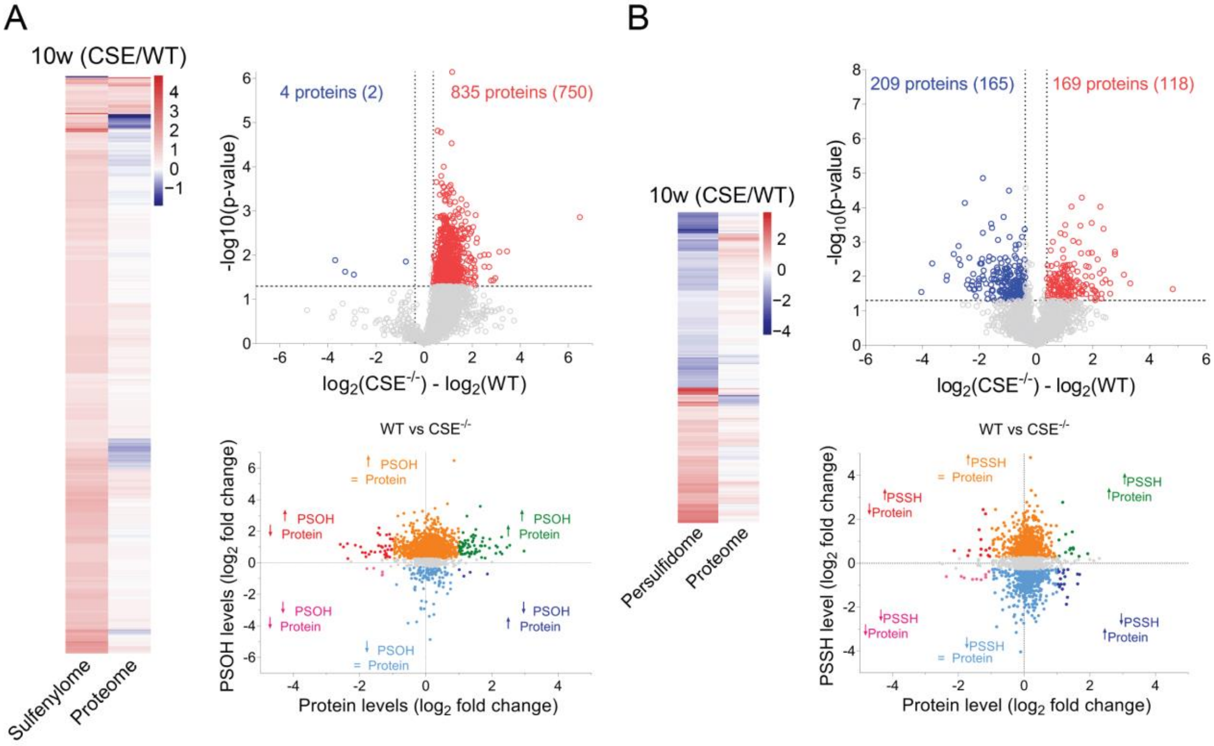


**Supporting Figure S8. Quantitative chemoproteomic analysis of the sulfenylome and persulfidome alterations in 10-week-old CSE$^{-/-}$ mice compared to WT.** (**A**) sulfenylome proteomic analysis of CSE$^{-/-}$ mice reveals higher sulfenylation levels in CSE$^{-/-}$. Top right: Volcano plot depicting statistical significance plotted against the log2-fold change of sulfenylated proteins between 10-week-old CSE$^{-/-}$ and WT mice. Significance was established using Welch's t-test (two-sided), with a p-value threshold of < 0.05. Fold change cut-offs were established at 30 %. Sulfenylated proteins exhibiting marked increases in 10-month-old mice are distinctly highlighted in red, while significantly decreased proteins are demarcated in blue. Left panel: Heatmaps representing the log2-fold change of sulfenylated proteins that undergo marked changes (Welch's test p-value < 0.05) in 10 weeks-old CSE$^{-/-}$ mice when compared to WT age-matched controls and their corresponding $log_2$-fold change of the protein expression level. Bottom right: Scatter plot of the log2 fold changes within the sulfenylation (y-axis) and the protein expression (x-axis) for the comparison between 10 weeks-old CSE-/- and WT mice. (**B**) Parallel analysis of the persulfidation changes in 10-week-old CSE$^{-/-}$ mice compared to WT mice.

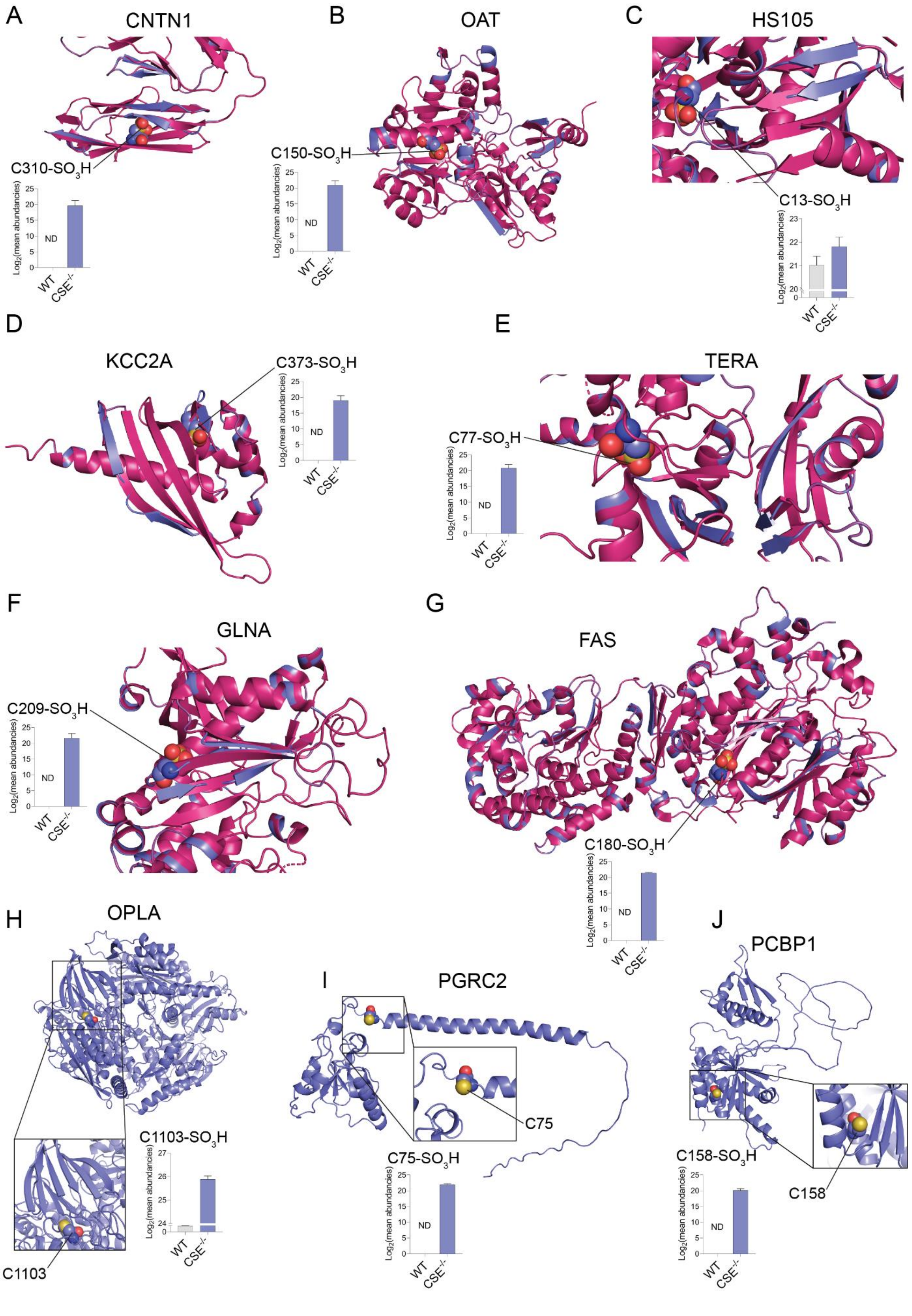
A
CNTN1
C310-SO3H
Log2(mean abundancies)
ND
WT
CSE-/-
B
OAT
C150-SO3H
C
HS105
C13-SO3H
D
KCC2A
C373-SO3H
E
TERA
C77-SO3H
F
GLNA
C209-SO3H
G
FAS
C180-SO3H
H
OPLA
C1103-SO3H
C1103
I
PGRC2
C75
C75-SO3H
J
PCBP1
C158-SO3H
C158

**Supporting Figure S9. Sulfonylation affects the structure of proteins.** Bar graph represents the $log_2$ (mean abundancies) of the corresponding sulfonylated peptide in 10 weeks-old WT and $CSE^{-/-}$ (n = 4, ND: Non-defined). (**A-G**) Prediction of the structure changes upon cysteine sulfonylation found to increase in $CSE^{-/-}$ mice using VIENNA-PTM. The original crystal structure is depicted in pink, and the predicted model in blue. (**A**) Contactin-1 (CNTN1) based on the 3S97.pdb model. (**B**) Ornithine aminotransferase (OAT) is based on the 2OAT.pdb model. (**C**) Heat shock protein 105 kDa (HS105) is based on the 6GFA.pdb model. (**D)** Calcium/calmodulin-dependent protein kinase type II subunit alpha (KCC2A) is based on the 1HKX.pdb model. (**E)** Transitional endoplasmic reticulum ATPase (TERA) is based on the 1R7R.pdb model. (**F**) Glutamine synthetase (GLNA) is based on the 2OJW.pdb model**.** (**G**) Fatty acid synthase (FAS) is based on 5MY2.pdb model (**H-J**) Alphafold prediction model of the proteins identifed to undergo cysteine sulfonylation in $CSE^{-/-}$. (**H**) 5-oxoprolinase (OPLA) is based on the AF-O14841-F1-model_v4.pdb model. (**I**) Intercellular adhesion molecule 5 (ICAM5) is based on the AF-Q9UMF0-F1-model_v4.pdb model. (**J**) Poly(rC)-binding protein 1 (PCBP1) is based on the AF-Q15365-F1-model_v4.pdb model.

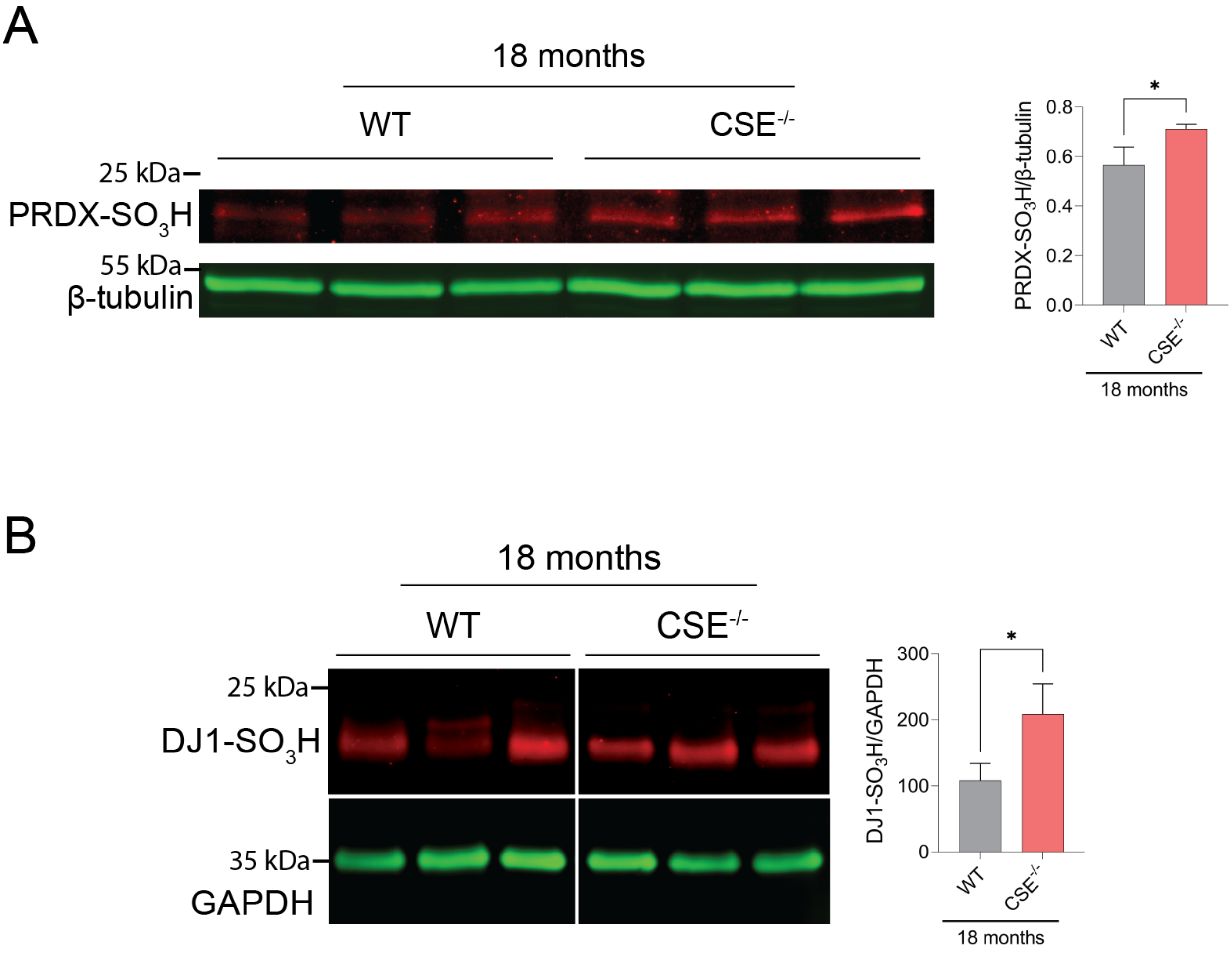


**Supporting Figure S10. Enhanced sulfonylation of PRDX and DJ1 in aged CSE$^{-/-}$ mice** (**A**) PRDX over-oxidation levels in 18-month-old WT and CSE$^{-/-}$ mice. The intensities were normalized to the β-tubulin expression levels. Values represent mean ± SD (Two-sided t-test,* $p < 0.05$, n = 3). (**B**) DJ1 (PARK7) over-oxidation levels in 18-months-old WT and CSE$^{-/-}$ mice. The intensities were normalized to the GAPDH expression levels. Values represent mean ± SD (Two-sided t-test,* $p < 0.05$, n = 3).

A

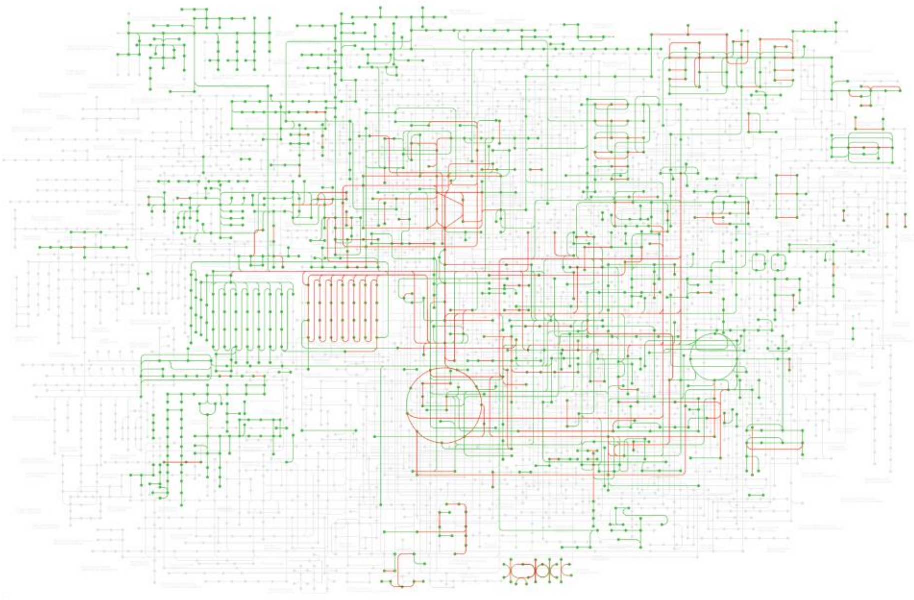

B

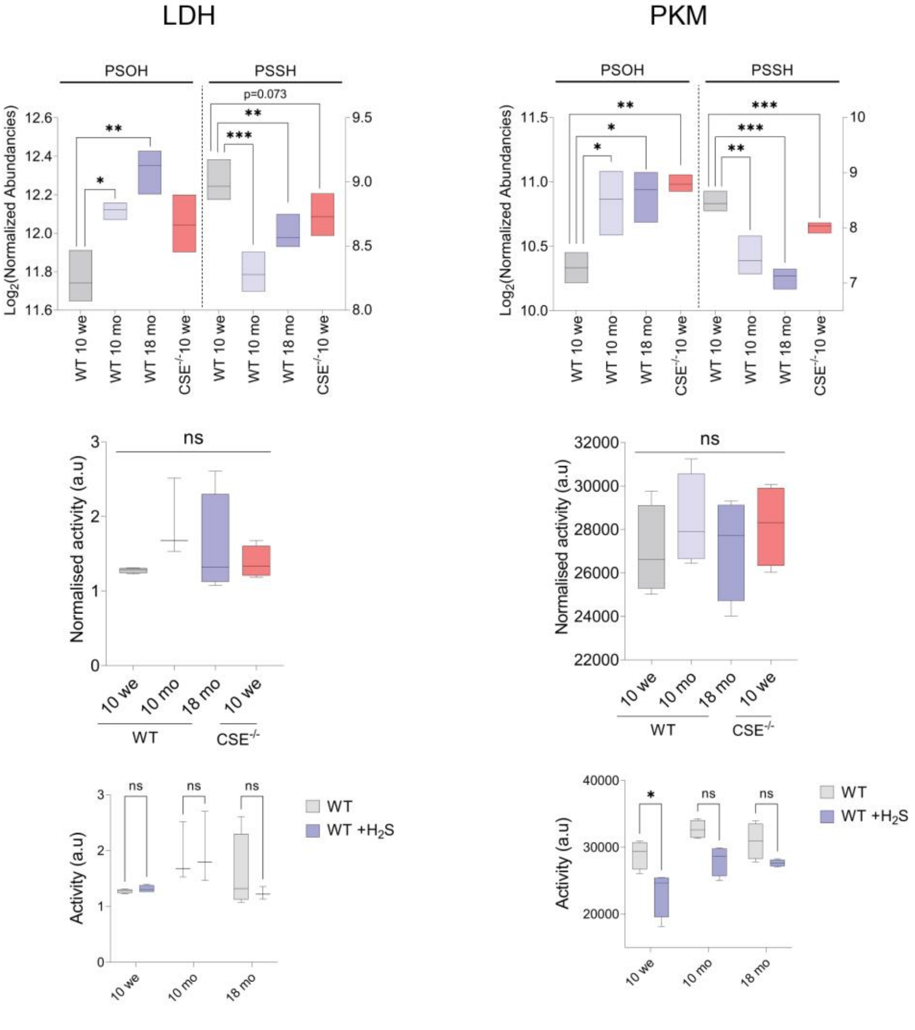
LDH
PKM
PSOH
PSSH
p=0.073
Log2(Normalized Abundancies)
WT 10 we
WT 10 mo
WT 18 mo
CSE-/- 10 we
ns
Normalised activity (a.u)
10 we
10 mo
18 mo
WT
CSE-/-
Activity (a.u)
WT
WT +H2S

**Supporting Figure S11. Age-induced sulfenylation and persulfidation changes of metabolic enzymes exhibit no impact on their activity.** (**A**) Kegg metabolic pathways map with the red lines connecting proteins whose persulfidation shows substantial alterations during the aging process. (**B**) Lactate dehydrogenase (LDH) and pyruvate kinase (PKM), both key metabolic enzymes, show significant changes in sulfenylation and persulfidation in aging and in 10-week-old CSE$^{-/-}$ mice but no change could be observed in their activity. Top Panel: Box plots of the $\log_2$ transformed abundancies for the sulfenylation and persulfidation levels in aging and in CSE$^{-/-}$ mice. The reported significance levels displayed with the plots directly correspond to the p-values calculated through the proteome analysis. (Welch's t-test, * < 0.05, ** < 0.01, *** < 0.001). Middle panel: The normalized activity (a.u.) of LDH and PKM in brain lysates is shown as a whisker and box plot. The activity levels of these enzymes remain unaltered across the tested conditions (2-way ANOVA and Tukey's HSD Ttst for multiple comparisons, ns: non-significant). Bottom panel: The enzymatic activity was also assessed in the brain lysates supplemented with $H_2S$. Significance was tested using 2-way ANOVA and Tukey's HSD test for multiple comparisons (ns: non-significant, * < 0.05).

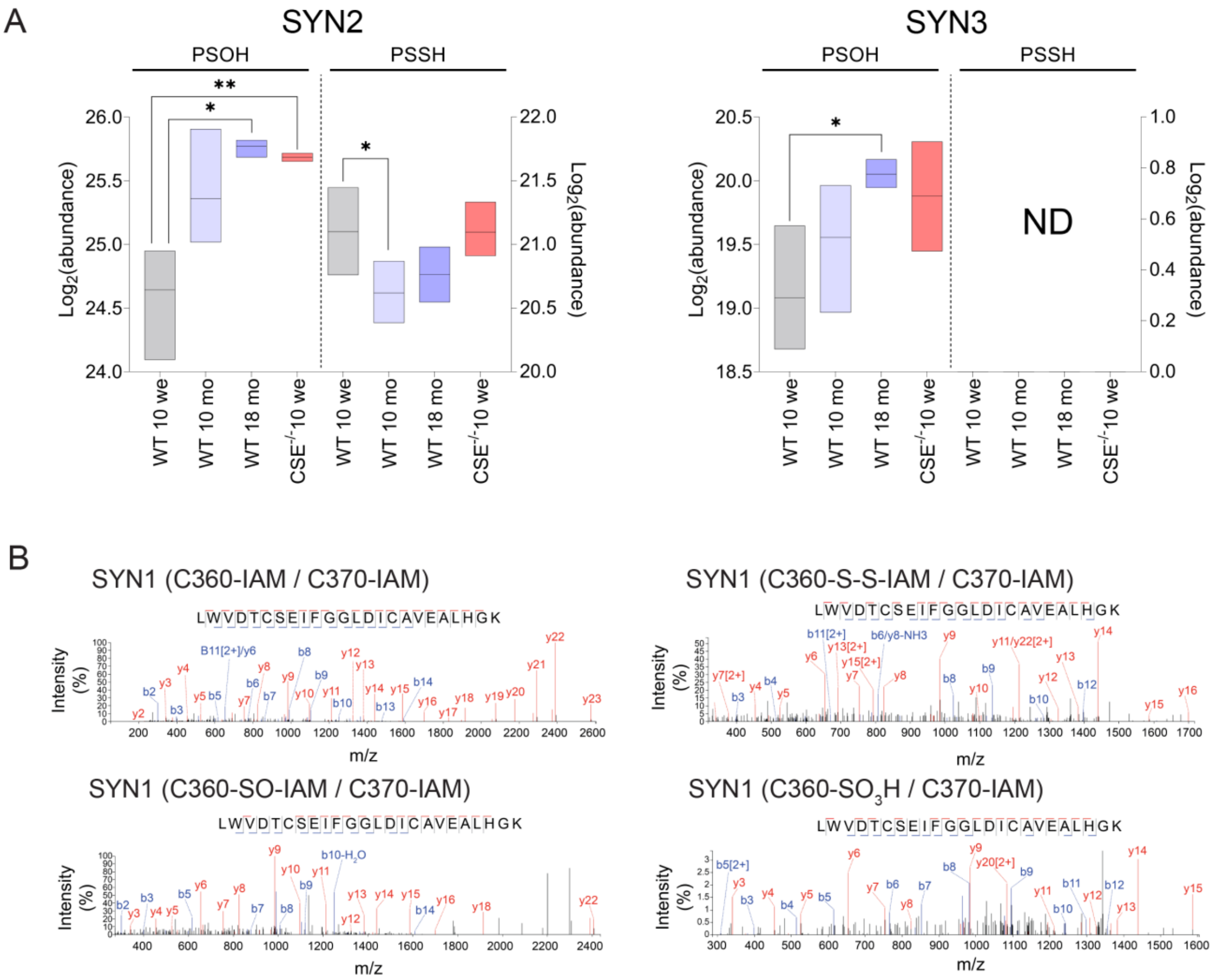


**Supporting Figure S12. Age-induced changes in synapsin's thiol oxidation** (**A**) Boxplot of the age-induced sulfenylation (PSOH) and persulfidation (PSSH) of Synapsin-2 and -3 (Welch's test, p-value: * <0.05, **<0.01, ***<0.001). (**B**) Annotated TimsTOF CID MS/MS spectra of synapsin-1 peptides highlighting distinct oxidation states of cysteine 360. Spectra originate from quantified peptides extracted from WT and CSE$^{-/-}$ 18-month-old mice, as depicted in Figure 2B. The y-ions are denoted in red, while the b-ions are indicated in blue.

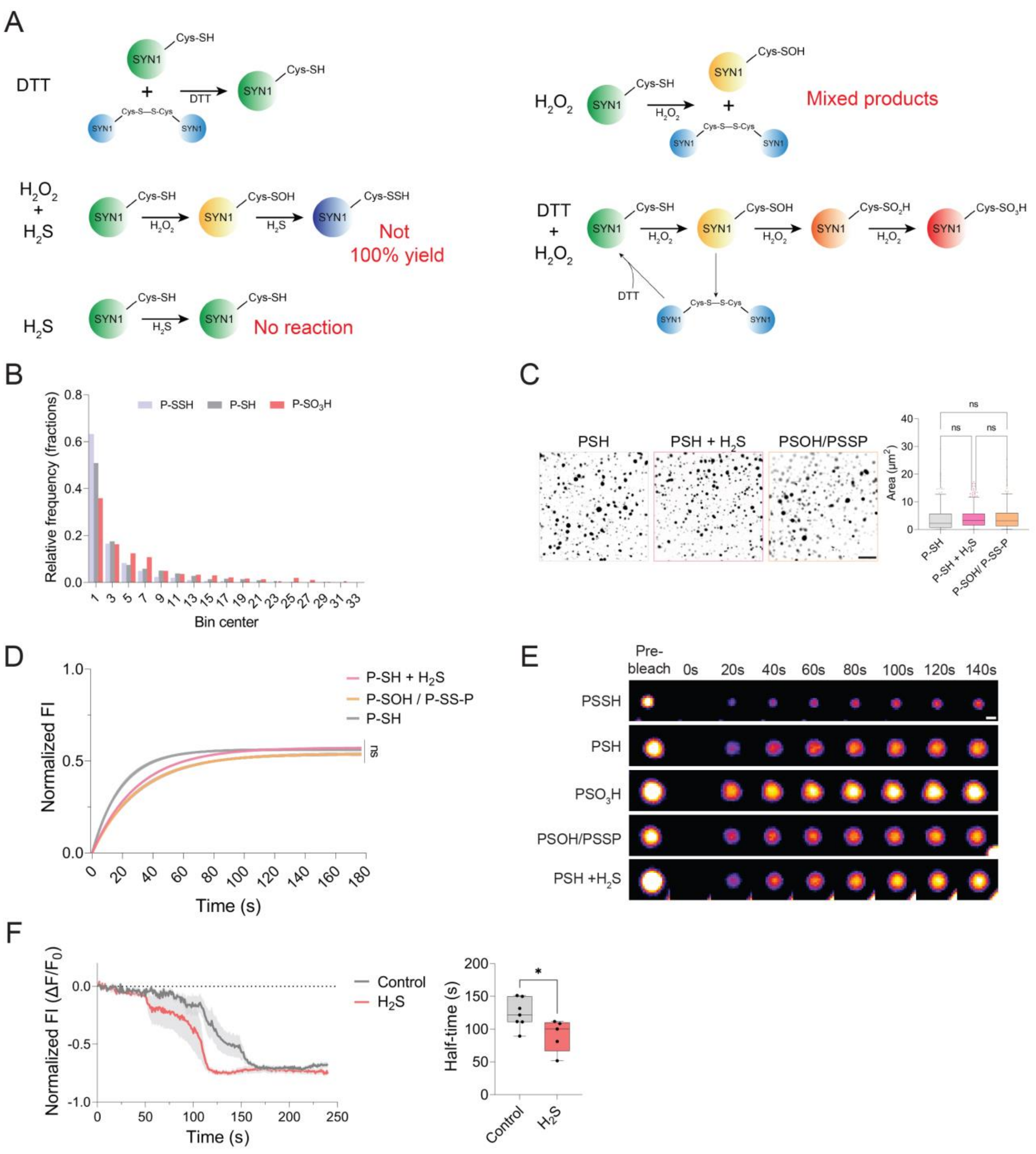


**Supporting Figure S13. Synapsin-1 phase separation is modulated through cysteine modifications.** (**A**) Schematic representation of the protocols for preparation of different cysteine modifications, used for the phase separation and FRAP experiments. PSH (100 µM DTT), PSSH (100 µM $H_2O_2$ + 500 µM $H_2S$), and $PSO_3H$ (100 µM DTT + 200 µM $H_2O_2$), PSOH/PSSP (200 µM $H_2O_2$). DTT reduced EFGP-synapsin-1 was also treated with 500 µM $H_2S$ as a negative control. (**B**) Relative frequency histogram of EFGP-synapsin-1 droplets in relation to their size. (**C**) Representative microscopy analysis of PSH (100 µM DTT), PSH + $H_2S$ (500 µM), and PSOH/PSSP (200 µM $H_2O_2$) EFGP-synapsin-1 droplets obtained 10 minutes after phase separation was initiated using 3%

PEG3000. Quantitative assessment of droplet area (µm²) is presented on the right (1-way ANOVA, Tukey's HSD test for multiple comparisons, ns: non-significant). Scale bar = 20 µm. (**D**) Quantification of fluorescence recovery after photobleaching. Prior to initiating phase separation, 10 µM EFGP-synapsin-1 samples underwent treatment with various cysteine-modifying agents as described in Material and Methods section. Subsequent phase separation was induced by 3% PEG3000. Recovery profiles are depicted by mean kinetic lines. Significance was assessed by 2-way ANOVA followed by the Šidák test for multiple comparisons, denoted as "ns" for non-significant. (**E**) Microscopy visualization of FRAP shown in (D) and Fig. 2E**.** Representative microscopy images capturing EGFP-synapsin-1 condensates before and after photobleaching. Scale bar = 1 µm. (**F**) Kinetics of mScarlet-synapsin-1 droplet dissolution in the synapse of primary neurons upon stimulation with 92.5 mM KCl ($n \geq 3$).

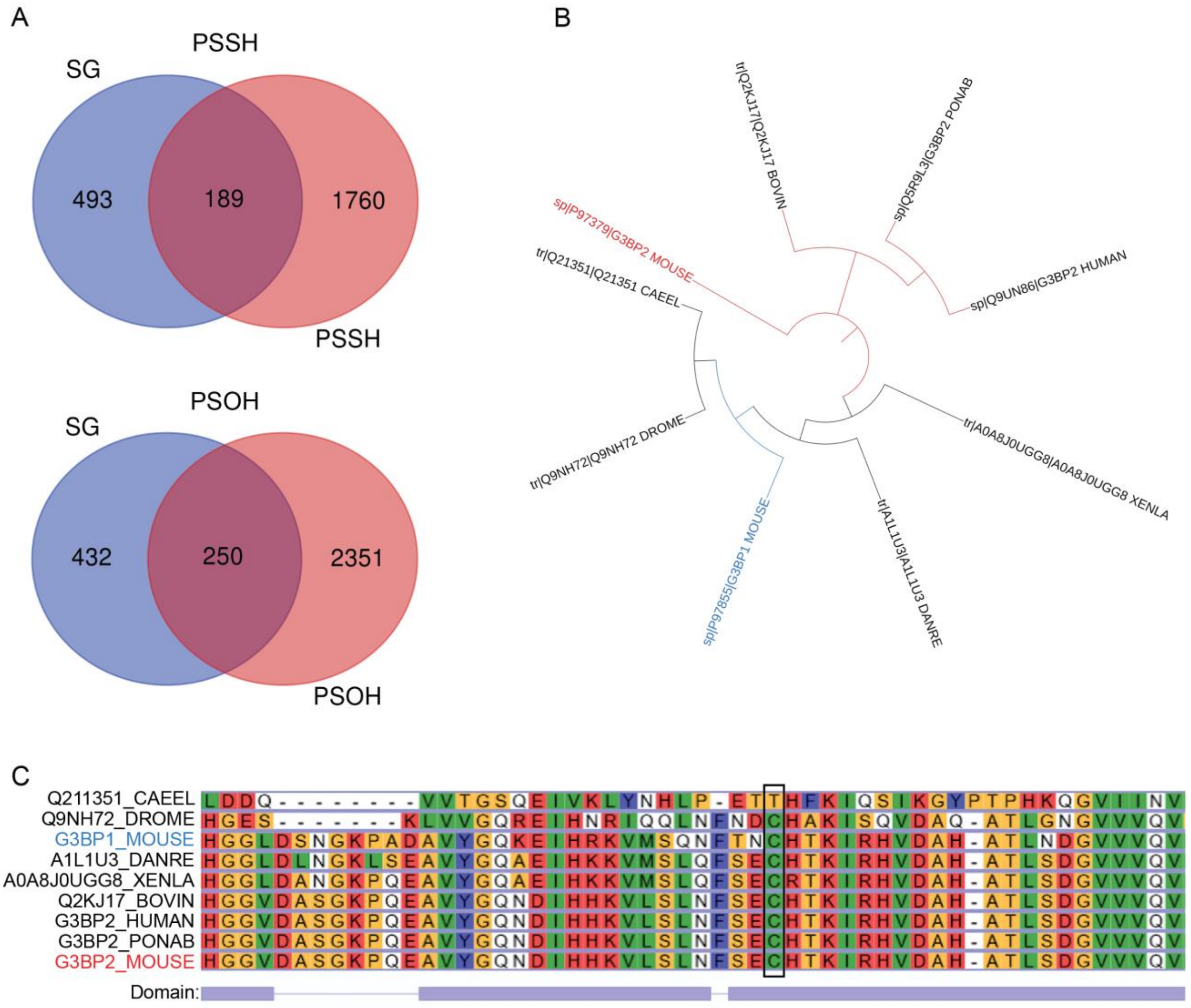


**Supporting Figure S14. Age-induced remodeling of persulfidome and sulfenylome affects stress granule proteins.** (**A**) Venn diagram representing the overlap of proteins present in stress granules (SG) and those identified in the persulfidome (PSSH) or sulfenylome (PSOH). (**B**) Circular phylogenetic cladogram build using G2BP2 sequences from a range of species: *Mus musculus* (MOUSE), *Bos taurus* (BOVIN), *Pongo abelii* (PONAB), *Homo sapiens* (Human), *Xenopus laevis* (XENA), *Danio rerio* (DANRE), *Drosophila melanogaster* (DROME), and *Caenorhabditis elegans* (CAEEL). For contextual comparison, G3BP1 from *Mus musculus* is also included. Of note, by increasing the FDR to 5 % we could observe G3BP1 in our persulfidome and sulfenylome analysis, suggesting that in other tissues where expression of G3BP1 is predominant, the same cysteine PTMs could occur. (**C**) Alignment of G3BP2 sequences across various species using the Uniprot alignment tool. The conserved cysteine residue, notably positioned at C73 in the mouse G3BP2 sequence, is highlighted with a black rectangle. Conservation of this cysteine across species underlines its potential important role in the protein functions such as its phase separation potential.

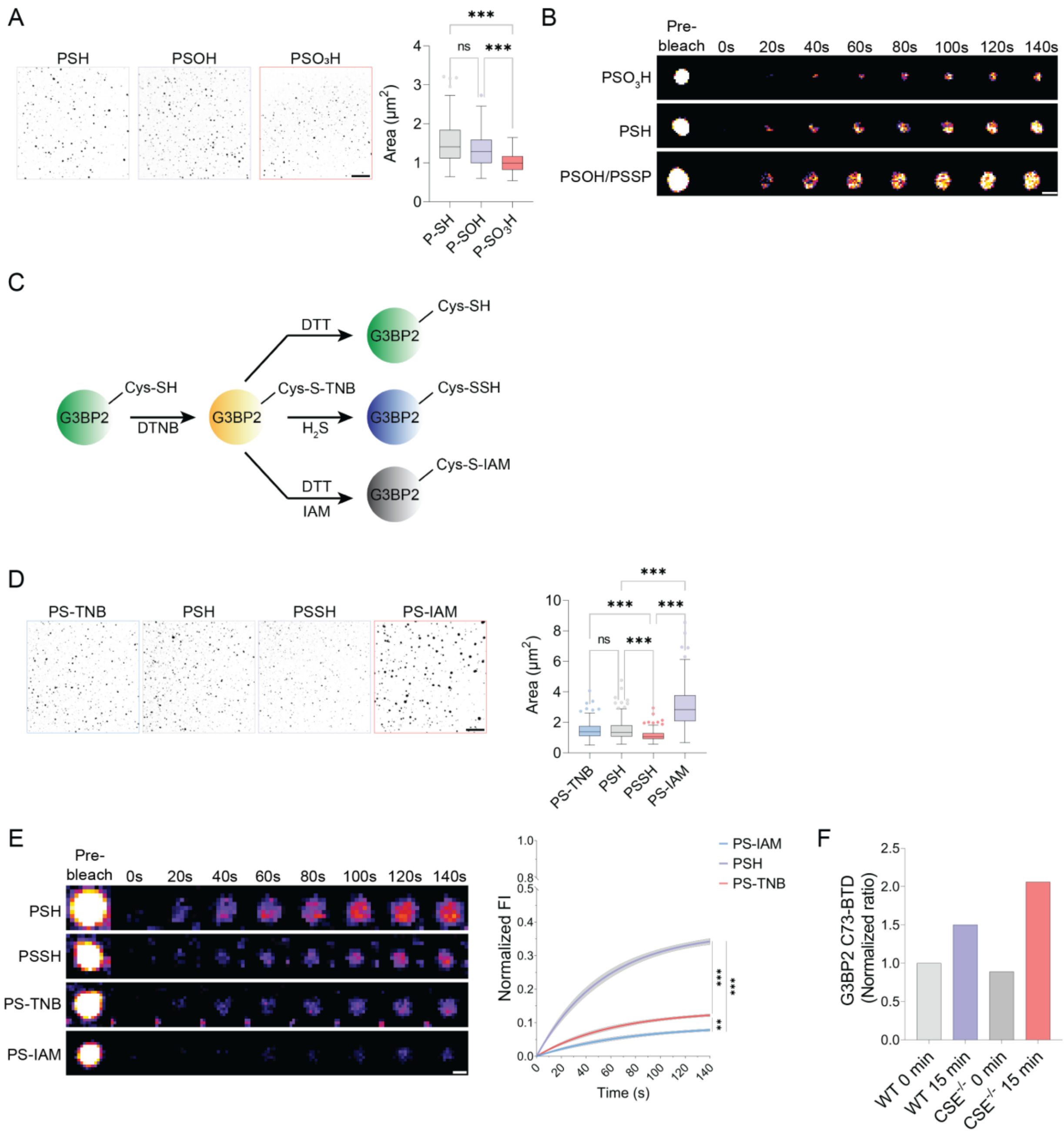


**Supporting Figure S15. Cysteine-mediated regulation of G3BP2 phase separation.** (**A**) Representative G3BP2 droplets images. Prior to phase separation induction, G3BP2 (10 µM) underwent treatment with DTT (100 µM; PSH), $H_2O_2$ (200 µM; PSOH), or a combination of DTT and $H_2O_2$ (100 µM and 200 µM respectively, P-$SO_3H$), for 30 min at 37 °C. LLPS was induced using 50 ng/µl of total RNA extract, and images were taken 5 min after induction. Quantification of droplet area shows a significant reduction in size for sulfonylated G3BP2 (1-way ANOVA, Tukey's HSD Test for multiple comparisons; ***<0.001). Scale bar = 20 µm. (**B**) Representative FRAP images depicting mobility dynamics of G3BP2 droplets under PSH, PSOH, and $PSO_3H$ conditions. Quantification of the FRAP data is presented in Fig. 3C. Treatment conditions were as mentioned above and FRAP was started 7 min after the addition of RNA. Scale bar = 2 µm. (**C**) Schematic illustration detailing G3BP2 treatment resulting in various cysteine modifications: PS-

TNB, PSSH, PSH, and PS-IAM. (**D**) Representative images of G3BP2 phase condensates following 5 min of LLPS induction for PS-TNB, PSH, PSSH, and PS-IAM conditions. G3BP2 modified with Ellman's reagent (PS-TNB) was treated with DTT (20 µM, to give PSH), $H_2S$ (20 µM, to give PSSH), or DTT (20 µM) first then iodoacetamide (200 µM, to give PS-IAM), for 5 min at 37 °C. Scale bar = 20 µm. Quantitative droplet area analysis, depicted as box plot and whiskers, demonstrates a significant reduction in PSSH G3BP2 droplet size, while PS-IAM treatment notably increased droplet area (1-way ANOVA, Tukey's HSD Test for multiple comparisons; ***<0.001). (**E**) Representative FRAP images for different thiol-modified G3BP2 condensates, accompanied by quantification of normalized fluorescence intensity over time. Recovery profiles are depicted by mean lines with standard errors shown in gray shade (2-way ANOVA, Šidák test for multiple comparisons; **<0.01, ***<0.001). Scale bar = 2 µm. (**F**) BTD-trapping of G3BP2 sulfenylation in MEF wild type (WT) and $CSE^{-/-}$ cells treated with 500 µM arsenite for indicated time. G3BP2 was immuno-pulled down and protein analyzed by MS for BTD-tagged C37-containing peptide. n = 3.

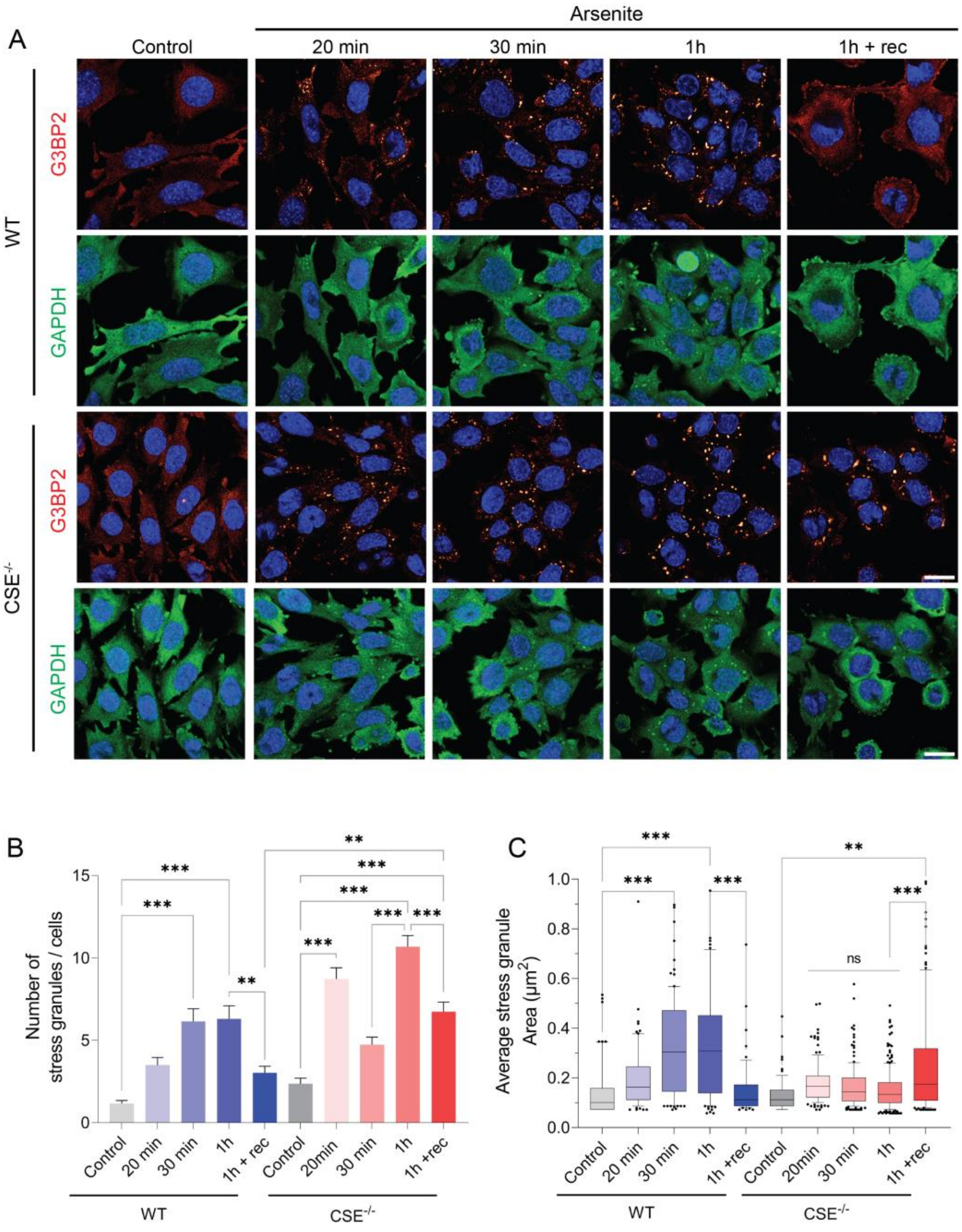


**Supporting Figure S16. Time-resolved G3BP2 and GAPDH condensates formation in MEF cells** (**A**) Representative confocal images of G3BP2 (dynamic red) and GAPDH (green) immunocytochemical staining upon 500 µM $NaAsO_2$ stress over time in MEF WT and $CSE^{-/-}$. All experiments were performed in at least triplicates with several images analyzed per replicate. Scale bar = 20 µm. (**B-C**) Quantitative analysis of G3BP2 stress granules. Quantification of the number (**B**) and size (in µm²) (**C**) of G3BP2 stress granules. Notably, cells treated for 1 hour and allowed to recover for an additional hour (1h + rec) exhibit reduction of stress granules in WT MEFs. In contrast, $CSE^{-/-}$ MEF maintain the presence of stress granules, which are further characterized by increased size (1-way ANOVA, Tukey's HSD test for multiple comparisons; *<0.05, **<0.01, ***<0.001).

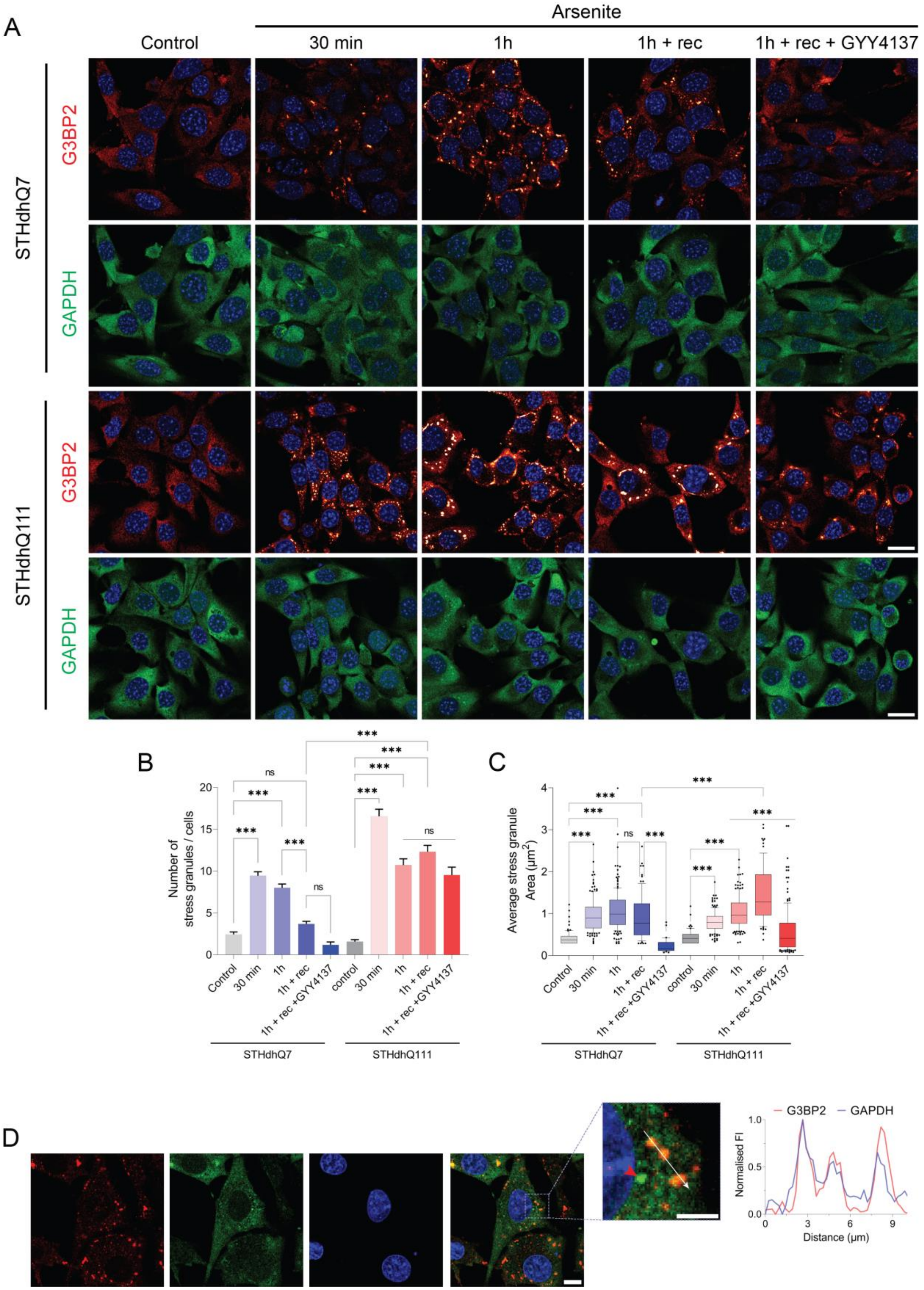

A
Arsenite
Control
30 min
1h
1h + rec
1h + rec + GYY4137
STHdhQ7
G3BP2
GAPDH
STHdhQ111
B
Number of stress granules / cells
Control
30 min
1h
1h + rec
1h + rec +GYY4137
control
STHdhQ7
STHdhQ111
C
Average stress granule Area (µm²)
D
G3BP2
GAPDH
Normalised FI
Distance (µm)

**Supporting Figure S17. Time-resolved G3BP2 and GAPDH condensates formation in STHdhQ111 cells.** (**A**) Representative confocal images of G3BP2 (dynamic red) and GAPDH (green) immunocytochemical staining upon 500 µM $NaAsO_2$ stress over time in STHdhQ7 and STHdhQ111, which are known to have diminished expression of CSE. Scale bar = 20 µm. (**B-C**) Quantitative analysis of G3BP2 stress granules. Quantification of the number (**B**) and size (in $\mu m^2$) (**C**) of G3BP2 stress granules. Notably, STHdhQ111 cells exhibited higher number of G3BP2-induced stress granules. Additionally, cells treated for 1 hour and allowed to recover for an additional hour (1h + rec) displayed a significant decrease in G3BP2 stress granules in STHdhQ7 cells, whereas STHdhQ111 cells maintained a persistent number of stress granules along with an increase in their size. The addition of GYY4137 (a slow-releasing $H_2S$ donor) during the recovery period resulted in a significant reduction in G3BP2 granule size in both cell lines. (1-way ANOVA, Tukey's HSD Test for multiple comparisons; *<0.05, **<0.01, ***<0.001). (**D**) Co-localization analysis of GAPDH (green) and G3BP2 (red) condensates in STHdh cell line upon $NaAsO_2$ treatment (1h) (representative confocal images). Fluorescence intensity profiling was performed along the line demarcated by the white arrow. Of note, isolated GAPDH condensates were also observed (red arrowhead), implying that the formation of GAPDH condensates could be independent from G3BP2 stress granules. Scale bar = 10 µm and in the zoomed intercept 5 µm.

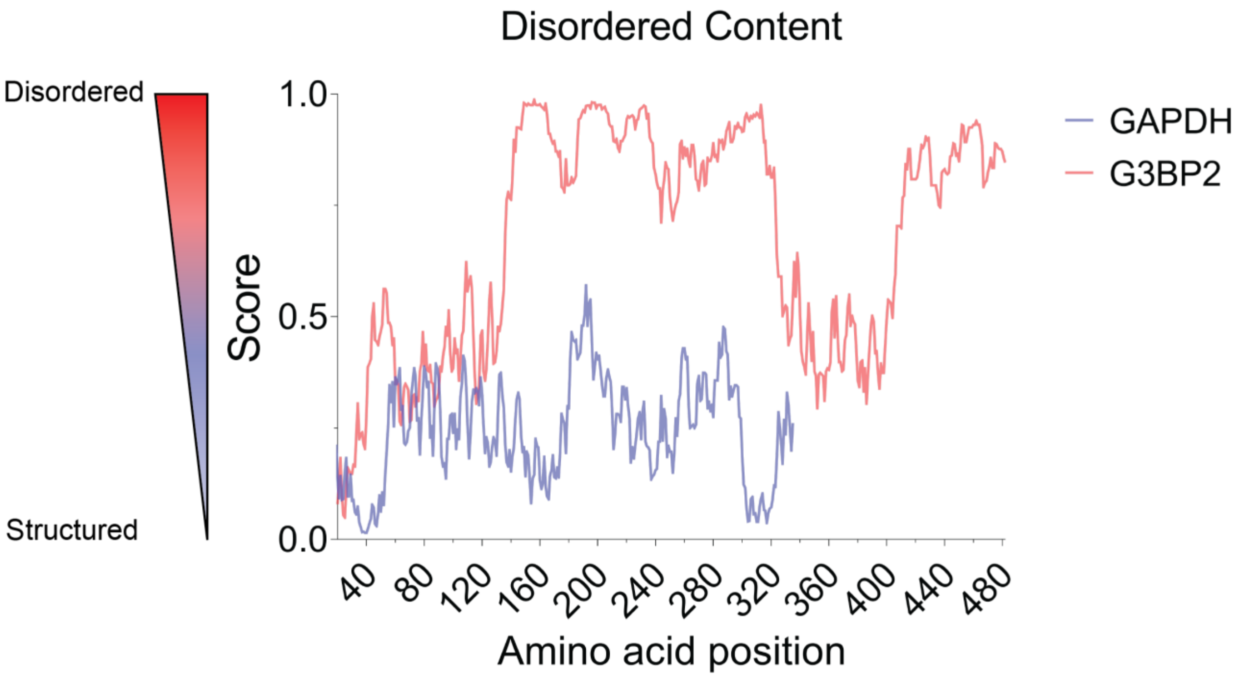


**Supporting Figure S18. Comparison of disordered content distribution in the structures of human GAPDH and G3BP2 obtained by CD-Code.**

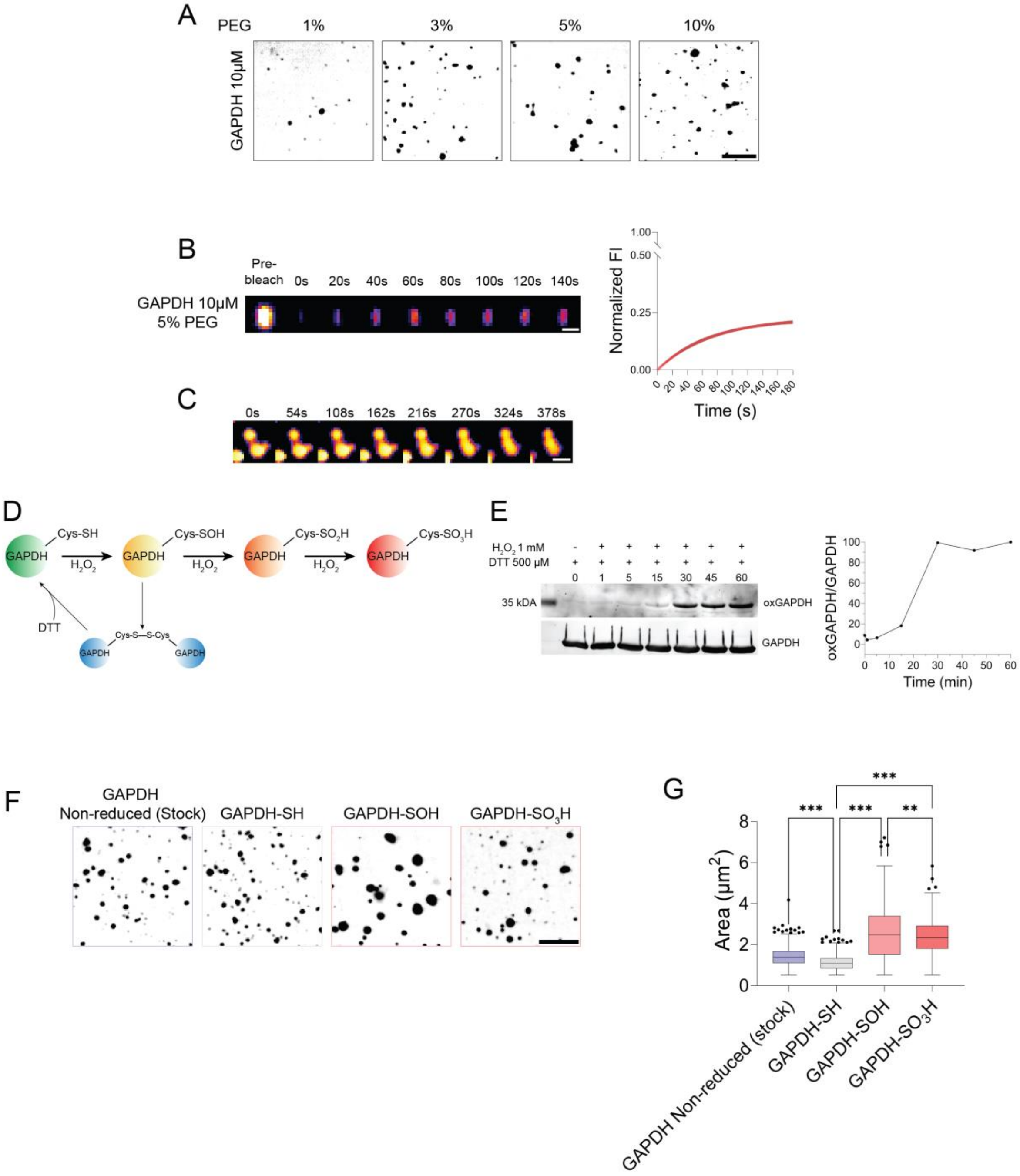


**Supporting Figure S19. GAPDH, a highly structured protein, is capable of liquid-liquid phase separation.** (**A**) Representative images of GAPDH (10 µM) liquid-liquid phase separations (LLPS) initiated through 10-minute incubation with different PEG3000 concentrations. Scale bar = 10 µm. (**B**) Representative microscopy visualization of FRAP assay on GAPDH condensates induced with 5% PEG3000**.** Quantitative evaluation of the FRAP experiment further underscores the recovery behavior of GAPDH droplets. Mean line depict recovery profile with standard errors shown in gray shades. Scale bar = 2 µm. (**C**) Liquid droplet-like properties of phase-separated GAPDH are exemplified by the depicted fusion event. Scale bar = 2 µm. (**D**) Schematic representation of protocol for preparing sulfonylated GAPDH. (**E**) GAPDH sulfonylation yield assessment through Western blot analysis using anti-GAPDH-$SO_3$H antibody. Quantitative analysis indicates the relative proportion of sulfonylated GAPDH normalized to the GAPDH loading control. GAPDH (50 µM) underwent treatment with DTT (500 µM) and $H_2O_2$ (1 mM) over varying time durations. Maximum over-oxidation yield was achieved after 30 minutes of treatment, a duration that was subsequently adopted for all further experiments. (**F**) Representative microscopy images of non-treated (GAPDH), fully reduced, $H_2O_2$-oxidized and sulfonylated GAPDH droplets after 5 min incubation with 30 ng/ul of total RNA extract. Prior to induction, GAPDH (10 µM) was either untreated (GAPDH) or treated with DTT (100 µM; reduced) or $H_2O_2$ (200 µM; oxidized) for 5 minutes at 37 °C. Sulfonylated GAPDH was prepared as described above. Scale bar = 10 µm. (**G**) Quantification of droplet area obtained for different cysteine oxidation states of GAPDH, prepared as described above (1-way ANOVA, tukey's HSD Test for multiple comparisons; *<0.05, **<0.01, ***<0.001).

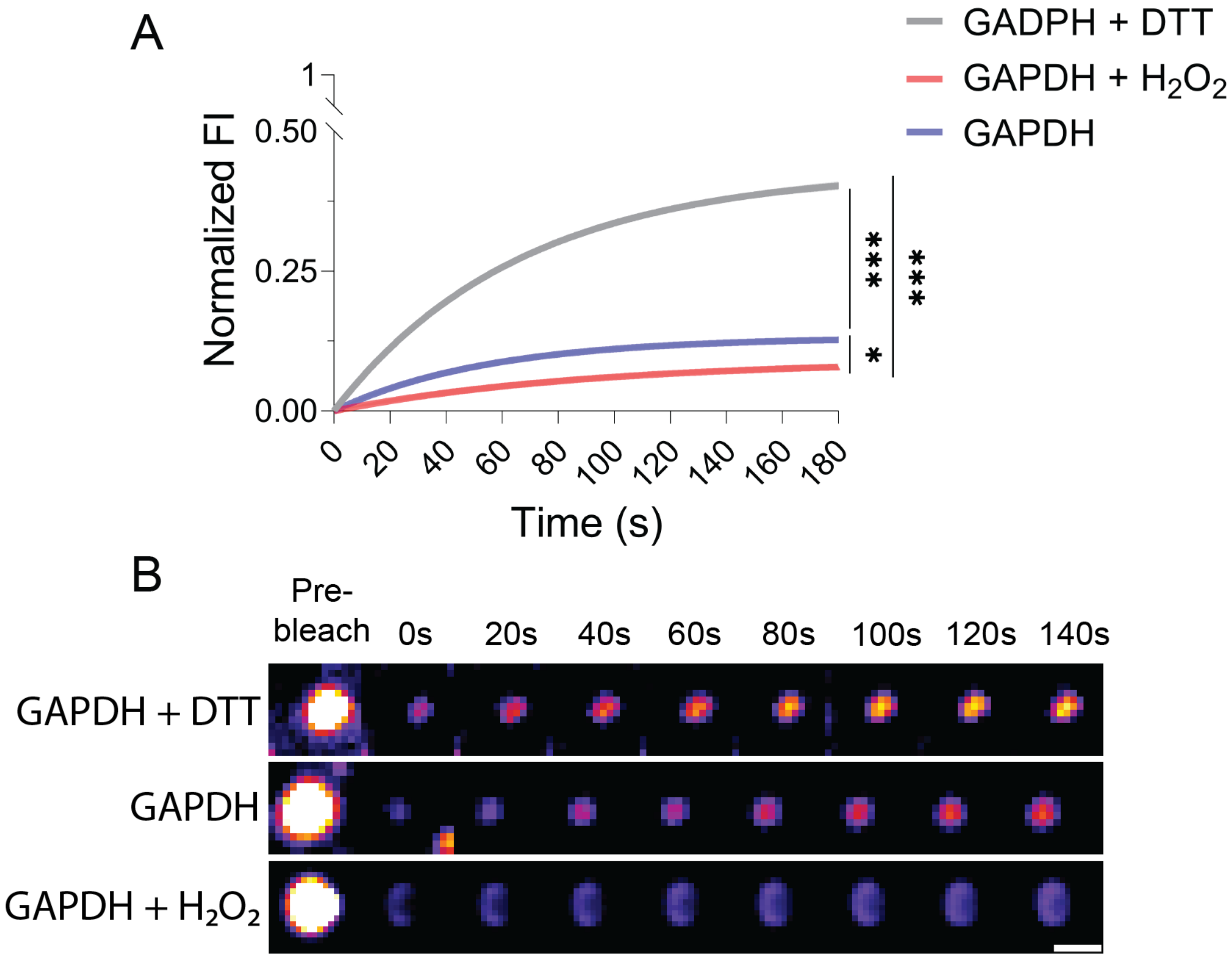


**Supporting Figure S20. FRAP analysis of GAPDH condensates.** (**A**) Quantitative evaluation of the FRAP assay performed on distinct forms of GAPDH: non-treated, reduced and oxidized. Prior to induction, human recombinant GAPDH (10 µM) was either kept as it is (GAPDH) or treated with DTT (100 µM; reduced) or $H_2O_2$ (200 µM; oxidized) for 5 minutes at 37°. Photobleaching was performed 5 min after the induction of phase separation with 30 ng/µl of RNA. Mean lines depict recovery profile (2-way ANOVA, Šidák test for multiple comparisons; **<0.01, ***<0.001). (**B**) Representative microscopy images depicting the FRAP assay of GAPDH condensates formed by different thiol modifications. Scale bar = 2 µm.

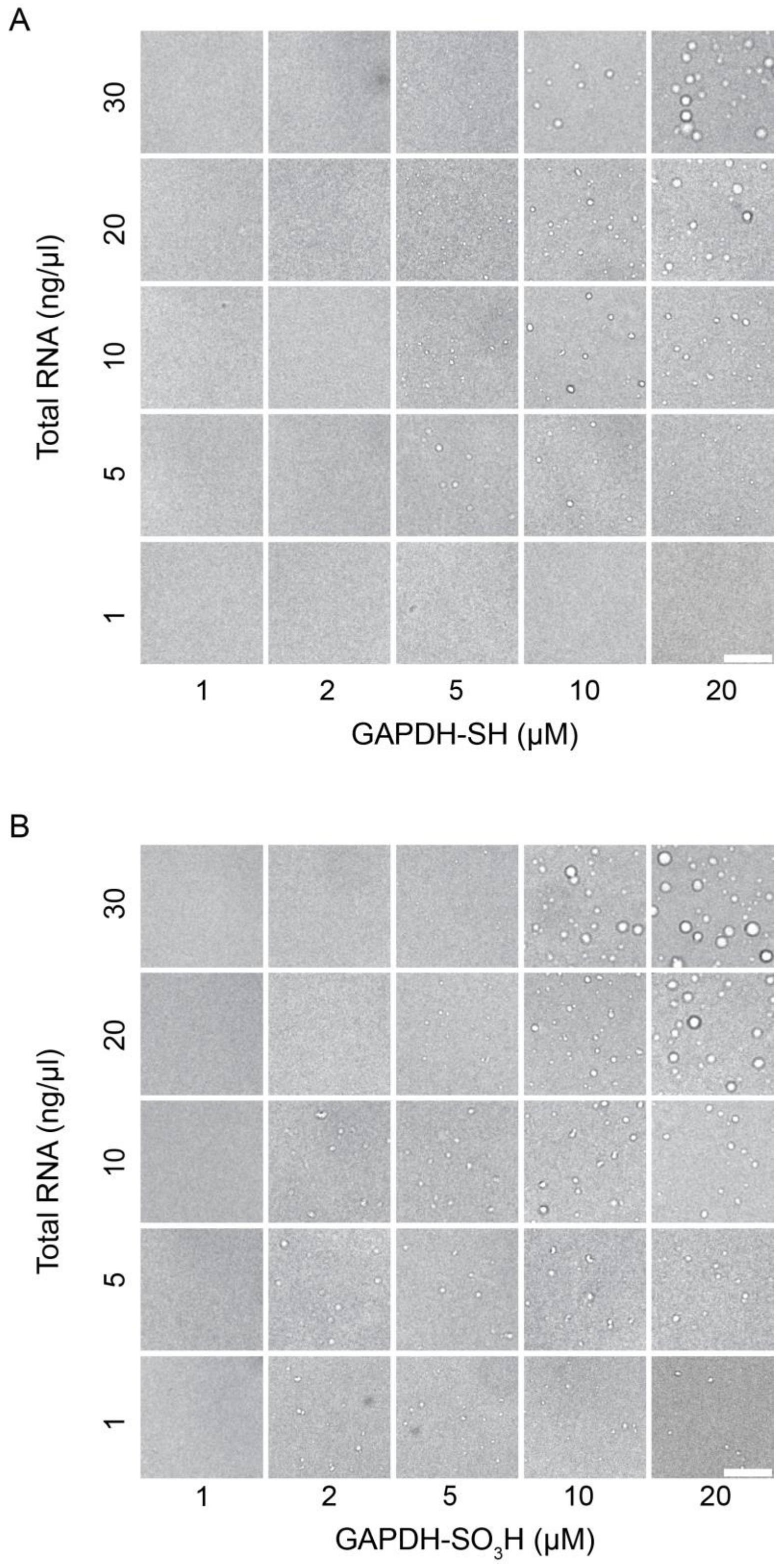


**Supporting Figure S21. Phase diagram of GAPDH LLPS.** (**A-B**) Representative images of the LLPS phase diagram formed with (**A**) the reduced or (**B**) sulfonylated GAPDH. The phase diagram was built using increased concentrations of both the protein and total RNA extract. Scale bar = 10 µm.

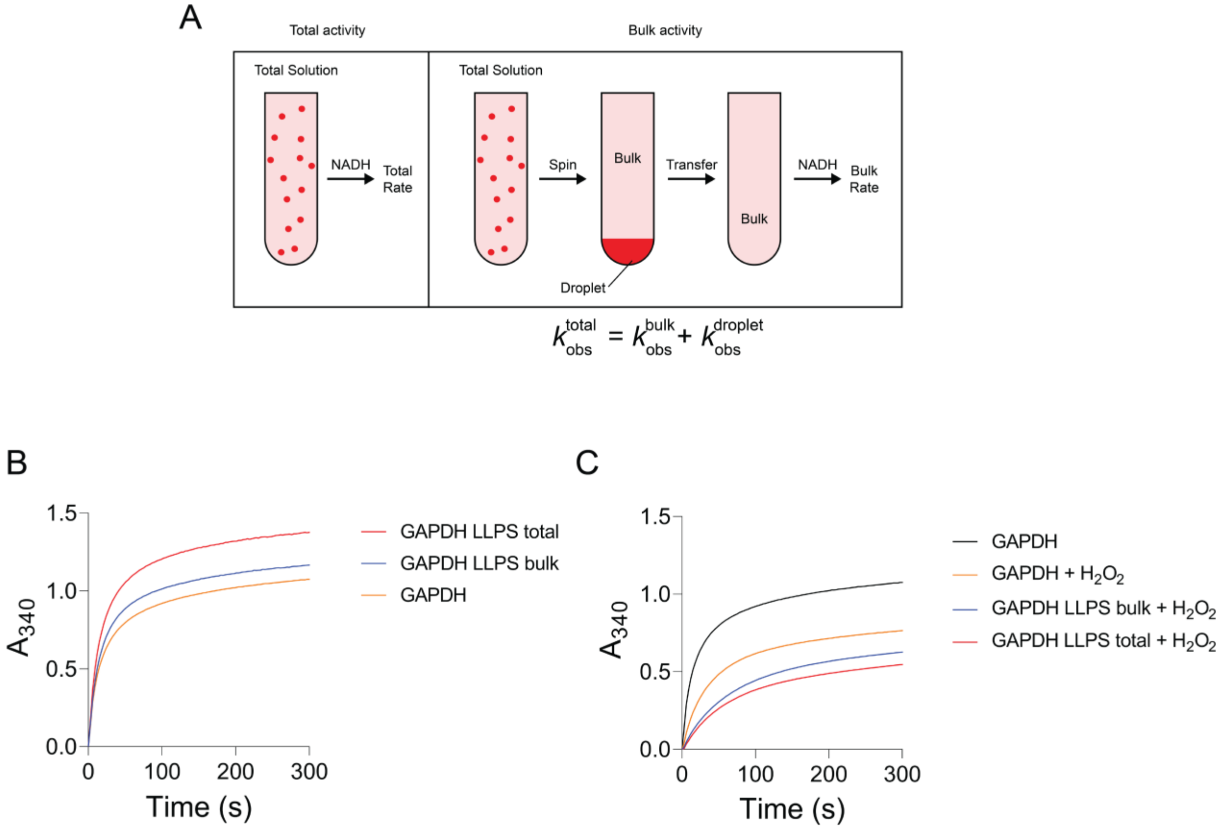


**Supporting Figure S22. GAPDH LLPS impact the protein activity and reactivity with $H_2O_2$.** (**A**) Depiction of the experimental design used to assess the effect of LLPS of GAPDH enzymatic activity. Modified from Peeples et al, 2021 (*74*). (**B**-**C**) Kinetics of GAPDH (10 µM) activity in solution (in absence of phase separation), in the bulk fraction, and within the total LLPS fraction (induced by the addition of 30 ng/µL of total RNA). Activity measurements were performed in phase separation buffer (see Material and Methods) (**B**) in the presence of 2 mM sodium bicarbonate and (**C**) in the presence of 2 mM sodium bicarbonate and 100 µM $H_2O_2$.

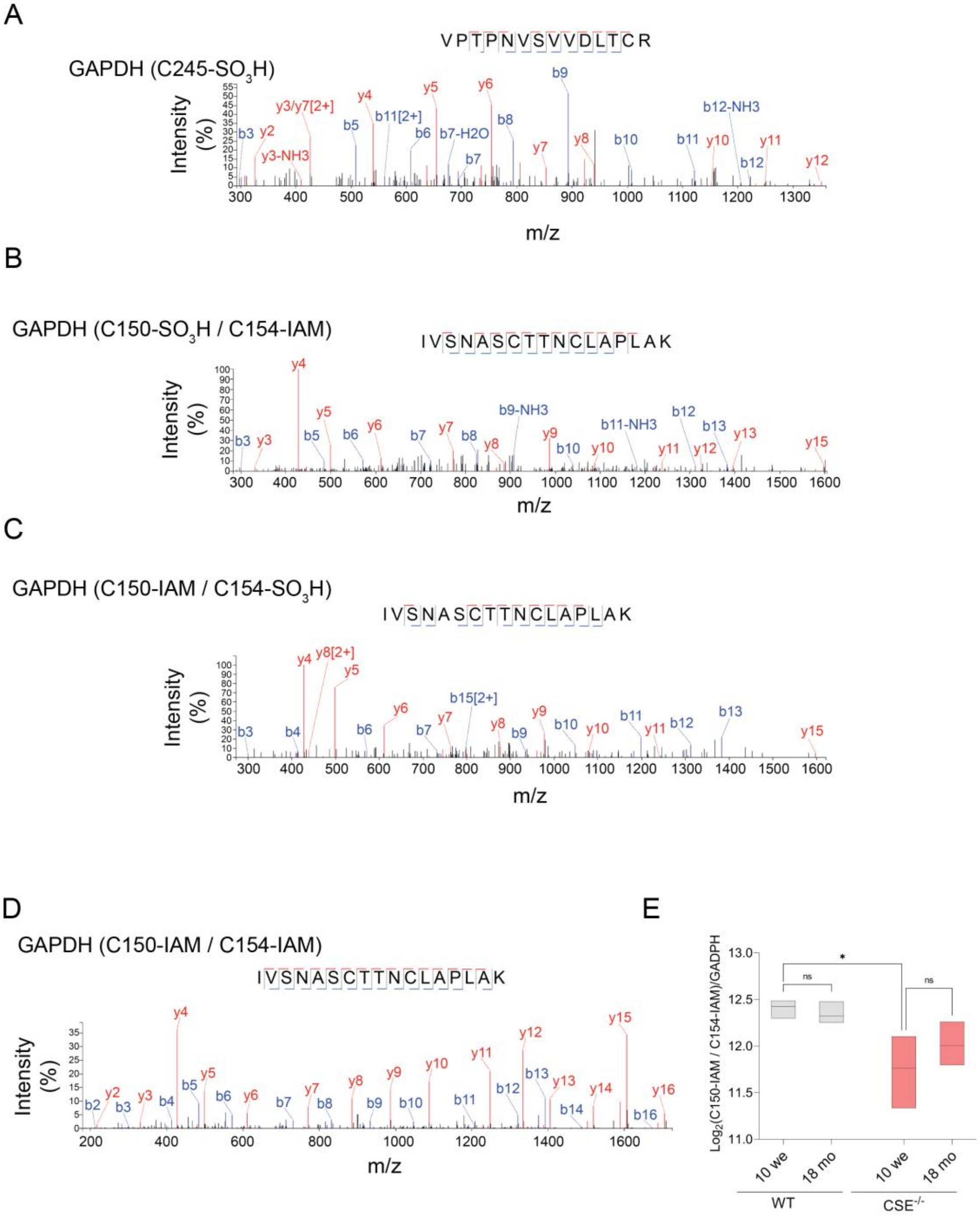


**Supporting Figure S23. Detection of GAPDH cysteine modifications in aged wild type (WT) and CSE$^{-/-}$ brain samples.** (**A**) Annotated TimsTOF CID MS/MS spectrum displaying the sulfonylated C245 residue in GAPDH. (**B**-**C**) Annotated TimsTOF peptide spectra depicting the sulfonylated C150 (**B**) and C154 (**C**) residues in GAPDH. Quantitative evaluation of these peptides was challenging due to their limited abundance. (**D**) Annotated TimsTOF CID MS/MS peptide spectra illustrating the double carbamidomethylated C150/C154 peptide. (**E**) Quantification of the fully reduced C150/C154 peptide, as depicted in (**D**). The quantification data demonstrates a noteworthy decrease of the fully reduced C150/C154 peptide levels in 10-week-old CSE$^{-/-}$ mice when compared to their WT counterparts (2-way ANOVA, Tukey's HSD Test for multiple comparisons; *<0.05).

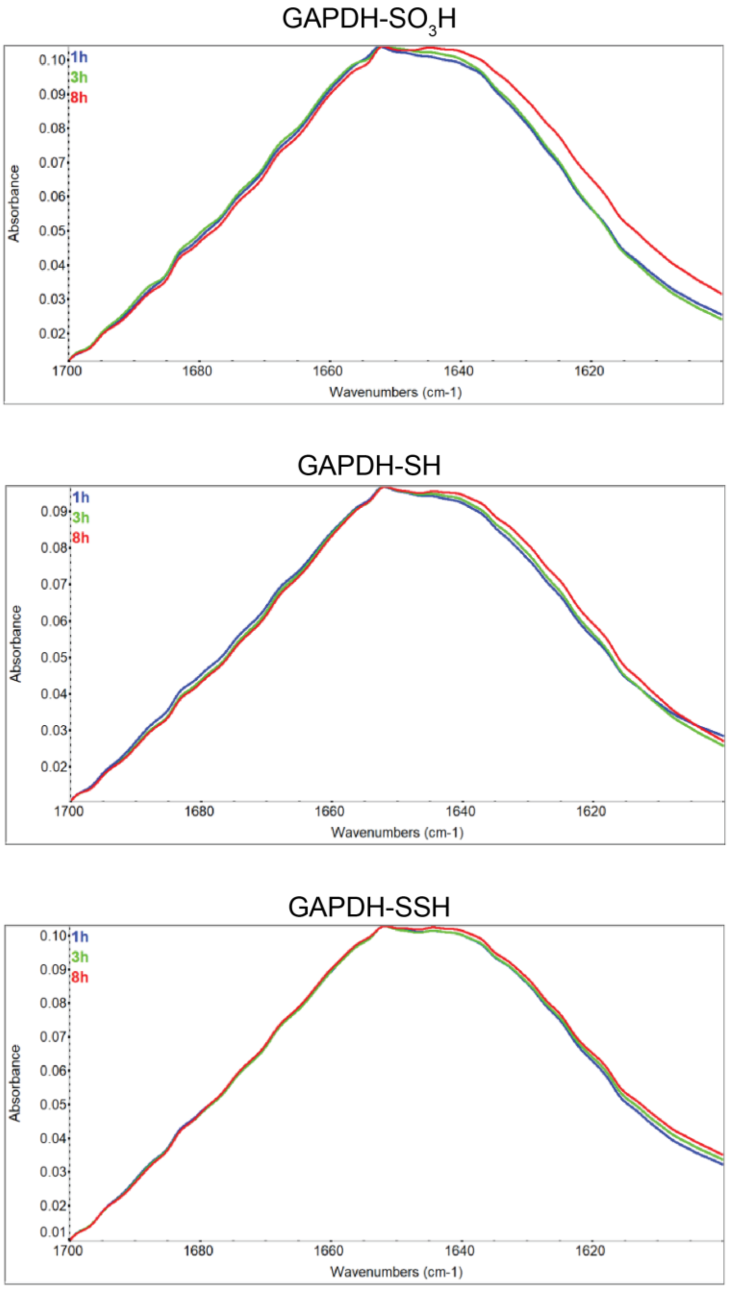


**Supporting Figure S24. Structural changes of GAPDH induced by sulfonylation, monitored within amide I region of FT-IR spectra.** Spectra of samples containing either sulfonylated, reduced, or persulfidated GAPDH (30 µM) were analyzed in OMNIC software. Amide I region arises mainly from C = O stretching vibrations thus is very sensitive to changes in secondary structures. Between 1600–1700 $cm^{-1}$ each secondary structure contributes to the absorption in a certain wavenumber range. The most prominent change detected in oxidized GAPDH after 8h was signal increase in the regions attributed to beta-sheets in general, namely 1616–1625 $cm^{-1}$ assigned to aggregation-prone inter/intramolecular beta-sheet, 1626–1640 $cm^{-1}$ assigned to intramolecular beta-sheet.

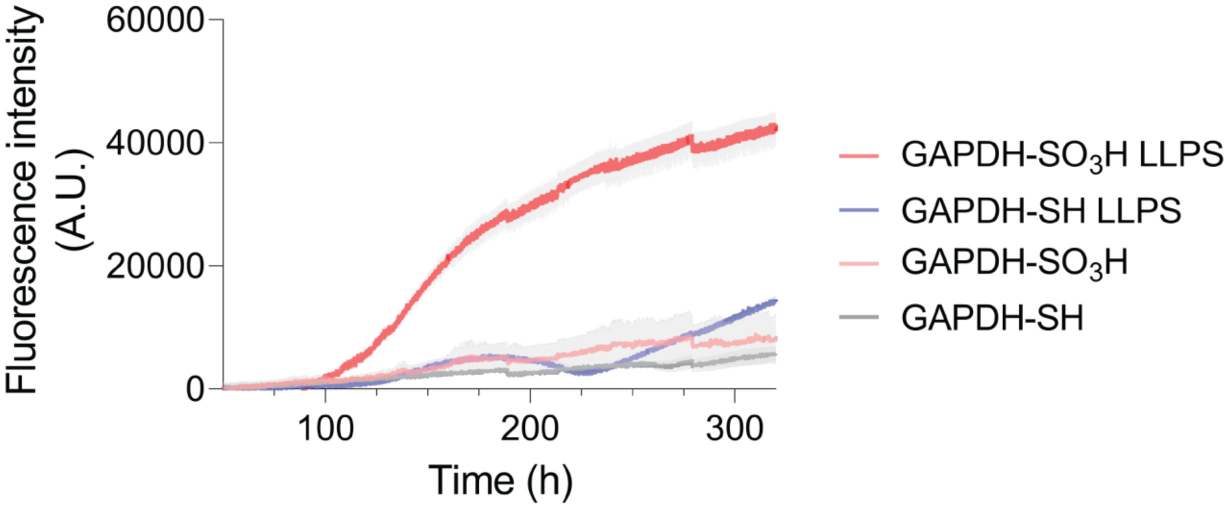


**Supporting Figure S25. LLPS of sulfonylated GAPDH promotes its aggregation.** The aggregation tendencies of sulfonyalted GAPDH were assessed using the PROTEOSTAT® assay. Solutions containing either reduced or sulfonylated GAPDH (10 µM) were combined with 500 µM DTT to keep cysteine under the reduced form. This combination was carried out in two conditions: (i) in the presence of total RNA extract, to simulate liquid-liquid phase separation (LLPS), and (ii) in the absence of total extract RNA, free of LLPS. The outcomes demonstrate that over a period of 100 hours, sulfonyalted GAPDH within the LLPS environment initiates an aggregation process, in stark contrast to conditions lacking LLPS where stability prevails.

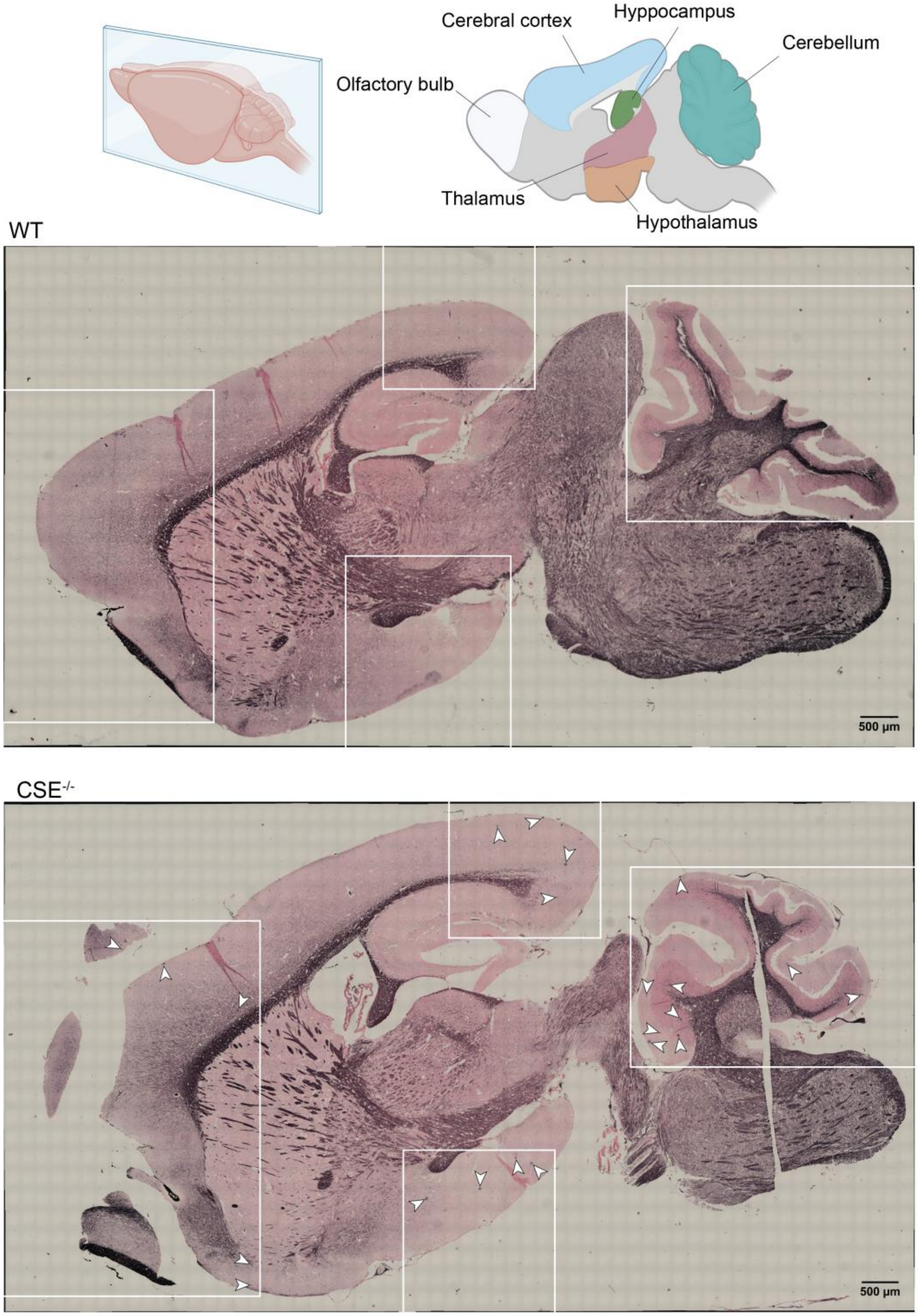


**Supporting Figure S26. Old CSE$^{-/-}$ mice exhibit multiple Gallyas silver–stained neuritic-like plaques.** Representative microscopy images of Gallyas staining of the sagittal brain section in 18-month-old WT and CSE$^{-/-}$ mice. Fibrillary tangles (white arrows) were present in different regions of the CSE$^{-/-}$ mice such as the hypothalamus, the cerebral cortex, and the cerebellum (indicated by the white boxes). WT mice did not show any detectable lesions
